# GABA_A_ receptor gating imaged on the millisecond timescale

**DOI:** 10.1101/2025.09.29.679341

**Authors:** Daniel B. Mihaylov, Tomas Malinauskas, Gisela D. Cymes, Jonas Miehling, Veronica T. Chang, Katerina Naydenova, Claudio Grosman, A. Radu Aricescu

**Affiliations:** MRC Laboratory of Molecular Biology, Francis Crick Avenue, Cambridge, CB2 0QH, UK; Division of Structural Biology, Centre for Human Genetics, University of Oxford, Oxford, OX3 7BN, UK; Department of Molecular and Integrative Physiology, University of Illinois at Urbana-Champaign, Urbana, IL 61801, USA

## Abstract

Type-A γ-aminobutyric receptors (GABA_A_Rs) are fast pentameric ligand gated ion channels (pLGICs)^1–5^. Within a millisecond, saturating agonist concentrations trigger activity bursts consisting of high-frequency fluctuations between conductive and non-conductive states^6,7^. These can last for tens to hundreds of milliseconds until, stochastically, receptors adopt stable, long-lived, desensitised conformations^2,8,9^. This highly dynamic process, known as gating, controls transient passages of chloride ions across plasma membranes to enable neurotransmission and other fundamental processes in animal physiology^10–12^. The conformational transitions inside activity bursts, inferred from electrophysiology, have remained inaccessible to structural biology investigation. Here we describe the gating motions of three GABA_A_ receptor variants imaged within the first 10 milliseconds of agonist application by cryogenic electron microscopy (cryo-EM)^13^. We illustrate how activation and desensitisation proceed through multiple asymmetric states, supported by major secondary, tertiary and quaternary structural rearrangements, and demonstrate that the same gating principles apply to both homomeric and heteromeric GABA_A_Rs. Furthermore, we show that cholesterol and phospholipids stabilise newly formed inter-subunit interfaces and obstruct channel pores in short-lived desensitised states, while phosphatidylinositol 4,5-bisphosphate (PIP_2_) precludes the opening of both α1β3 and α1β3γ2 GABA_A_R channels. Our results provide a novel framework to interpret decades of electrophysiology observations and suggest a broadly applicable approach to investigate mechanistically the vast arrays of physiological and pharmacological modulators of GABA_A_Rs^5,14^ and other fast neurotransmitter receptors^15,16^. Moreover, the subunit interfaces and lipid-binding pockets that form and disappear during GABA_A_R gating provide new opportunities to discover modulators with improved specificity and therapeutic properties.

**One sentence summary:** Single-particle cryo-EM was used to explore the dynamic conformational landscape of three human GABA_A_ receptor variants within the first 10 milliseconds of interaction with their neurotransmitter agonists.

---

Vertebrate pLGICs are a large class of cell-surface receptors that mediate the fast signalling actions of neurotransmitters including γ-aminobutyric acid (GABA), acetylcholine, serotonin, glycine and histamine^8,16^. The term “fast” covers almost six orders of magnitude: from microseconds to seconds^2,17–19^. This range reflects the vast potential of pentameric assembly diversity, the impact of membrane lipids, post-translational modifications and a plethora of positive and negative allosteric modulators, natural and synthetic^5,20–22^. Neurotransmitter agonists bind to orthosteric pockets formed between extracellular domain (ECD) interfaces, under so-called loops C, a feature structurally conserved across all pLGIC families. However, the molecular “gates” that control the flow of ions across cellular membranes are located in the transmembrane domain (TMD) region, approximately 50-60 Å away from these sites^9,23–28^. The current wisdom is that pLGICs obey a classic “Monod-Wyman-Changeux” (MWC) gating mechanism whereby receptors oscillate between closed and open states through cooperative and symmetric quaternary structure changes, which can be preferentially stabilised by small molecule ligands^16,29–31^. Structural results, so far, have been consistent with this model. Upon agonist binding, loops C are stabilised in “closed” conformations that “cap” the orthosteric sites as a result of inwards ECD tilting, counter- clockwise rotation and overall compaction, a mechanism described as “unblooming”^29^ or “lock and pull”^26^. These conformational changes are transmitted to the TMD resulting in a transient, concerted and (pseudo)symmetric opening of the activation gate while the desensitisation gate closes^26–28^. However, with a single exception, all pLGIC structures reported to date have been solved at equilibrium and did not sample time windows that are meaningful for the physiological gating process. The exception comes from classic work in tubular crystals grown from the *Torpedo marmorata* electric organ membranes, where an open nAChR conformation has been observed upon brief (less than 5 ms) agonist incubation^32,33^. However, such reconstructions are limited in resolution, and most ion channels do not form tubular lattice arrangements. As a result, the pLGIC gating mechanism remains unknown owing to the lack of structural information beyond highly stable, equilibrium snapshots.

Electrophysiology studies, with their high temporal resolution, suggest that the pLGIC gating process is considerably more complex. For example, GABA_A_ receptors undergo millisecond-scale transitions among conductive and non-conductive states^2,6,7^ that could be driven by independent movements of individual subunits^34^. Molecular dynamics simulations of glycine receptors suggest that a degree of activation gate asymmetry might be essential for channel opening^35^. Intriguingly, asymmetric structures at equilibrium have occasionally been observed^36–39^ but their significance remains unclear. In extreme cases, such asymmetric conformations are, unambiguously, sample preparation artefacts^40–43^, which highlights the challenges and potential pitfalls of studying pLGICs in non-native environments.

Here we report how three prototypical human GABA_A_R variants, reconstituted in brain-lipid nanodiscs, gate on a physiologically meaningful timescale.

## Gating mechanism of the homomeric GABA_A_R-β3

To capture the fast GABA_A_R gating motions, we used the plunge-freezer instrument developed for the pioneering *Torpedo* nAChR investigations^13^. Samples were applied onto EM grids positioned 35 mm above a cryo-holder filled with liquid ethane. An atomizing sprayer was triggered to deposit ligand droplets onto accelerating grids, 5 mm above the ethane, effectively limiting the ligand exposure to a maximum of 10 ms before vitrification (Supplementary Fig. 1 and Supplementary Discussion).

Using this approach, we first sought to explore the gating dynamics of the simplest GABA_A_R arrangement, a homopentamer assembled from β3 subunits (GABA_A_R-β3)^8^. This receptor adopts a five-fold symmetric desensitised conformation upon incubation with agonists including benzamidine^8^ and histamine (HSM)^44^. To boost protein yields, as required for the time-resolved setup, we replaced the M3-M4 loop with the SQPARAA linker, known to have minimal impacts on the kinetic properties of pLGICs^45–47^. To compare the behaviour of the full- length and truncated constructs, we applied 10 mM HSM to whole cells expressing each. Because the rate of solution exchange is <1 ms, the conditions mimic the rapid diffusion observed on the EM grid. Both constructs showed broadly similar responses, although the truncated β3 homomer displayed slightly lower rates of desensitisation and activation as well as a reduced peak amplitude (Supplementary Fig. 2). GABA_A_R-β3 was reconstituted in lipid MSP2N2 nanodiscs, offering a closer to physiological environment than other nanodiscs such as Saposin A (see Supplementary Discussion)^9,48^. Upon spraying HSM, we reconstructed six distinct receptor states with spatial resolutions between 2.5 Å and 3.5 Å. We present them in a hypothetical chronological order, guided by structural divergence from the unliganded, resting conformation and progressing towards the long-lived desensitised state, which is analogous to previously published agonist-bound structures solved at equilibrium^8,44^ (Fig. 1, Extended Data Fig. 1-4).

**Figure 1.**
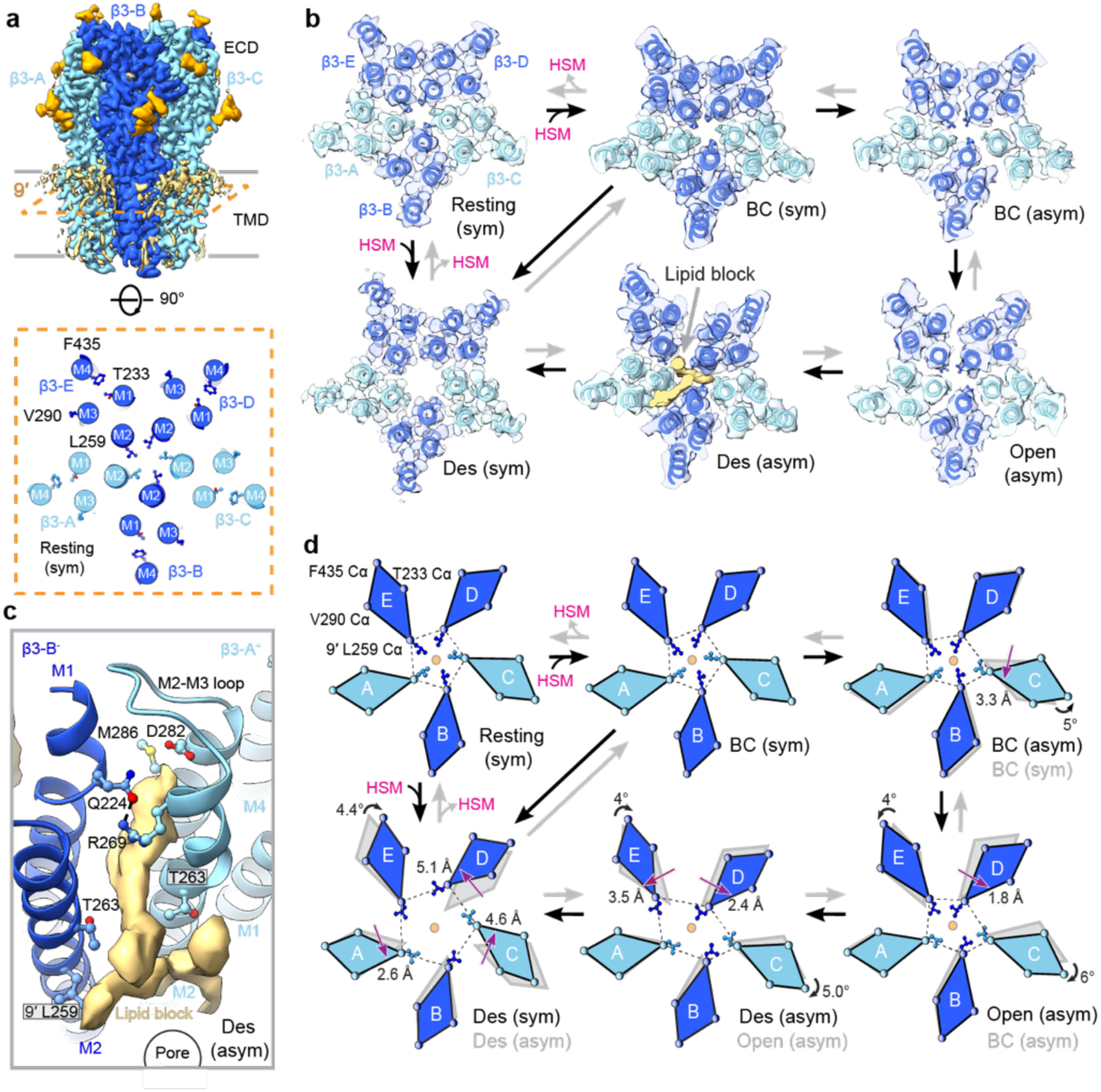
The GABA_A_R-β3 homomer gating mechanism. **a,** Cryo-EM map of the β3 homomer in a resting state, viewed parallel to the lipid bilayer plane (top) and a cross-section at the level of the 9′ activation gate showing the residues (Leu259, Thr233, Val290, Phe435) whose Cα atom positions were used to generate the tetragons in d). **b,** Cryo-EM maps of all states reconstructed from the dataset upon HSM exposure, viewed from the extracellular space and cross-sectioned at the level of the 9′ activation gate. Lipids are shown in khaki. The six states are labelled as follows: Resting, bound-closed symmetric – BC (sym), bound-closed asymmetric – BC (asym), open asymmetric – Open (asym), desensitised asymmetric – Des (asym) and desensitised symmetric – Des (sym). **c,** Close-up inside the channel pore in the Des (asym) state, which is blocked by a lipid, coloured in khaki. The amino acid side chains lining the binding site are shown in a ball-and-stick representation. Dashed lines indicate putative hydrogen bonds. **d,** Schematic representation of TMD motions where subunits are shown as tetragons. Distances (depicted with a straight purple arrow) represent the Leu259 Cα displacement from the previous position (in grey) to the current one (coloured). Rotation angles (shown as curved black arrows) about the central pore axis (depicted as a transparent orange circle) illustrate the displacement of Phe435 Cα atoms from the previous state to the current.

The resting state is symmetric, and no agonist molecules are present in the orthosteric pockets. The GABA_A_R-β3 TMD region, in addition to the “classic” pLGIC activation and desensitisation gates formed by pore-lining residues at the 9’ and -2’ M2 helix levels, respectively, features an additional gate at the 17’ level which can be controlled by pH changes^49^. The gate is formed by a ring of β3His267 side chains previously shown to coordinate Zn^2+^ ions which in turn inhibits α1β3 GABA_A_ heteromers (Extended Data Fig. 3)^7,50^. In this state, all three gates are closed, i.e. cannot allow the passage of fully dehydrated chloride ions, whose Pauling radii are approximately 1.8 Å^51^ (Fig. 1, Extended Data Fig. 5a).

Upon HSM binding, the five orthosteric sites become capped through closure of all five loops C and the symmetric ‘unblooming’ of the ECD (Supplementary Video 1; Extended Data Fig. 3b). Compared with the heteromeric synaptic α1β3γ2 GABA_A_R^26^, GABA_A_R-β3 exhibits a less pronounced inward tilt, anticlockwise ECD rotation, and loop C closure. As the receptor progresses through its gating cycle, the ECD remains largely structurally unchanged (Supplementary Figs. 3-4). Within this trajectory, the second and most populated state features five HSM molecules bound but retains a resting-like TMD in which the 17′, 9′, and –2′ gates remain shut (Fig. 1b,d; Extended Data Figs. 2c, 3c, 5a). The predominance of this shut conformation indicates that HSM acts as a weak agonist. To test this, we applied HSM alone or together with the positive modulator etomidate (ETM) to cells expressing GABA_A_R-β3. With HSM and ETM, activation and desensitisation rates were faster and the peak amplitude greater, confirming that HSM alone fails to activate the channel as effectively as the combination (Supplementary Fig. 5). We designate this conformation as the bound-closed (BC) symmetric state and will discuss its significance later.

The TMD symmetry breaks in the third state observed, which displays an outwards translation and anticlockwise rotation of the transmembrane helical bundle in the subunit we label, for convenience, β3-C (Fig. 1b, d, Extended Data Fig. 3c and 4d). The process can be initiated stochastically by any subunit, owing to the homomeric nature of the receptor. This conformational change is accompanied by an outwards clockwise displacement of the β3-B M2-M3 loop and is stabilised by a hydrogen bond network involving β3Glu270-B, β3Thr271, β3Lys274-B and β3His267-C (Extended Data Fig. 4d). The channel pore remains non-conductive, with all 3 gates shut (Extended data Fig 5a), therefore we refer to this state as bound-closed (BC) asymmetric.

The fourth state is also asymmetric and represents an open GABA_A_ receptor. To accurately describe the pore geometry of the asymmetric open state, we used PoreAnalyser, which reports both semi-minor and semi-major radii of asymmetric pores and estimates channel conductance^52^. The narrowest region of the pore measures 2.2 Å – a value between the 1.8 Å radius of a dehydrated Cl⁻ ion and the 3.2 Å radius of a fully hydrated one^51,53^ - suggesting that only partially hydrated Cl⁻ ions can pass (Supplementary Fig. 6). This agrees with the ∼2.8 Å average pore radius measured in neurons, which reflects the combined contributions of various open GABA_A_ receptor assemblies^54^. Partial hydration is likely stabilised by interactions with hydrophilic pore-lining residues, many of which are epilepsy mutants, underscoring their functional relevance (Supplementary Fig. 7). The predicted conductance of 27 pS for the GABA_A_R-β3 open state matches the reported conductance of the β1 homomeric homologue^55^. Pore opening relies on the rotation and translation of both β3-A and β3-E subunits, which further accentuates the TMD asymmetry leaving only two β3His267 side chains oriented towards the pore (Fig. 1b,d, Extended Data Fig. 3c). The open pore conformation is stabilised by new hydrogen bond networks between M2 β3-C^+^ and M1 β3-D^-^, as well as between M2 β3-D^+^ and M1 β3-E^-^. Additional reinforcement involves interactions between the M2-M3 loop of β3-D and the β9 loop of β3-E (Extended Data Fig. 4e).

The open state is followed by another asymmetric conformation that we term “short-lived desensitised”. Here, the β3-C TMD moves further out of the pentameric bundle and the β3-E TMD is drawn closer to β3-A. As a result, the channel pore is primarily delineated by the M2 helices of the other four subunits (Fig. 1b,d, Extended Data Fig. 3c and 5f). The asymmetric desensitised state could have been classified as open based solely on the pore profile (Extended Data Fig. 5a). However, the weaker β3-D^+^/β3-E^-^ TMD interface has opened a path for lipids to “invade” the pore and block it (Fig. 1b-c). The high degree of asymmetry at the TMD level is transmitted to the ECD, where loop C at the β3-D^+^/β3-E^-^ interface opens to release the agonist leaving only four HSM molecules bound. In contrast, all other states apart from resting are fully liganded at all five orthosteric sites (Extended Data Fig. 3b, Supplementary Figs. 3-4). The TMD regions of β3-A and β3-B subunits are already in conformations that resemble those previously reported for the long-lived desensitised state^8,44^, which is finally reached upon substantial rearrangements of β3-C, β3-D and β3-E (Fig. 1d, Extended Data Fig. 4g, Supplementary Video 1).

It is important to emphasise that all asymmetric states described here were only be observed during millisecond ligand-receptor interactions and that synthetic fiducials (such as nano/megabodies or Fab fragments) were not used. Control experiments with the same construct either sprayed with buffer alone (see Supplementary discussion) or incubated with HSM on longer timescales^44,56^, led to symmetric states exclusively, either resting or long-lived desensitised. The latter is indistinguishable from those we solved in the presence of megabody, Mb25^44,56,57^. Our results suggest that the asymmetric states represent short-lived intermediates in the gating cycle of the receptor. Unlike the detergent-damaged asymmetric GABA_A_ receptor conformations reported previously^40–42^, the ECDs and TMDs for all subunits seen in the short- lived states remain structurally coupled (Supplementary table 1).

### PIP_2_ restricts the opening of GABA_A_R heteromers

To determine whether the homomeric β3 gating mechanism is conserved across GABA_A_ receptor subtypes, we decided to investigate next the full-length α1β3 diheteromeric receptor. We sprayed a mixture of GABA and HSM, the latter acting as a positive allosteric modulator^7^ of this receptor type. Surprisingly, we could only separate three states in this dataset, all symmetric: resting, bound-closed and desensitised (Extended Data Fig. 6a-c). The α1β3 GABA_A_R is extrasynaptic and displays a lower open probability than synaptic variants even at saturating agonist concentrations^50^. We therefore tested the prototypical synaptic tri- heteromeric α1β3γ2L variant, again full-length, with a saturating concentration of GABA and supplemented with the general anaesthetic ETM. This combination of molecules is known to elicit a maximum GABA_A_R open probability and activate virtually all channels in a population^58,59^. Moreover, α1β3γ2L was reconstituted in nanodiscs with ∼40% cholesterol to mimic the synaptic environment as accurately as possible^22,60,61^ (see Supplementary discussion). Despite all these attempts, only three states, all symmetric, were present in this dataset: bound-closed, desensitised with only GABA bound and desensitised with both GABA and ETM bound, none of which were conductive (Extended Data Fig. 6d-f).

Therefore, it became evident that, unlike the β3 homomer, di- and triheteromeric receptors that contain α1 subunits cannot undergo activation in our reconstitution system. Instead, they appear to transition straight from the bound-closed symmetric state to the long-lived symmetric desensitised state without opening even on the millisecond timescale. To determine the mechanism of inhibition, we purified the full-length α1β3γ2L receptor in SMALPs, which directly solubilise membrane proteins in their endogenous lipid environment^62^. However, upon incubation with GABA, the TMD region became distorted and the ECD region appeared uncoupled from the TMD, precluding further investigations of the gating mechanism in SMALPs. Nevertheless, we were able to reconstruct the resting state with well coupled ECD and TMD regions. Importantly, we found this conformation to be identical to the resting state we have observed in brain lipid nanodiscs (see Supplementary Discussion).

Unexpectedly, 3D classification of the resting α1β3γ2L receptor in SMALPs revealed four distinct classes with different occupancy of the PIP_2_ binding sites^9^. Most particles have a PIP_2_ molecule bound to each α1 subunit. We could also separate classes with either of the two PIP_2_ sites occupied, as well as receptors without PIP_2_ bound (Fig. 2a-b). PIP_2_ binds to the α1 subunit on the intracellular side of the TMD and its inositol head group stabilises the membrane-proximal segments of the α1 M3-M4 loop in the intracellular domain (ICD). Further interactions with the M3 extensions from neighbouring β3 subunits, and with the side-chain of M2 α1Arg249 residues, interlock the β3-α1 subunits (Fig 2c). As a result, at least when both PIP_2_ sites are occupied, the -2′ desensitisation gate cannot expand, and the channel pore cannot open in response to agonist binding. Our results suggest that (*i*) Currents recorded by electrophysiology are likely generated by receptors that have either one, or most probably no, PIP_2_ molecules bound; (*ii*) Between a third and three quarters of receptors expressed at the cell surface, at least in recombinant systems, can only adopt non-conductive conformations, either resting or long-lived desensitised; (*iii*) These observations will apply to all receptor arrangements that contain α1/2/3/5 subunits, all of which harbour PIP_2_ binding sites^9^.

**Figure 2.**
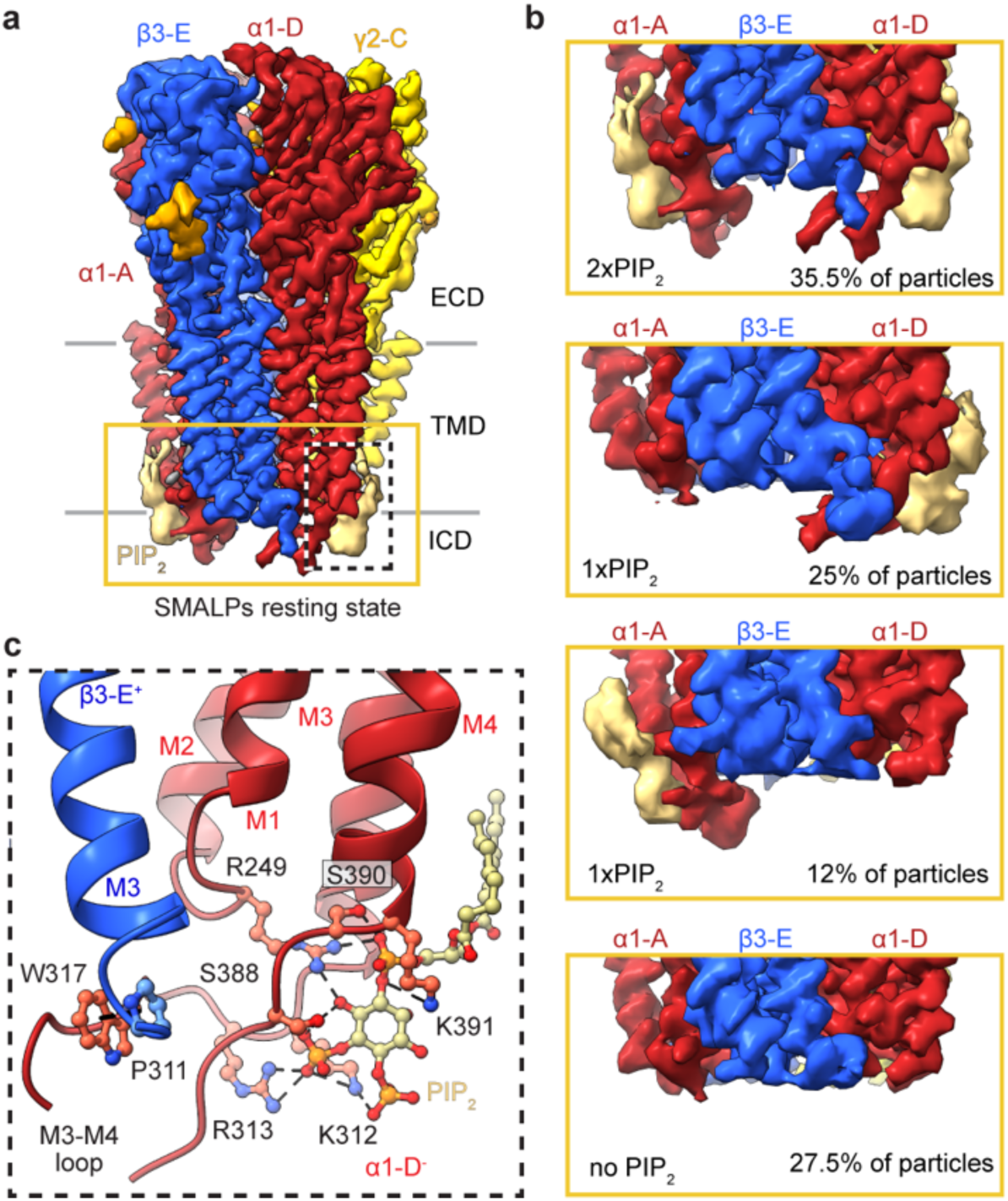
Variable PIP_2_ occupancy in the full-length GABA_A_R-α1β3γ2 isolated directly from plasma membranes. **a,** Cryo-EM map of the full-length α1β3γ2 in SMALPs 30100 viewed parallel to the membrane. **b,** The variable PIP_2_ occupancy in the native membrane environment. Two PIP_2_ molecules are bound to the receptor in 35.5% of the particles. One PIP_2_ molecule is bound to either of the α1 subunits in 37% of the particles. 27.5% of the particles have no PIP_2_ bound. **c,** The PIP_2_ binding pose. The amino acid side chains lining the binding site are shown as ball and sticks. Dashed lines indicate putative hydrogen bonds and π–π interactions.

### The GABA_A_R-α1β3 gates via two branches

To remove the inhibitory action of PIP_2_, we truncated the α1 M3-M4 intracellular loops as described above for β3 subunits and shown previously to abolish PIP_2_ binding^7^.

Upon application of GABA, but not buffer alone, we resolved nine distinct states with resolutions ranging from 3.4 Å to 2.5 Å (Extended Data Fig. 1, 7, Supplementary discussion). Unambiguous densities for the agonist could be observed at the orthosteric β3^+^/α1^-^ ECD sites, consistent with the binding mode previously reported (Extended Data Fig. 8)^7,26^. As in the β3 homomer, the α1β3 activation cycle begins with GABA binding, which shifts the receptor from a quasi-symmetric resting state to a bound-closed (BC) state. This transition involves loop C closure together with ECD ‘unblooming’, anti-clockwise rotation, and tilting, while the channel pore remains shut and symmetric. Compared with the β3 homomer, the α1β3 diheteromeric GABA_A_R undergoes larger ECD motions, though still less pronounced than in the synaptic α1β3γ2 receptor (Supplementary Figs. 8-9). Whereas GABA_A_R-β3 progresses along a linear gating path, α1β3 receptors diverge with equal probability into two alternative branches (Fig. 3; Extended Data Fig. 8c–d and 9). Eventually, both paths converge into the same symmetric long-lived desensitised state.

**Figure 3.**
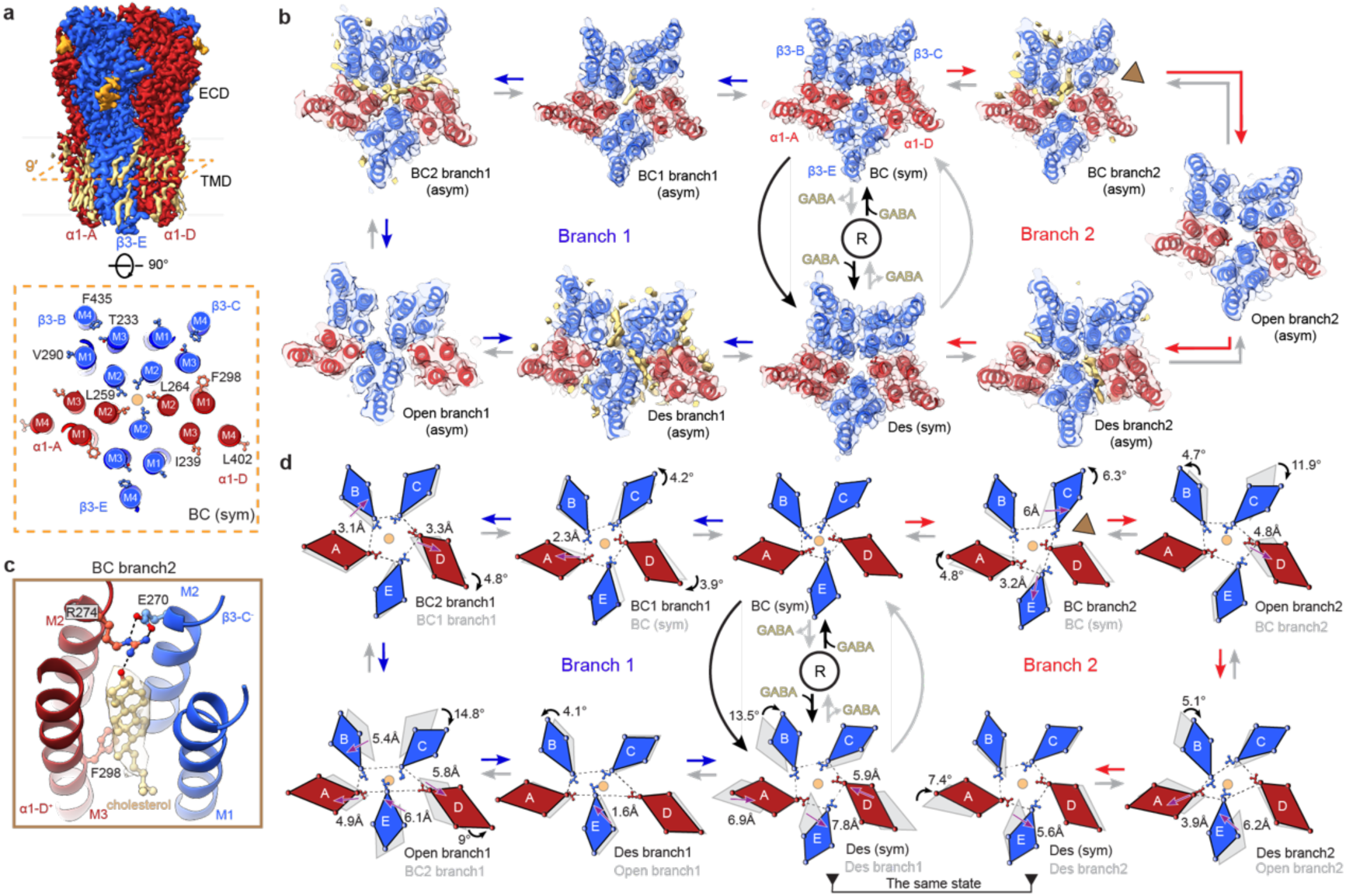
The GABA_A_R-α1β3 diheteromers gate via two branches. **a,** Cryo-EM map of the α1β3 diheteromer viewed parallel to the membrane plane (top) and a cross-section at the level of the 9′ activation gate showing the residues (α1 – Leu264, Ile239, Phe298, Leu402; β3 **-** Leu259, Thr233, Val290, Phe435) used for the generation of the tetragons in d). **b,** Cryo-EM maps of all states classified from the dataset upon millisecond-scale GABA exposure viewed from the extracellular space and cross-sectioned at the level of the 9′ activation gate. α1β3 gates via two branches – branch 1 (left) and branch 2 (right) emerging from the same resting state (labelled as R in the figure) and bound-closed (sym) state and converging in the same desensitised symmetric state. Lipids are shown in khaki. The acronyms of the states are the same as in Fig. 1. **c,** View of the cholesterol (carbons khaki, oxygens red) binding site in BC (asym) of branch 2. The amino acid side chains lining the binding site are shown as ball and sticks. Dashed lines indicate hydrogen bonds. **d,** Representation of the motions with the subunits shown as tetragons. The distances (depicted with a straight purple arrow) were measured based on the movement undergone by the M2 Leu259 Cα atoms from the current state (coloured) compared to the previous (in grey). Rotations (shown as curved black arrows) were measured between the centre of the resting state (depicted as a transparent orange circle) and the M4 Cα atoms of the current state compared to the previous.

In the first branch of the gating pathway, activation is initiated with the destabilisation of the α1-D^+^/β3-C^-^ TMD interface, allowing β3-C to rotate away from the pore and form a hydrogen bond network with β3-B (Fig. 3d, Extended Data Fig. 9a-b). As branch 1 transitions to bound-closed 2 branch 1 (asym), β3-B undergoes an outward drift, while α1-D rotates further counter-clockwise, intensifying the asymmetry of the pore (Fig. 3b,d). This is followed by a shift to open branch 1 (asym) that involves the collective translation and rotation of β3-B, α1- A, and β3-E as a bundle while β3-C experiences a substantial counter-clockwise rotation of 14.5°. The inter-subunit space of this state is wide enough to allow lipids to protrude the pore, however, no clear lipid density can be observed suggesting that if such block can occur it is only transient, and the state can be labelled as conducting (Fig. 3b, Extended Data Fig. 4b, Supplementary Fig. 10). The pore of open branch 1 is lined with hydrophilic residues, has a radius of 3.8 Å allowing a fully hydrated Cl^-^ ion through and has an estimated conductance of 40.1 pS. For reference, the conductance reported experimentally is 19 pS^7^ (Supplementary Fig. 8-9). Such high degree of asymmetry has been proposed previously for the open state of α1β3 receptors *in vivo*^63^. In desensitised branch 1 (asym), β3-B rotates counter-clockwise, and β3-E translates away from α1-D, excluding it from the pore and allowing lipids to invade the pore and block the passage of ions. During the transition to the symmetric long-lived desensitised state, chains α1-A, β3-B, α1-D, and β3-E undergo significant conformational changes, whereas chain β3-C is already in a state close to that in the symmetric desensitised (Fig. 3b,d, Supplementary Video 2).

In contrast to branch 1, branch 2 enters the asymmetric cycle via GABA binding to the β3-B^+^/α1^-^A- interface allowing β3-C to rotate towards α1-D in bound-closed branch 2 (asym) (Fig. 3b,d, Extended Data Fig. 9c). The newly formed interaction between β3-C and α1-D is stabilised by the critical insertion of cholesterol at the interface which makes a hydrogen bond with α1Arg274-D (Fig. 3c, Extended Data Fig. 9c-d). Cholesterol has been shown to be pivotal for the gating of several pLGICs, including GABAAR and for the coupling between the ECD and TMD^22,64,65^. During opening, α1-D moves away from the pore, while β3-C undergoes a substantial clockwise rotation in open branch 2 (asym) (Fig. 3b,d). The open state has well insulated inter-subunit space thereby precluding lipids from blocking the pore (Extended Data Fig. 4b, Supplementary Fig. 10). It has a radius of 3.4 Å allowing a fully hydrated Cl- ion to pass and an estimated conductance of 38.4 pS (Supplementary Fig. 6). In desensitised branch 2 (asym), β3-E drifts significantly towards the centre of the pore, alongside β3-B. This results in the exclusion of α1-A from the pore which is now formed by the remaining four subunits. In branch 1 the other α1-D is displaced. Throughout the desensitisation of branch 2, β3-E translates, while α1-A rotates counter-clockwise. The other three subunits are already close to the conformation they attain in the desensitised state and therefore do not undergo substantial conformational changes (Supplementary Video 3).

### Gating mechanism of the α1β3γ2 triheteromer

After confirming that PIP_2_ inhibits the opening of full-length α1-containing receptors in our nanodiscs, we investigated the impact of PIP_2_ removal through subunit truncation on the synaptic α1β3γ2 triheteromer, which is the predominant receptor type in the hippocampus^66^. Triheteromeric α1β1-3γ2 receptors with truncated subunits maintained signalling profiles similar to WT receptors^40–42^.

Upon millisecond-scale application of GABA to the α1β3γ2 receptor reconstituted in MSP2N2 nanodiscs with ∼40% cholesterol, we could observe seven distinct states (Extended Data Fig. 10). Like the β3 homomer and the α1β3 diheteromer, the activation cycle begins with the symmetric resting state. This is followed by the symmetric bound-closed state, where the rotation of loop C, ECD tilt and ECD ‘unblooming’ are substantially more pronounced than for the β3 homomer and α1β3 diheteromer (Supplementary Figs. 11-12). The receptor then transitions into a series of intermediate asymmetric states (Fig. 4b).

**Figure 4.**
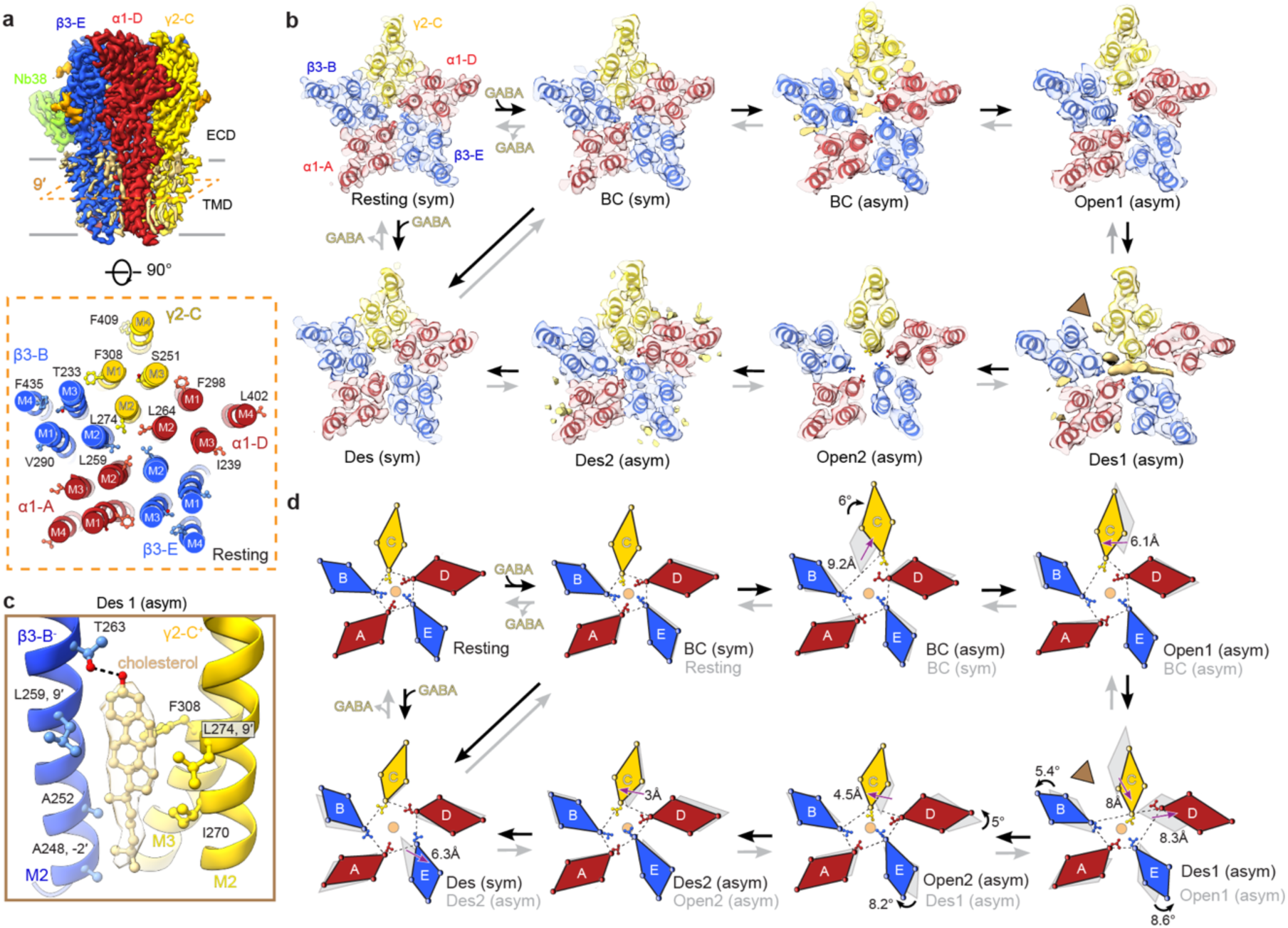
The gating of motions of triheteromeric α1β3γ2 receptors. **a,** Cryo-EM map of the resting α1β3γ2 triheteromer viewed parallel to the membrane plane (top) and a cross-section at the level of the 9′ activation gate showing the residues (α1 – Leu264, Ile239, Phe298, Leu402; β3 **-** Leu259, Thr233, Val290, Phe435; γ2 – Leu274, Phe308, Ser251, Phe409) used for the generation of the tetragons in d). **b,** Cryo-EM maps of all states classified from the dataset upon millisecond-scale GABA exposure viewed from the extracellular space and cross-sectioned at the level of the 9′ activation gate. The resting state was obtained from a separate dataset. The acronyms of the states are the same as in Fig. 1 **c,** View of the cholesterol (carbons khaki, oxygens red) binding site in Des1 (asym). The amino acid side chains lining the binding site are shown as ball and sticks. Dashed lines indicate hydrogen bonds. **d,** Representation of the motions with the subunits shown as tetragons. The distances (depicted with a straight purple arrow) were measured based on the movement undergone by the M2 Leu259 Cα atoms from the current state (coloured) compared to the previous (in grey). Rotations (shown as curved black arrows) were measured between the centre of the resting state (depicted as a transparent orange circle) and the M4 Cα atoms of the current state compared to the previous.

Unlike α1β3, which gates via two branches, the α1β3γ2 receptor is activated through a single gateway (Fig. 4). During the transition to the asymmetric bound-closed state, the γ2 subunit rotates towards the neighbouring α1 stabilised by an unidentified lipid as well as a hydrogen bond interaction between α1Tyr294 and γ2Thr278 (Fig. 4b,d, Extended Data Fig. 12a-b). Only one GABA molecule can be observed in the ECD at the β3-B^+^/α1-A^-^ interface of bound-closed (asym) (Extended Data Fig. 11b). This suggests that agonist binding to the interface at the clockwise position with respect to γ2 as viewed from the ECD, triggers an inter-subunit conformational wave. This wave pulls the β3-B^+^/α1-A^-^ subunits counter-clockwise and, in turn, pushes the γ2-C in the clockwise direction. The reason for the single gateway likely stems from the fact that the γ2^+^/β3^-^ TMD interface is less stable than between the α1^+^ and γ2^-^ subunits. Therefore, the free energy penalty for disrupting it is lower (Supplementary Table 1). As the cycle progresses from bound-closed (asym) to open 1 (asym), the γ2 subunit moves, re-establishing the TMD interaction with β3-B through a hydrogen bond network involving γ2Asp297-C, γ2Arg284-C, and β3Tyr220-B, along with γ2Ser291 and β3Pro184 (Fig. 4b,d, Extended Data Fig. 12c-d). The open state has a pore radius of 3.4 Å, permitting the passage of fully hydrated Cl⁻ ions, with an estimated conductance of 40.2 pS. The experimentally measured conductance is 27 pS^7^ (Supplementary Fig. 6). Subsequently, the receptor enters the short-lived desensitised 1 (asym), where the pore is blocked by lipids. During this phase, β3-B and β3-E rotate counter-clockwise while γ2-C and α1-D translate together (Fig. 4d). Stabilisation of this state is notably facilitated by a hydrogen bond network encompassing α1Asp287-D, α1Arg274-D, α1Lys279-D, γ2Asn239-C, and γ2Asn200-C (Extended Data Fig. 12e-f). We detected cholesterol bound at the γ2-C^+^/β3-B^-^ interface (Fig. 4c), consistent with its proposed role in GABA_A_ receptor gating^22^.

Moving into open 2 (asym), γ2-C translates whereas α1-D and β3-E rotate (Fig. 4b,d). This state is stabilised through a hydrogen bond network involving α1S270-D, α1T230-D, and β3Arg269-E (Extended Data Fig. 12g-h). The narrowest pore radius in the open 2 (asymmetric) conformation is 2.8 Å and its calculated conductance is 32.9 pS, consistent with the experimentally determined value of 27 pS (Supplementary Fig. 6). The pore is wide enough to allowing the passage of partially hydrated Cl^-^. The loss of hydration could be compensated by the hydrophilic residues now facing the pore (Supplementary Fig. 7). Moreover, the number of open and closed/desensitised states we observe are in agreement with findings from single- channel electrophysiology^7^. After the second opening, the receptor enters the short-lived desensitised 2 (asym), primarily driven by the translation of γ2-C (Fig. 4b,d,). Notably, this asymmetric state is stabilised by POPG inserting at the α1-D^+^/γ2-C^-^ interface (Extended Data Fig. 12i-j). During the final stage of the cycle, the receptor progresses into the long-lived desensitised state governed by the translation of β3-E (Fig. 4b,d, Supplementary Video 4).

### Principles of GABA_A_R activation and desensitisation

In this study, we explored the structural gating dynamics of multiple GABA_A_ receptor assemblies using millisecond-scale ligand application. We determined a common gating mechanism despite differences in subunit composition. Our proposed activation pathway is derived from structural similarities between equilibrium states (resting and long-lived desensitised) and the asymmetric states observed on the millisecond timescale. These intermediate states, captured by single particle, time-resolved cryo-EM, can be designated as closed, open, or desensitised. However, given the non-equilibrium conditions where all states coexist, the precise temporal ordering of these transitions remains inferential rather than definitive (Fig. 5). The mechanism we propose can be summarised in five steps as follows:

**Figure 5.**
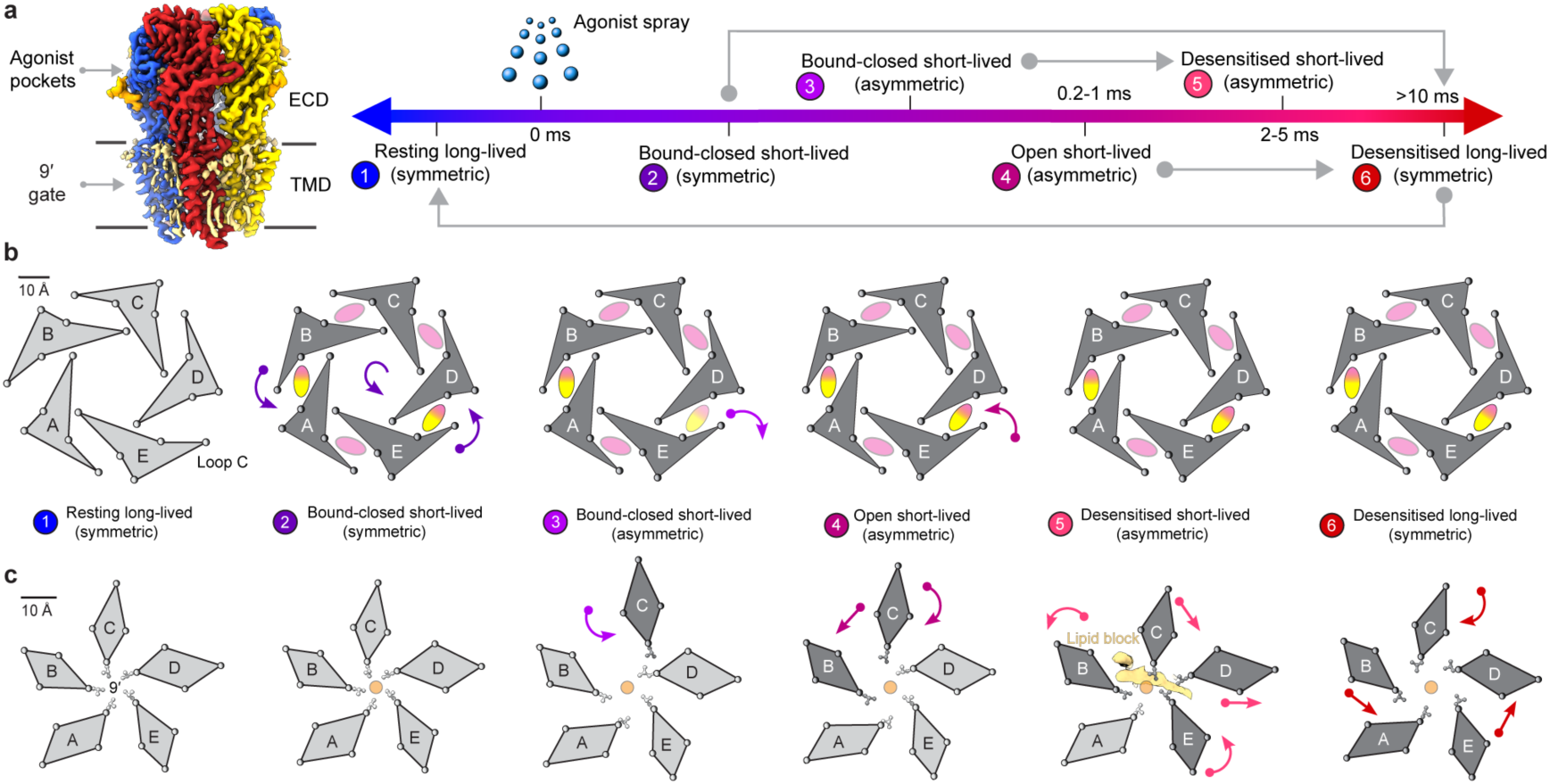
Principles of GABA_A_ receptor activation and desensitisation. **a,** Cryo-EM map of the α1β3γ2 triheteromer viewed parallel to the membrane plane, illustrating the location of the agonist binding pockets and the 9′ activation gate (left). A timeline on the right depicts the states involved in the receptor gating cycle. Grey lines represent possible bypasses of the otherwise stepwise transitions. **b,** A cross-section at the agonist-binding pocket level. The receptor structure is depicted, with individual subunits labelled A to E. In β3 homomers, all subunits are β3. In α1β3 heteromers, subunits A and D are α1, whereas the remaining subunits are β3. In α1β3γ2 triheteromer, subunits A and D are α1, B and E are β3, and C is γ2. Conformational changes and ligand binding/unbinding events in each state are indicated by arrowheads, with ligands represented as ellipses. The agonists that can be both HSM or GABA depending on the different receptor assemblies are depicted as a mixture of both GABA (yellow) and HSM (magenta). Regions of the receptor undergoing significant motion when compared to the resting state are highlighted in darker grey. **c,** A cross-section at the level of the 9′ activation gate. The spheres of the tetragons represent the Cα atoms of the M1, M2, M3 and M4 residues lying on the 9′ plane. The centre of the resting state is marked by an orange circle. Curved arrowheads represent rotational movements measured between the resting state centroid and the subunits, whereas straight arrowheads indicate translations between corresponding subunits across different states.

#### Step 1: Ligand binding and entry into the bound-closed state

Activation of the receptor begins with agonist binding. In the symmetric unliganded resting state, loop C is open, the ECD is in the low-affinity agonist state and both activation and desensitisation gates are closed. Upon agonist binding, loop C closes while the ECD tilts and ‘unblooms’. The bound-closed state, however, retains a resting-like TMD and is symmetric (Fig. 5b-c). It may seem tempting to identify this bound-closed conformation with the “primed” or “flip” states previously proposed to be visited as agonist-bound closed pLGICs open^67–69^. However, the low occupancy predicted for these intermediates on the basis of their estimated short mean lifetimes (tens of microseconds) contrasts with the high proportion of particles adopting this conformation. Functionally, BC (sym) likely corresponds to the intra-cluster shut state observed in single-channel recordings, which predominates when weak agonists are applied. Consistent with this, we show that HSM is a weak agonist for GABA_A_R-β3 (Supplementary Fig. 5), and structurally the bound-closed state is likewise the most populated (Extended Data Fig. 2). The bound-closed state appears to be a key branching point in the gating pathway: from this state the receptor either transitions into the bound-closed asymmetric states *en route* to opening or directly enters the symmetric, long-lived desensitised state without opening (Fig. 5a). Such a bifurcation has also been observed functionally in the case of nAChR, where only a small fraction of receptors open, while the majority rapidly enter the desensitised state, likely via the BC (sym) state^70,71^.

#### Step 2: Symmetry breaking and the formation of a pre-active intermediate

The second step of the gating cycle is the transition into the short-lived asymmetric states, where a single subunit (labelled as position C), which does not participate in ligand binding in heteromeric receptors, breaks the symmetry of the pore across all GABA_A_ receptor subtypes (Fig. 5c). A similar phenomenon has been observed for the *Torpedo* nAChR in tubular membranes^33^. The displacement of the first chain from the bundle is triggered by agonist binding to the adjacent ECD interface (Fig. 5b) and in the case of α1β3γ2 the state is mono- liganded. This weakens the interaction with the subunit in position C thus allowing it to rotate and disrupt the TMD interface. Although the GABA_A_ receptor can open rapidly within ∼160 μs^3^, the rise time for whole-cell activation, where 90% of receptors are in the open state, is approximately 4 ms^72^. In our non-equilibrium experiments, receptors were exposed to ligand for up to 10 ms, enabling the capture of transient intermediate states. Structurally, the asymmetric BC state occupies an intermediate position between the open and BC (sym) states, suggesting it may correspond to the “primed” or “flipped” states discussed above.

#### Step 3: Asymmetric channel opening via branched or linear gating pathways

The third step involves the opening of the receptor (Fig. 5). We find that open states of both homomeric and heteromeric receptors are invariably asymmetric with unobstructed pores, and that gating follows a branched pathway for α1β3 but linear routes for α1β3γ2 and β3 GABA_A_Rs. During these gating motions, the TMD spacing between subunits changes substantially, creating opportunities for lipids such as cholesterol, POPG, or other acyl chains to insert and act as “molecular lubricants”. This remodelling also generates new cavities that may be functionally important, as they could also serve as binding pockets for general anaesthetics^73^. The structural mechanisms underlying the action of these modulators remain unclear. This is expected because the structural aspects of the interactions between these compounds and GABA_A_Rs have been studied only under equilibrium conditions^48^, thus missing altogether the transient events taking place between the stable resting and long-lived desensitised end states. For GABA_A_R-β3, we show that HSM alone is a weak agonist, whereas co-application with etomidate accelerates rise times, increases peak amplitude, and enhances activation and desensitisation rates (Supplementary Fig. 5). Yet, structures at equilibrium reveal little difference between HSM- and HSM+ETM-bound GABA_A_R-β3^44,56^, suggesting that the functional effects of etomidate arise through interactions with short-lived inter-subunit spacings that form in the intermediate open states.

Intriguingly, in the presence of PIP_2_, both full-length α1β3 and α1β3γ2 GABA_A_ receptors failed to open, transitioning directly from the BC (sym) state to a long-lived desensitised state. Removal of the PIP_2_ binding site, however, allowed α1-containing receptors to open. Recent studies reveal that GABA_A_ receptors in the resting state are associated with cholesterol-rich lipid rafts but translocate to PIP_2_-rich microdomains upon GABA application^74^. This shift may regulate the pool of activatable receptors, preventing signalling by physically associating the receptors with PIP_2_. The balance between receptors in cholesterol- and PIP_2_-rich domains may thus influence synaptic strength and plasticity, acting as a feedback mechanism to limit prolonged inhibitory transmission during high-frequency neurotransmitter release. PIP_2_ levels in neurons can oscillate and be reduced by the activation of metabotropic GPCRs or the BDNF/TrkB signalling cascade and the activation of PLCγ^75,76^. In turn, the time that receptors reside in PIP_2_-rich microdomains will likely be shortened, therefore enhancing GABAergic signalling. PIP_2_ binding residues on the α1 subunit are shared with the α2, α3 and α5 subunits^9^, suggesting that these predominantly synaptic receptors will remain inactive in PIP_2_-rich environments, while extrasynaptic α4 and α6 subunits, which lack PIP_2_ binding sites^77^, remain constitutively activatable.

#### Step 4: Lipid-mediated asymmetric desensitisation

The fourth step is the entry into the short-lived desensitised states where lipids bind to the newly formed intra-subunit interfaces but also bind the pore thus obstructing the ion flow unveiling a novel short-lived closure mechanism within the Cys-loop receptor family occurring a few milliseconds after opening^2^ (Fig. 5a,c). The lipids are absent in the equilibrium resting and long-lived desensitized states, indicating that their presence is not an artefact of sample preparation but rather a transient phenomenon of likely physiological relevance. Such lipid-mediated pore lining has been previously reported in unrelated ion channels, suggesting that it might be a common phenomenon^78^.

#### Step 5: Entry into the long-lived symmetric desensitised state

For all receptors the last long-lived desensitised state transitions back to be quasi-symmetric with an open activation and closed desensitisation gate (Fig. 5).

Our analysis of GABA_A_R dynamics on the millisecond timescale reveals that ion channel gating proceeds through a series of asymmetric short-lived closed, open, and desensitised states, contrasting with the pseudo-symmetric conformations observed under equilibrium conditions. While previous studies on other pLGICs have suggested asymmetric states^36,37,39,79^, they were performed under equilibrium conditions and could not resolve the transient intermediates^17,18,80^. Applying this time-resolved single-particle cryo-EM approach more broadly may uncover conserved or divergent gating principles across the pLGICs family.

A key limitation, shared with earlier studies, is the isolation of receptors from their native membrane environment, where auxiliary proteins, the membrane’s potential and lipid composition may modulate receptor’s function^81^. Investigating receptors within intact membranes and ultimately in intact synapses will be necessary for a more comprehensive understanding of receptor gating in a physiological context. The combined temporal and spatial resolution achieved here establishes a mechanistic framework for interpreting receptor function and pharmacology. This approach can be used to guide the design of therapeutics targeting distinct binding interfaces and intermediate states in the GABA_A_ receptor gating cycle.

## Materials and Methods

### Construct design

All constructs with a truncated M3-M4 loop used in this study were ordered as synthetic cDNAs and codon-optimised for human cell expression. Construct truncated α1-1D4 was assembled from the following: residues 1 to 28 of the PTPRS secretion signal sequence (Uniprot ID: F1NWE3) followed by ETG to include the AgeI cloning site, followed by GABRA1 residues to 28 to 336 (Uniprot ID: P14867) followed by the short amino acid sequence SQPARAA^40,45^ to replace the M3-M4 loop, followed by GABRA1 residues 418 to 456 (Uniprot ID: P14867), followed by the KpnI cloning site and 1D4 tag. Construct truncated α1-SBP was assembled from the following residues: residues 1 to 28 of the PTPRS secretion signal sequence (Uniprot ID: F1NWE3) followed by an SBP-tag and a TEV cleavage site. The Age1 cloning site was introduced immediately after the TEV site. This was followed by GABRA1 residues to 28 to 336 (Uniprot ID: P14867) followed by the short amino acid sequence SQPARAA to replace the M3-M4 loop, followed by GABRA1 residues 418 to 456 (Uniprot ID: P14867). Construct truncated β3-SBP was assembled from the following residues: residues 1 to 28 of the PTPRS secretion signal sequence (Uniprot ID: F1NWE3) followed by an SBP-tag and a TEV cleavage site. The Age1 cloning site was introduced immediately after the TEV site. This was followed by GABRB3 residues 26 to 332 (Uniprot ID: P28472), followed by the short amino acid sequence SQPARAA to replace the M3-M4 loop, followed by GABRB3 residues 447 to 473 (Uniprot ID: P28472). Construct truncated γ2-1D4 was assembled from the following: residues 1 to 28 of the PTPRS secretion signal sequence (Uniprot ID: F1NWE3) followed by ETG to include the AgeI cloning site, followed by GABRG2 residues 39 to 360 (Uniprot ID: P18507), followed by the short amino acid sequence SQPARAA to replace the M3-M4 loop, followed by GABRG2 residues 447 to 475 (Uniprot ID: P18507), followed by the KpnI cloning site and 1D4 tag.

### Protein production and purification

#### Truncated α1β3 diheteromer and β3 homomer production

The truncated α1-1D4 and truncated β3-SBP genes were cloned into the pHL vector and produced by transient transfection following published procedures^8^. For 1L of Expi293F cells (Thermo Fisher Scientific), 1.1 mg of the required DNA was mixed with 33mL of Expi expression media (Gibco). In the case of the truncated α1β3, a plasmid mixture containing a 5:1 ratio of α1:β3 vectors was used to favour the presence of two α1 subunits in the pentameric GABA_A_ receptor assembly. The solution was mixed with 33mL of media supplemented with 3mL (1mg/mL) of PEI ‘MAX’ (PolySciences). After a 15min incubation at room temperature, the DNA+PEI mixture was added to the cell suspension at cell density of 2 million/mL. The cultures were then grown for 24h and harvested by centrifugation. The pellets were snap frozen in LN_2_ and stored at -80°C until further use.

#### Truncated α1β3γ2 triheteromer production

For the production of the truncated α1β3γ2 triheteromer, the truncated α1-SBP, truncated β3- SBP and truncated γ2-1D4 genes were cloned into the pHR-CMV-TetO2 vector^82^. For expression, a lentiviral cell pool was generated in HEK293S GnTI^-^ TetR cells as described previously^83^. Aiming for an invariant stoichiometry of β3/α1/γ2/β3/α1 receptors, the subunits were transfected in a ratio 10:1:1 = α1:β3:γ2. Cells were grown in FreeStyle 293 expression medium (Gibco) supplemented with 1% fetal bovine serum (Gibco), 1mM L-glutamine (Gibco), 1% NEEA (Gibco) and 5 μg/mL blasticidin (Invivogen) at 37 °C, 130 r.p.m., 8% CO2. Expression was induced by the addition of 2 μg/mL doxycycline. The cultures were then grown for additional 24h and harvested by centrifugation. The pellets were snap frozen in LN_2_ and stored at -80°C until further use.

#### Full-length α1β3 and α1β3γ2 production

The full-length α1β3 and α1β3γ2 were produced as described before^9,26,84,85^.

#### GABA_A_ receptor purification

The cell pellets were resuspended in PBS pH=7.4 supplemented with a cOmplete Protease Inhibitor Cocktail EDTA-free tablet (Roche) and the membranes solubilized with 1% (w/v) lauryl maltose neopentyl glycol (LMNG) for 1h. Cell debris and other insoluble material were removed by centrifugation (15,000×g, 15 min). The supernatant from 1L of cells was incubated with 300μL of the desired affinity resin: 1D4 resin, which was generated in-house by attaching anti-Rho-1D4 antibody (University of British Columbia) to CnBr-activated Sepharose 4B (GE Healthcare), for the full-length α1β3γ2L, truncated α1β3γ2 and truncated α1β3; Streptavidin agarose (Thermo Fisher) for the truncated β3 homomer; and FLAG M2 monoclonal antibody affinity gel (Sigma) for the full-length α1β3. For the truncated β3 homomer, the buffering solutions were kept at pH=8, while for the heteromeric receptors the solutions were kept at pH=7.4. The beads were gently mixed at 4°C for 2h and washed in batch with PBS (125 mM NaCl, 25 mM sodium phosphate), 0.1% (w/v) LMNG to remove impurities. The receptors were then reconstituted in MSP2N2 nanodisc. For the extrasynaptic full-length and truncated α1β3 and β3 GABA_A_Rs, the beads were incubated for 30min with 500μg of phosphatidylcholine (POPC, Avanti) and bovine brain lipid (BBE) extract (type I, Folch fraction I, Sigma-Aldrich) mixture (POPC:BBE = 70:30 w/w). 100μL MSP2N2 (5mg/mL) was then added to the beads and further incubated for 30min. For the synaptic full-length and truncated α1β3γ2, 8mM soluble cholesterol in the form of MβCD-cholesterol (Sigma) was added together with the lipids to increase the cholesterol content of the disc to ∼40% (see Supplementary discussion). Next, Bio-beads SM-2 (Bio-Rad) equilibrated in PBS were incubated with the resin for 90 minutes. The beads were then washed 6 times with 1mL of PBS and the receptors eluted overnight in 2.5mM Biotin, PBS pH=8 for Streptavidin resin, 100μg/mL FLAG peptide, PBS pH=7.4 for FLAG resin and 2mM 1D4 peptide, PBS pH=7.4 for 1D4 resin.

For the SMALPs reconstitution, the cells were resuspended, homogenised and spun at 15,000×g for 15 minutes to remove cell debris. The supernatant was collected and ultracentrifuged at 195,000×g for 1 hour to collect the membrane fraction. The membrane pellet was then resuspended in TBS (20mM Tris, pH=8, 150mM NaCl) and homogenised using a Dounce homogenizer. The membranes were resuspended in TBS, pH=8 at 80mg/mL. 1% (w/v) Xiran 30100 was then added to the solution and incubated for 2 hours at 4° C. Cell debris and insoluble material were removed by centrifugation (15,000×g, 15 min). The supernatant was then incubated for 2h with 1D4-affinity resin, washed 3 times with 50mL TBS, pH=8 and eluted with 5mM 1D4 peptide, TBS pH=8 overnight at 4°C.

### Cryo-EM sample preparation

Prior to freezing, purified truncated α1β3 and truncated α1β3γ2 receptors were incubated with 3μM Mb25^44^ or 2μM Nb38^40^ respectively to assist with particle alignment during cryo-EM processing. For the SMALPs datasets, the full-length α1β3γ2 was incubated with 2μM Mb38^26^. For the β3 homomer, full-length α1β3 and full-length α1β3γ2 time-resolved datasets no fiducial nanobody was added.

### Non-equilibrium grid preparation

For all time-resolved datasets, 3.5 μL of sample was applied to glow-discharged Quantifoil 1.2/1.3 (PELCO easiGlow, 30 mA for 60 s) and manually plunged at ∼8 °C and ∼95% humidity. The total ligand exposure time was limited to ∼10ms by adjusting the spraying nozzle to 5mm above the surface of the liquid ethane. The grid was released by free fall 35mm above the ethane level, and spraying was initiated by an optical trigger as previously described (Supplementary Fig. 1)^13^. The full-length α1β3γ2L was sprayed with 100mM GABA + 7mM ETM, 2mg/mL Ferritin (Sigma) in PBS pH=7.4; the full-length α1β3 was sprayed with 300mM GABA + 300mM HSM, 2mg/mL Ferritin in PBS pH=7.4; the truncated β3 was sprayed with 150mM HSM, 2mg/mL Ferritin in PBS pH=8; and the truncated α1β3 and truncated α1β3γ2 were sprayed with 300mM GABA, 2mg/mL Ferritin in PBS pH=7.4.

### Conventional grid preparation

For the SMALPs reconstituted receptor, 3.5 μL of sample was applied to a freshly glow-discharged (PELCO easiGlow, 30 mA for 120s) gold R1.2/1.3 300 mesh UltraAuFoil grid (Quantifoil) and blotted for 3.5s before plunge-freezing in liquid ethane using a Leica EM GP2 plunger (Leica Microsystems; 95% humidity, 14 °C). 1mM GABA and 10µM ETM were applied directly to the protein on the grid immediately before blotting resulting in ∼10s ligand exposure time.

### Cryo-EM data collection

All time-resolved cryo-EM datasets (apart from the full-length α1β3γ2L) were collected on a Titan Krios G2 microscope at the MRC LMB. The microscope was equipped with a Falcon 4 detector later upgraded to Falcon 4i. Data was collected in electron counting mode at 300kV and at a nominal magnification of 96,000 corresponding to a calibrated pixel size of 0.824 Å. The sprayed full-length α1β3γ2L was collected at eBIC on a Titan Krios G4 microscope equipped with a Falcon 4i detector. Images were acquired at 165,000 magnification corresponding to a calibrated pixel size of 0.723 Å The SMALPs dataset was collected at magnification of 75,000 with a pixel size of 1.055 Å. The resting state of the truncated α1β3γ2 was imaged on a Titan Krios G1 microscope at the MRC LMB equipped with a K3 detector in super-resolution bin2, nominal magnification of 105,000 and pixel size of 0.713 Å. All datasets were acquired using EPU (Thermo Fisher Scientific) with total exposure on the sample of ∼40 e^-^/Å^2^ and exposure time of 6.43s (Falcon 4, 96k magnification), 8.43s (Falcon 4, 75k magnification), 4.33s (Falcon 4i, 96k magnification), 4.31 (Falcon4i, 165k magnification, eBIC) or 1s (K3, 105k magnification). Before processing, all Falcon 4 EER images were converted to the tiff format using relion_convert_to_tiff and dose-fractionated into groups of 40 frames, corresponding to an accumulated dose of ∼1 e^-^/Å^2^ per frame. The K3 dataset was saved as tiff into groups of 37 frames, corresponding to an accumulated dose of ∼1.1 e^-^/Å^2^ per frame.

### Cryo-EM image processing

The movies were motion and gain corrected using MotionCor2^86^. Contrast transfer function (CTF) estimation was performed with CTFFIND-4.1.13^87^. Particle picking was performed using the neural networks WARP^88^, Topaz^89^ or the Laplacian of Gaussian autopicking function in RELION4.0^90^ and extracted in RELION4.0. The picked particles were subjected to 2D classification in cryoSPARC^91^. Particles resembling intact receptors from the 2D classes were used for ab-initio model generation with two seeds in cryoSPARC and were subjected to several rounds of heterogeneous refinement with four seeds to remove damaged particles or particles with preferential orientation. The best class following the multi-step classification was then refined with homogenous refinement and anisotropic magnification, per-particle defocus and 4^th^-order aberrations corrected.

The consensus particles were subjected to 3D classification with a focused TMD mask and no alignment with 30-40 seeds for each dataset. Each of the states classified by this approach were visually inspected and the structures that retained high-resolution features e.g. clear helical pitch, were subjected to homogenous refinement.

In the case of the β3 homomer, owing to the symmetry of the ECD region, the particles could not be aligned with the first consensus refinement prior to 3D classification. The 3D classification classes were subject to visual inspection, and those showing the same conformations but differing only in their rotation along the z-axis were manually aligned using the volume alignment tool function within cryoSPARC (Extended Data Fig. 3). For the α1β3 diheteromeric receptors, misalignment was resolved by having Mb25 bound to the ECD thus breaking the pseudo-symmetry and assisting the alignment (Extended Data Fig. 6). For the truncated α1β3γ2, both Mb25 and Nb38 were added to the sample to separate two populations of receptors with the following stoichiometries: β3/α1/γ2/β3/α1 and β3/β3/γ2/β3/α1, the latter lacking a second GABA binding site as the α1-D subunit is replaced by a β3 subunit (Extended Data Fig. 9)

The refined particles were subjected to a further round of heterogenous refinement with three classes using frequencies up to Nyquist. The best class was then subjected to non-uniform refinement and anisotropic magnification, per-particle defocus and 4^th^ order aberrations corrected. The particles were then polished in RELION4.0 and refined by non-uniform refinement in cryoSPARC which generated the final maps.

The final half-maps were subjected to post-processing in RELION4.0 for resolution estimation. All map and model rendering was performed in ChimeraX^92^. The pore profiles were calculated using HOLE^93^.

### Model building and refinement

The initial models used for the structures were the highest resolution reported models for both truncated and full-length subunits. Restraints for all ligands were generated by the Grade webserver (Global Phasing Ltd) using SMILES from PubChem as input. Iterative rounds of refinement and model building were performed in Coot v0.9.6^94^ and Phenix v1.19.2^95^. The models were validated by MolProbity v4.2^96^.

### Electrophysiology and heterologous expression

β3 GABA_A_R homomers (both full-length and truncated constructs) were heterologously expressed in transiently transfected adherent HEK-293 cells grown at 37°C and 5% CO_2_ in 35- mm cell-culture dishes. Transfections were performed using a calcium-phosphate-precipitation method and the same cDNAs (3.0 mg cDNA/dish) as those used for structural work. Cells were incubated with the cDNA-calcium-phosphate complexes for 16–18 hr after which the medium was replaced with fresh DMEM (ThermoFisher) containing 10 mM sodium butyrate and 10 mM sodium valproate to boost expression. Macroscopic currents were recorded in the whole-cell configuration of the patch-clamp technique at ∼22°C with an effective bandwidth of DC– 5 kHz using an Axopatch 200B amplifier (Molecular Devices). Currents were digitized at 10 kHz and analyzed using pCLAMP 11.1 software (Molecular Devices). Series-resistance compensation was used and set to ∼80%. The reference Ag/AgCl wire was connected to the extracellular solution through an agar bridge containing 200 mM KCl. Agonist-concentration jumps were applied to whole cells using a piece of double-barreled θ-tubing (Siskiyou). The flow of solution through the θ-tube was controlled using a gravity-fed system (ALA BPS-8; ALA Scientific Instruments), and the movement of the θ-tube was achieved using a piezoelectric arm (Burleigh-LSS-3100; discontinued) controlled by pCLAMP 11.1 software^97^ (Molecular Devices). The latter signals were low-pass-filtered (900C; Frequency Devices) at a cutoff frequency of 22 Hz prior to their arrival at the piezoelectric arm to reduce ringing in the θ-tube motion. During experiments, patched cells remained attached to a piece of collagen– coated glass coverslip (Neuvitro) placed at the bottom of the recording chamber. In this configuration, the perfusion system achieved a solution-exchange time of ∼0.9 ms for the onset (*t*_10 to 90%_) and ∼2.6 ms for the offset (*t*_90 to 10%_), as estimated from changes in the liquid-junction current (measured with an open-tip patch pipette) upon stepping the concentration of a KCl solution between 0.1 and 1 M. Although slower and more variable than the perfusion that can be achieved with excised patches^98^, the whole-cell configuration was favored here so as to increase the size of the currents. In an attempt to mimic the conditions used during cryo-EM imaging, both the intracellular (pipette) and extracellular (bath and perfusion) solutions had the same composition (125 mM NaCl, 25 mM sodium phosphate, pH=8.0) as that of the solution used here for structural work. Patch pipettes, pulled from thin-walled borosilicate-glass capillary tubing (Sutter Instrument), had resistances of ∼5 MΩ when filled with pipette solution.

## Data availability

Atomic coordinates for the models reported here have been deposited to Protein Data Bank the and the cryo-EM density maps have been deposited in the Electron Microscopy Data Bank. For truncated β3 homomeric GABA_A_ receptor sprayed with histamine or PBS: PDB accession codes 9FEU – 9FF2, 9G2D – 9G2E; Electron Microscopy Data Bank accession codes EMD-50341 – EMD-50349, EMD-50973 – EMD-50974. For truncated α1β3 diheteromeric GABA_A_ receptor sprayed with GABA or PBS: PDB accession codes 9FFL – 9FFU; Electron Microscopy Data Bank accession codes EMD-50365 – EMD-50374. For truncated α1β3γ2 triheteromeric GABA_A_ receptor sprayed with GABA: PDB accession codes 9FFV – 9FG3; Electron Microscopy Data Bank accession codes EMD-50375 – EMD-50383. For full-length α1β3 diheteromeric GABA_A_ receptor sprayed with GABA and histamine: PDB accession codes 9FG4 – 9FG6; Electron Microscopy Data Bank accession codes EMD-50384 – EMD-50386. For full-length α1β3γ2 triheteromeric GABA_A_ receptor sprayed with GABA and etomidate: PDB accession codes 9FG7 – 9FG9; Electron Microscopy Data Bank accession codes EMD-50387 – EMD-50389. For full-length α1β3γ2 triheteromeric GABA_A_ receptor in SMALPs, Saposin A or large MSP2N2 nanodiscsc: PDB accession codes 9FGA – 9FGH; Electron Microscopy Data Bank accession codes EMD-50390 – EMD-50393, EMD-50397, EMD-50399 – EMD-50402.

## Acknowledgements

We thank J. Grimmett, T. Darling and I. Clayson for scientific computing support; S. Chen, G. Cannone, G. Sharov, A. Yeates and B. Ahsan for electron microscopy support; N. Unwin for valuable insights into sample preparation and access to a custom-built cryo-EM grid plunger; K.W Miller for providing advice and the stable cell lines expressing full-length α1β3γ2 and α1β3 GABA_A_ receptors; S. Lövestam, C. Russo, N. Unwin and V. Kasaragod for helpful discussions and comments on this work. We acknowledge funding from the UK Medical Research Council (grants MR/L009609/1, MC_UP_1201/15 and MC_EX_MR/T046279/1 to A.R.A.), the National Institute for General Medical Sciences (1R01-GM135550 to A.R.A.), the National Science Foundation (NeuroNex 2014862 to A.R.A.) and the School of Molecular and Cellular Biology of the University of Illinois Urbana-Champaign to C.G.

## Author contributions

D.B.M. and A.R.A. conceived the project. D.B.M. and V.C. generated cell lines. D.B.M. characterised the cell lines. D.B.M. purified the proteins, prepared the cryo-EM samples, collected and processed the cryo-EM data. T.M. built and refined the atomic models. G.D.C. and C.G. performed and analysed the electrophysiology experiments. J.M. purified and solved the GABA_A_ receptor structures in large MSP2N2 nanodiscs. K.N. calculated the millisecond-scale diffusion rates. D.B.M. and A.R.A. wrote the manuscript, with input from all authors.

## Competing interests

A.R.A. and V.T.C. are part-time employees of BioNTech UK Ltd. However, the work reported here was performed independently of this affiliation. The other authors declare that they have no competing interests.

