## Extended Data Figures for "GABA_A_ receptor gating imaged on the millisecond timescale"

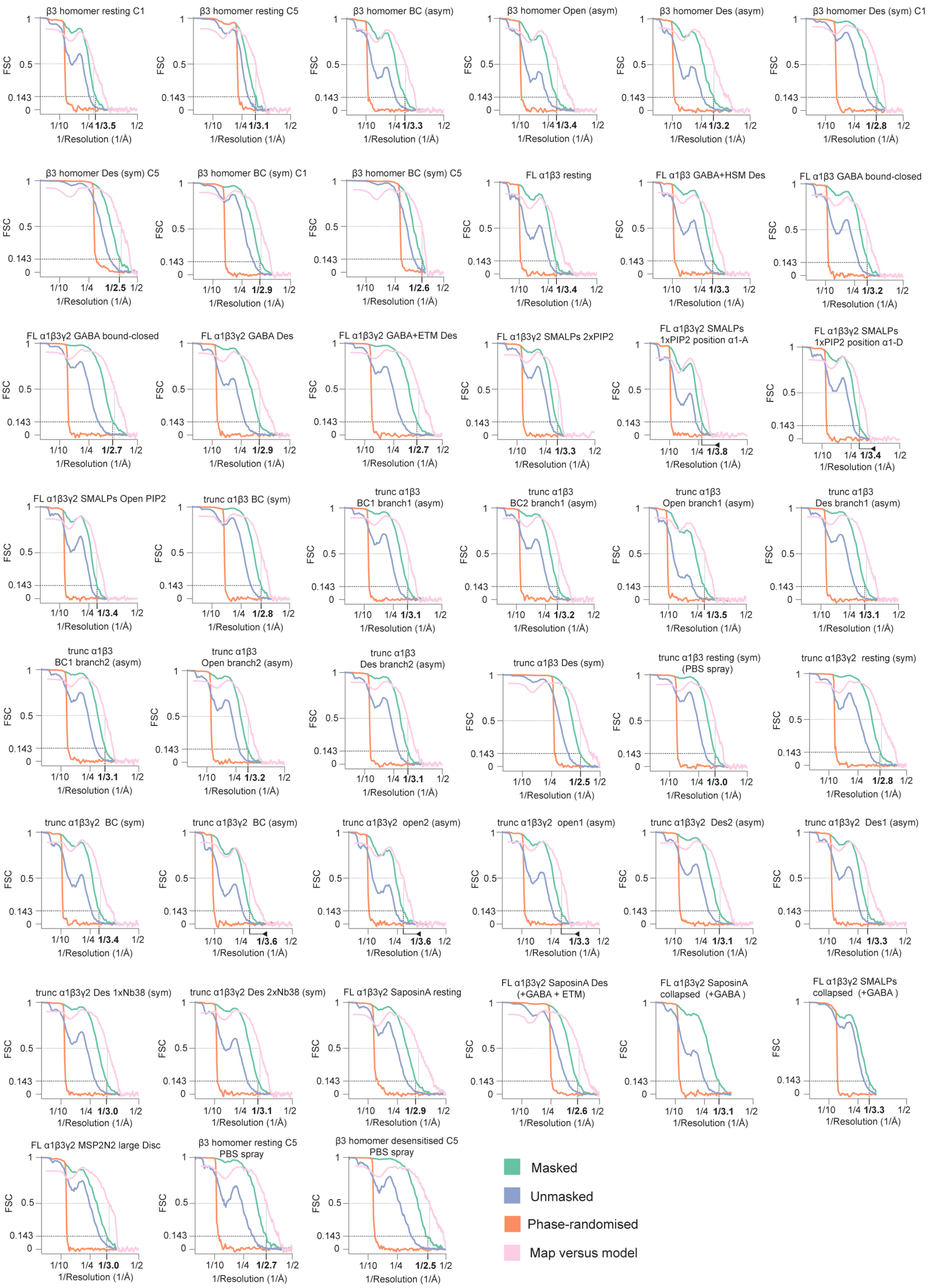


**Extended Data Figure 1 | Quality of cryo-EM maps.**

FSC plots for all maps and models. Each plot contains masked (green), unmasked (blue), phase-randomized (orange) and map vs. model Fourier shell correlation (FSC) curves.

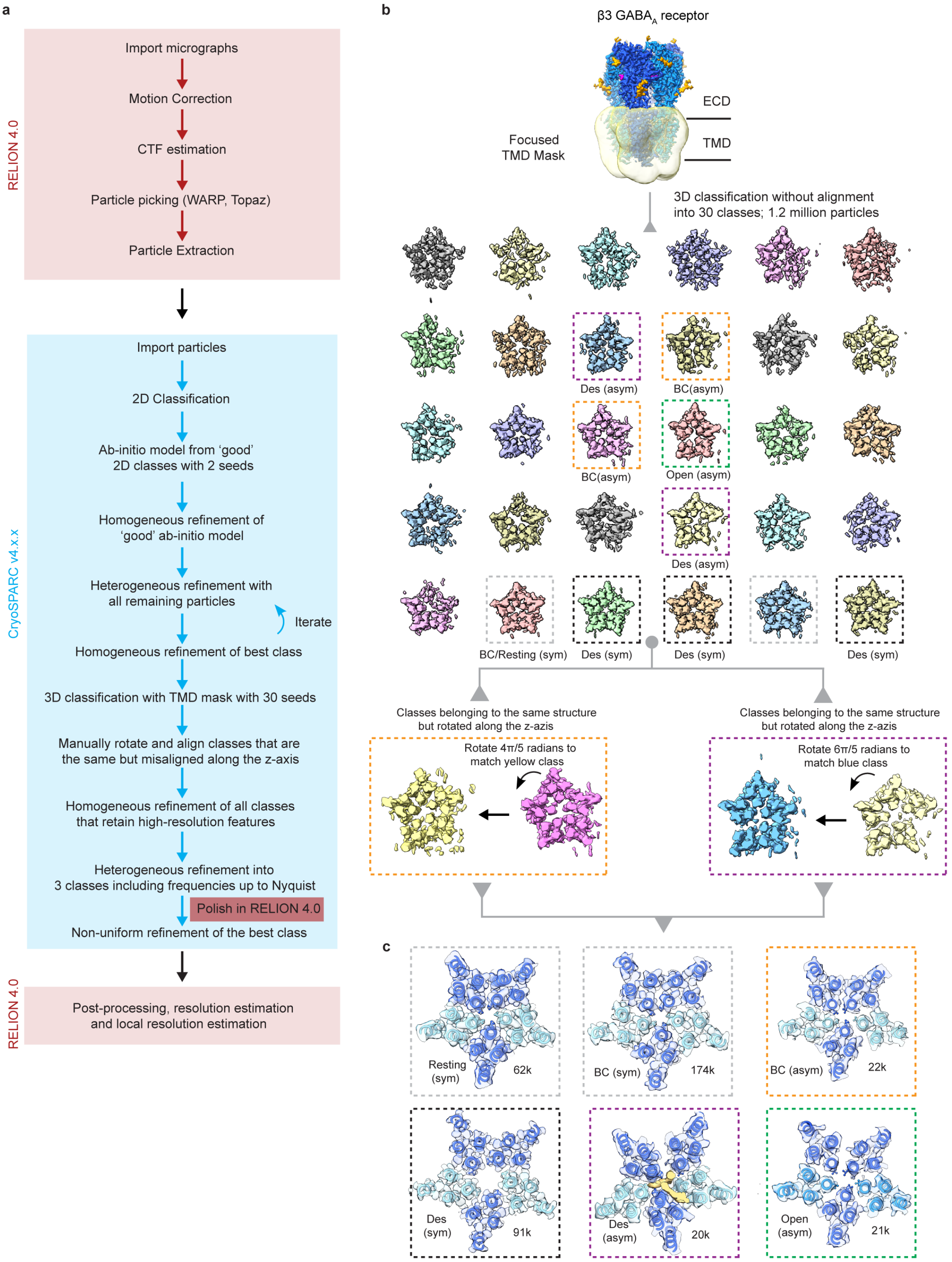


**Extended Data Figure 2 | β3 homomer processing pipeline.**

**a,** The detailed processing pipeline used for the β3 homomer. **b,** A soft TMD mask was used to classify the multitude of states. In total 30 seeds were used. Classes with well-defined high-resolution features i.e clear helical pitch, were subjected to further refinement. Classes which are the same but rotated along the z-axis were manually aligned. **c,** Cross-sections at the level of the 9′ activation gate for the final maps including the number of particles in the final reconstruction.

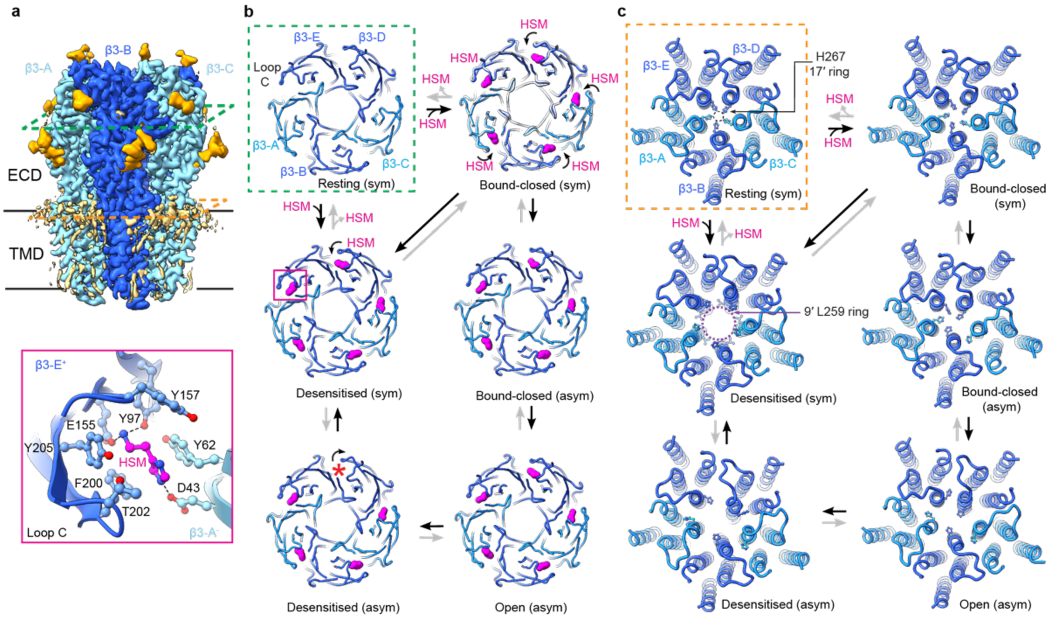


**Extended Data Figure 3 | The β3 homomer gating motions.**

**a,** Cryo-EM map of the β3 homomer viewed parallel to the membrane plane highlighting the plane of the HSM binding sites in green and the plane of the upper part of the TMD at the level of the headgroups of the outer leaflet in orange (top). One HSM-binding pocket viewed from the extracellular space (bottom). The amino acid side chains lining the binding site are shown as ball and sticks. Dashed lines indicate hydrogen bonds. **b,** Views of the ECD cross-sectioned at the level of the HSM binding pocket showing the ligand occupancy of each state. Pockets labelled with a red Asterix indicate the absence of HSM **c,** Cross-section of the TMD depicting the movement of His267.

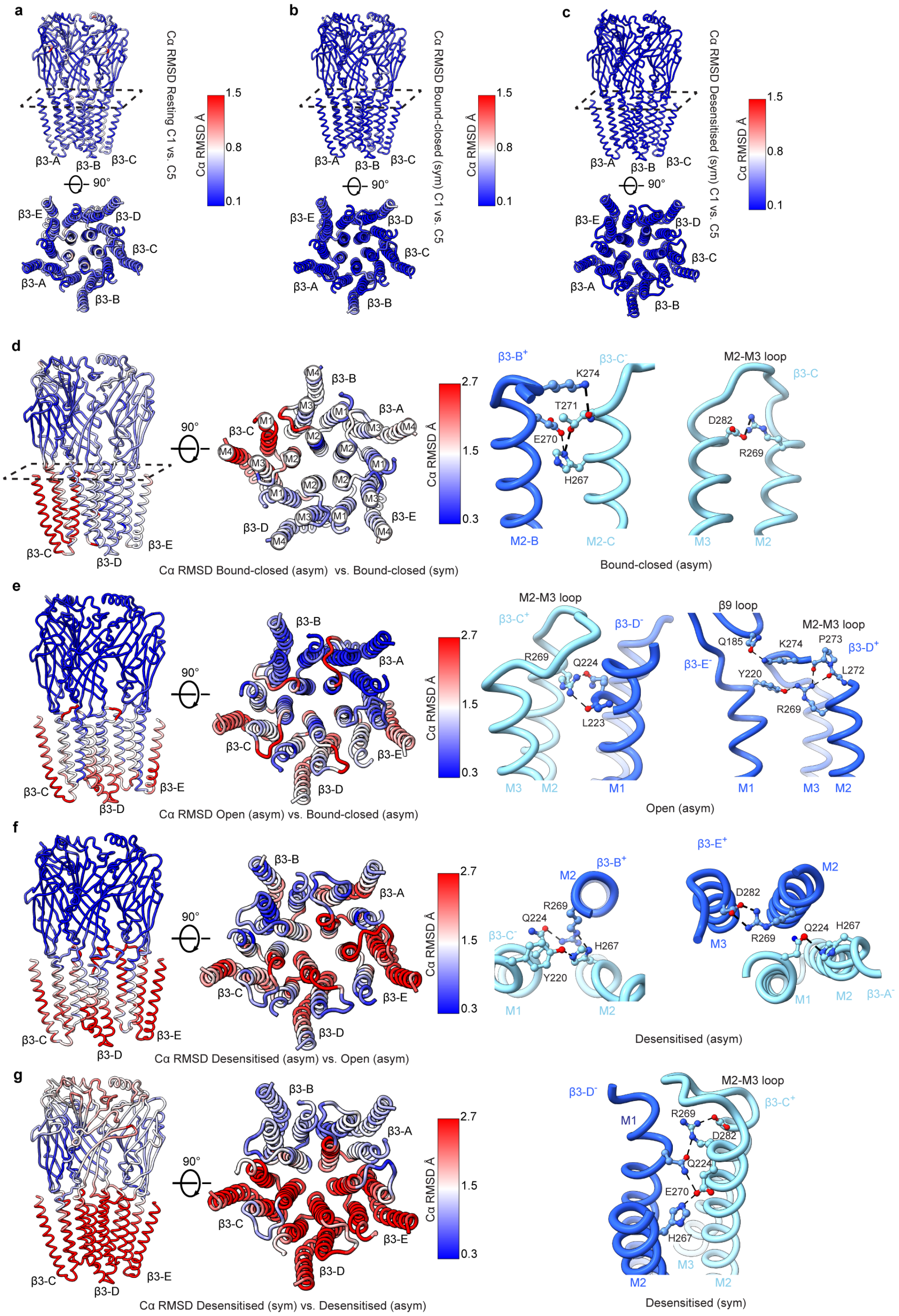


**Extended Data Figure 4 | The mechanism of β3 homomer gating.**

**a-c,** Plots of the per-residue Cα r.m.s.d. between the receptors processed in C1 and C5 symmetry for **(a)** the resting state, **(b)** the bound-closed state and **(c)** the long-lived desensitised state. **d,** Plots of the per-residue Cα r.m.s.d. between the bound-closed (asymmetric) and bound-closed (symmetric) states (left) and key residue interactions that stabilise the bound-closed (asymmetric) state (right). **e,** Plots of the per-residue Cα r.m.s.d. between the open (asymmetric) and bound-closed (asymmetric) states (left) and key residue interactions that stabilise the open (asymmetric) state (right). **f,** Plots of the per-residue Cα r.m.s.d. between the desensitised (asymmetric) and open (asymmetric) states (left) and key residue interactions that stabilise the desensitised (asymmetric) state (right). **g,** Plots of the per-residue Cα r.m.s.d. between desensitised (symmetric) and desensitised (asymmetric) states (left) and key residue interactions that stabilise the desensitised (symmetric) state (right).

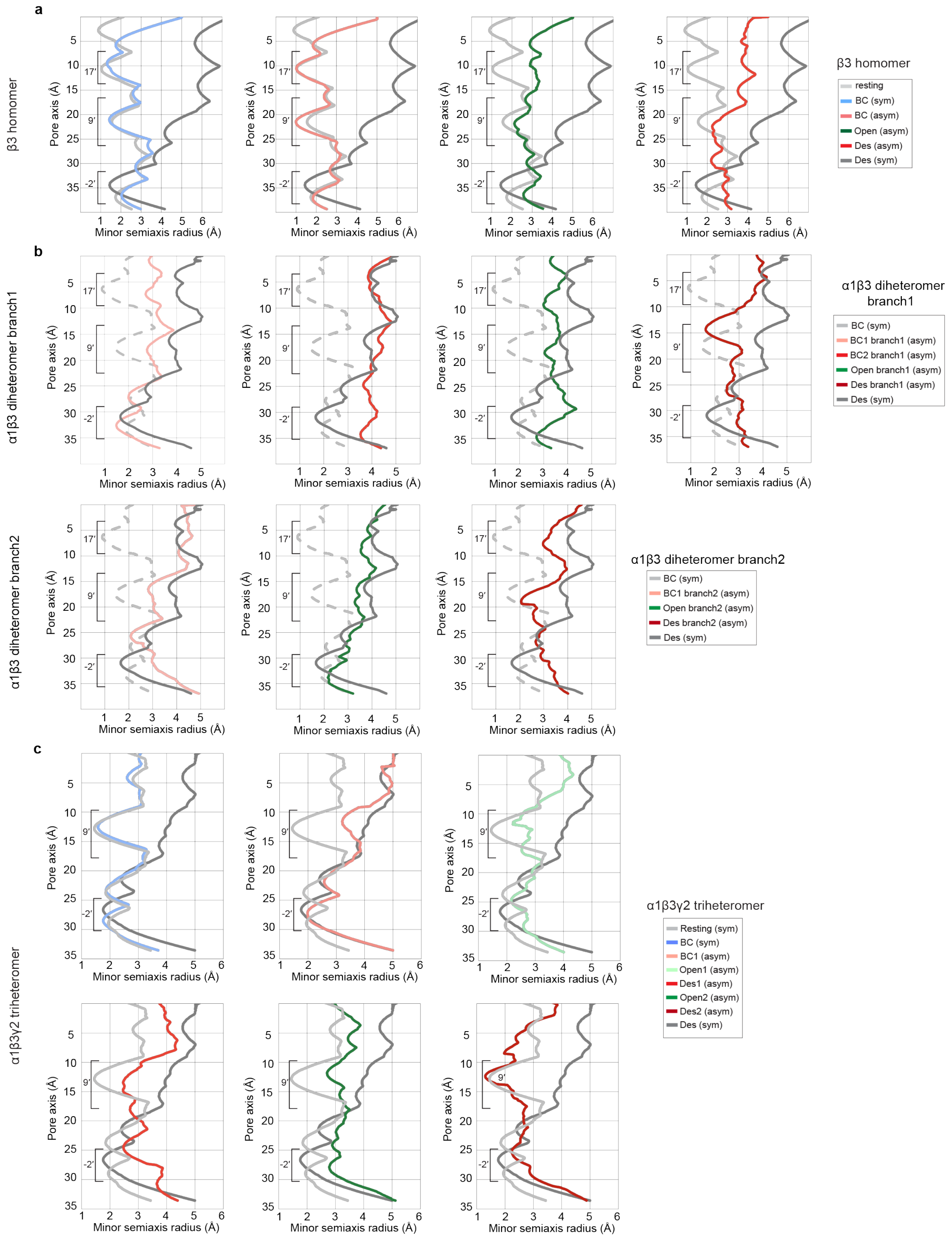


**Extended Data Figure 5 | Pore profiles.**

Pore profiles of all states classified on the millisecond timescale for the **(a)** β3 homomer; **(b)** α1β3 diheteromer and **(c)** α1β3γ2 triheteromer.

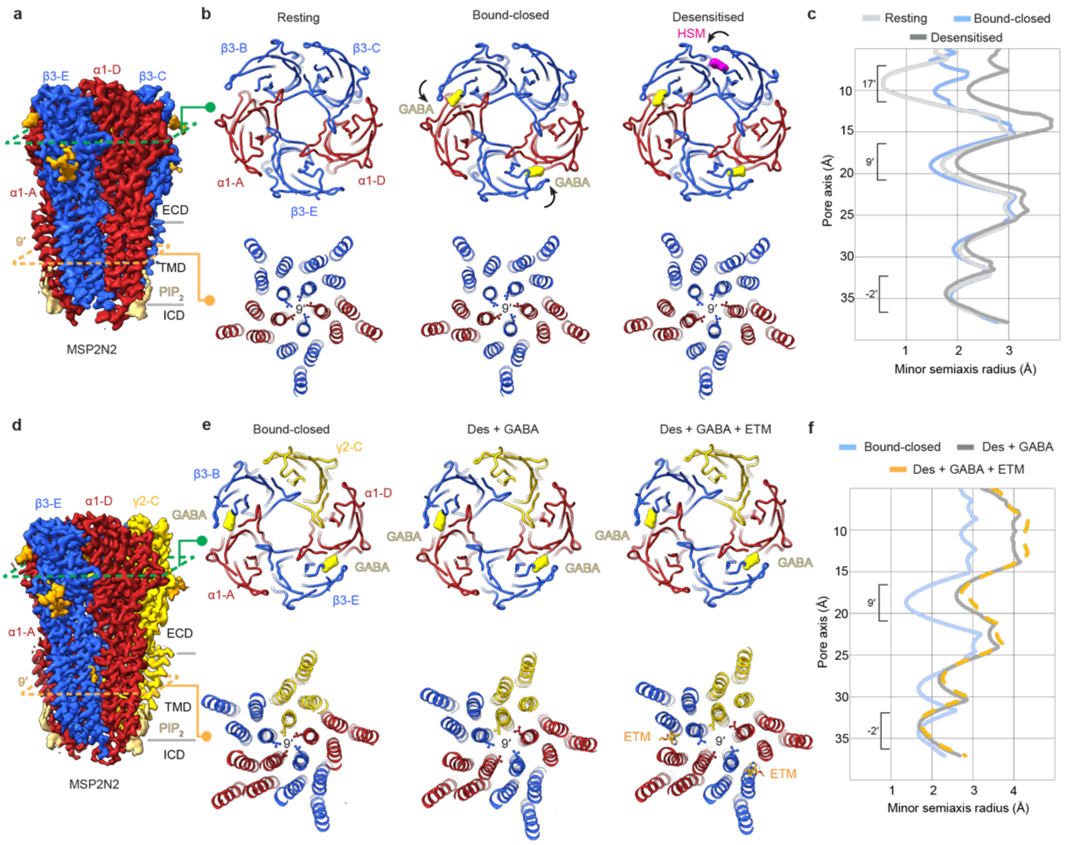


**Extended Data Figure 6 | $PIP_2$ regulates the opening of the full-length α1β3 and α1β3γ2 receptors.** **a,** Cryo-EM map of the full-length α1β3 in MSP2N2 nanodisc viewed parallel to the membrane. **b,** Cross-section at the level of the agonist binding pocket of the full-length α1β3 resting state (top-left), bound-closed state (top-middle) and desensitised state (top-right). Cross-section at the level of the 9′ activation gate of the full-length α1β3 resting state (bottom-left), bound-closed state (bottom-middle) and desensitised state (bottom-right). **c,** Pore profiles of the full-length α1β3 resting, bound-closed and desensitised states. **d,** Cryo-EM map of the full-length α1β3γ2 in MSP2N2 nanodisc viewed parallel to the membrane. The etomidate pocket is marked in orange **e,** Cross-section at the level of the agonist binding pocket of the full-length α1β3γ2 bound-closed state (top-left), desensitised state with GABA (top-middle) and desensitised state with GABA and etomidate (top-right). Cross-section at the level of the 9′ activation gate of the full-length α1β3γ2 bound-closed state (bottom-left), desensitised state with GABA (bottom-middle) and desensitised state with GABA and etomidate (bottom-right). **f,** Pore profiles of the full-length α1β3γ2 bound-closed, desensitised with GABA and desensitised with etomidate state.

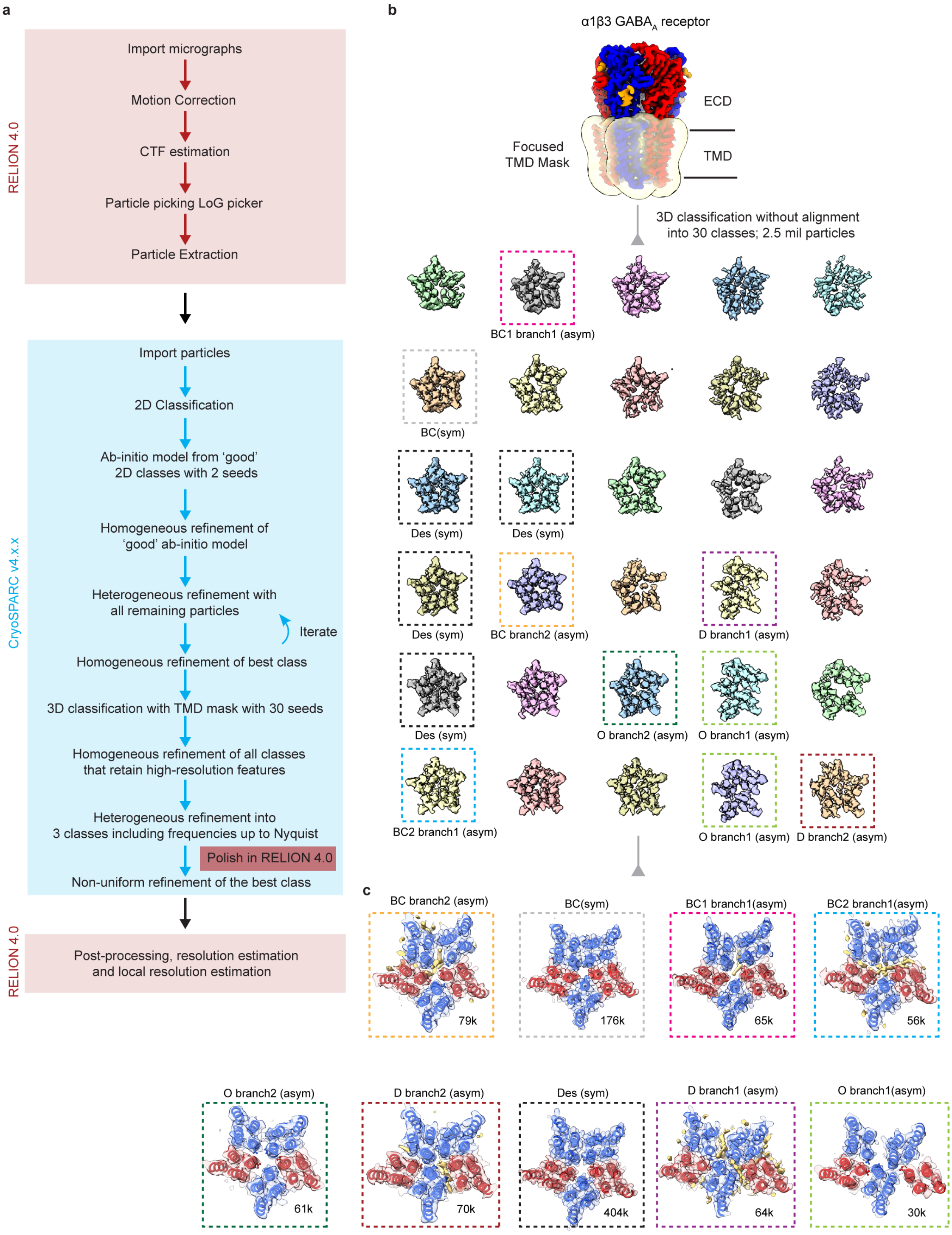


**Extended Data Figure 7 | α1β3 diheteromer processing pipeline.**

**a,** The detailed processing pipeline used for the α1β3 diheteromer. **b,** A soft TMD mask was used to classify the multitude of states. In total 30 seeds were used. Classes with well-defined high-resolution features i.e clear helical pitch, were subjected to further refinement. **c,** Cross-sections at the level of the 9′ activation gate for the final maps including the number of particles in the final reconstruction.

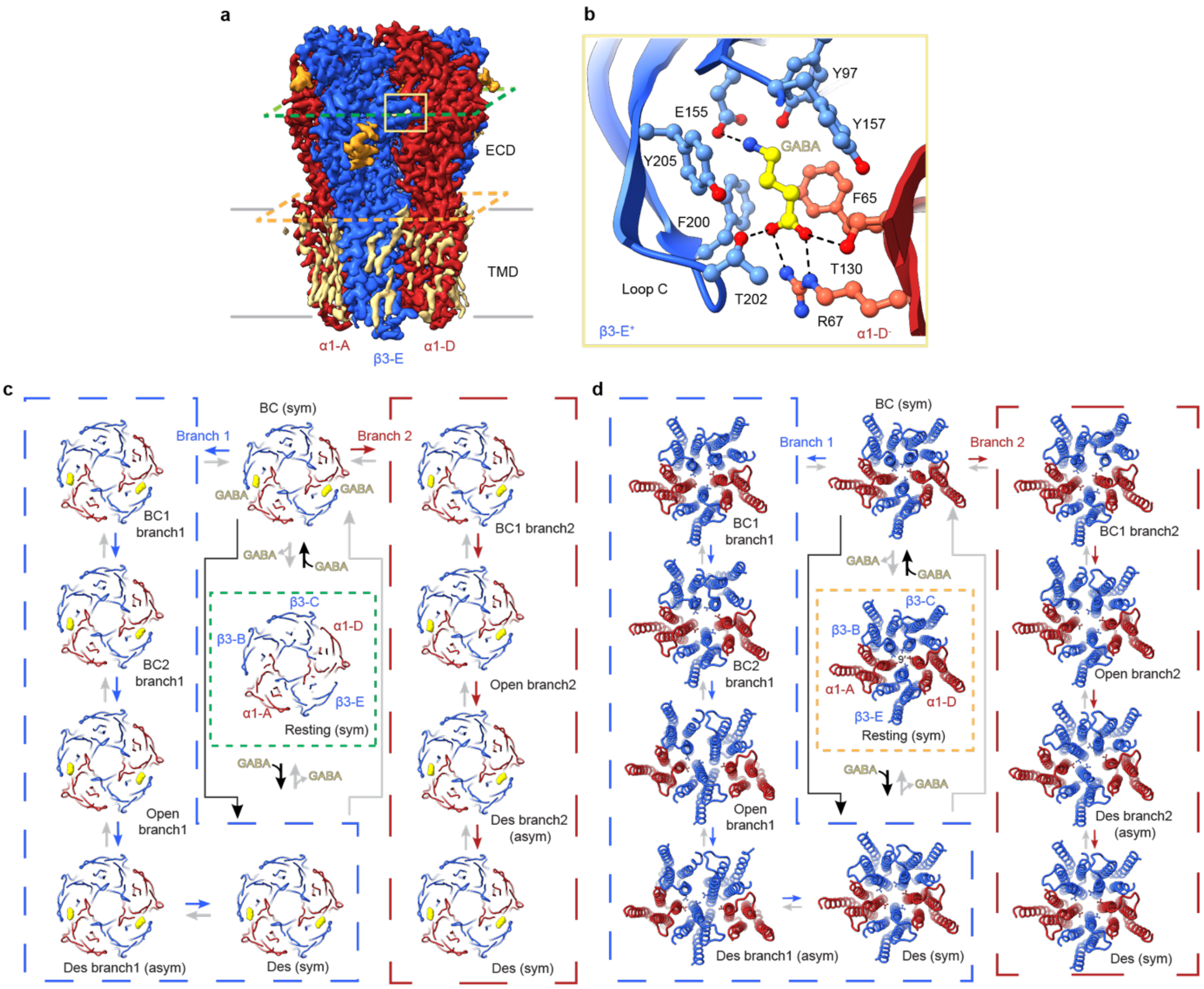


**Extended Data Fig. 8 | The α1β3 diheteromer gating motions.**

**a,** Cryo-EM map of the α1β3 diheteromer viewed parallel to the membrane plane highlighting the plane of the GABA binding sites in green and the plane of the upper part of the TMD at the level of the headgroups of the outer leaflet in orange. **b,** One GABA-binding pocket viewed from the extracellular space. The amino acid side chains lining the binding site are shown as ball and sticks. Dashed lines indicate hydrogen bonds. **c,** Views of the ECD cross-sectioned at the level of the GABA binding pocket showing the ligand occupancy of each state. **d,** Cross-section of the TMD depicting the motions of the alpha helices.

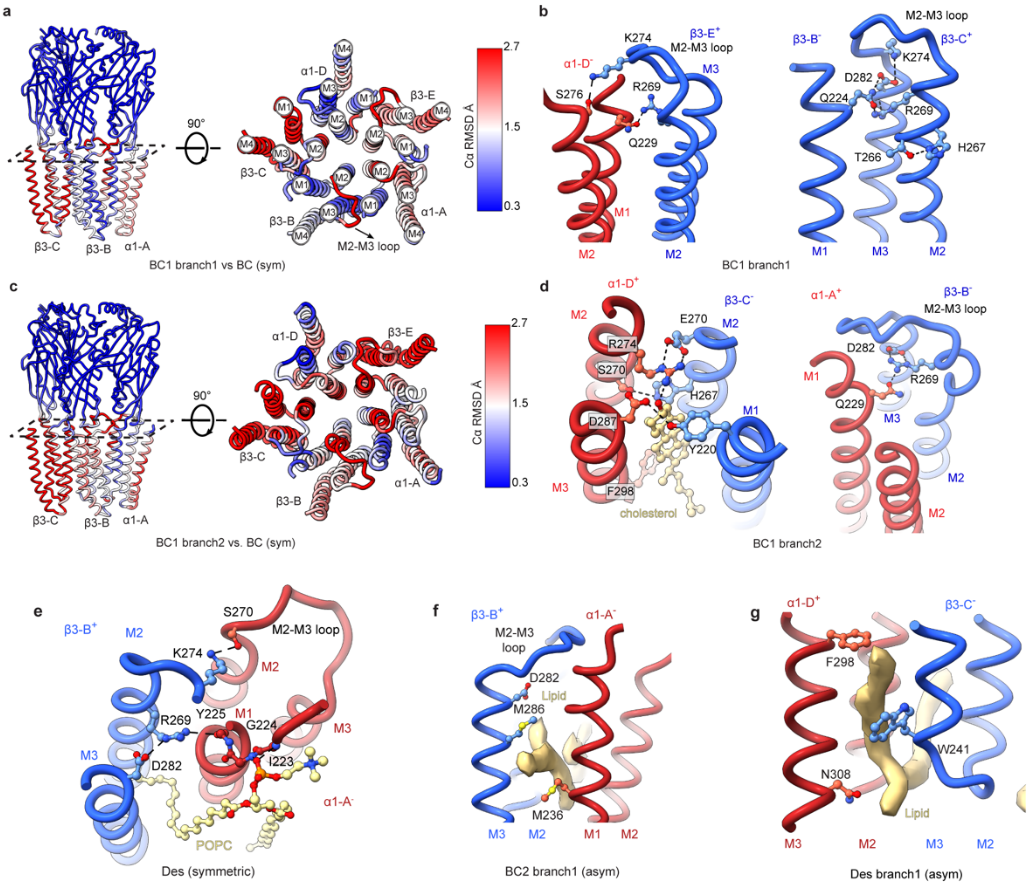


**Extended Data Fig. 9 | The mechanism of α1β3 gating.**

**a,** Plots of the per-residue Cα r.m.s.d. between the bound-closed branch 1 (asymmetric) and bound-closed (symmetric). **b,** Key residue interactions that stabilise the bound-closed branch 1 (asymmetric) state. **c,** Plots of the per-residue Cα r.m.s.d. between the bound-closed branch 2 (asymmetric) and bound-closed (symmetric). **d,** Key residue interactions that stabilise the bound-closed branch 1 (asymmetric). **e,** Key residue interactions that stabilise the desensitised (symmetric) state. POPC is shown in khaki. **f,** Unidentified lipid stabilising the $\beta3\text{-B}^{+}/\alpha1\text{-A}^{-}$ TMD interface of bound-closed 2 branch1 (asymmetric). **g,** Unidentified phospholipid stabilising the $\alpha1\text{-D}^{+}/\beta3\text{-C}^{-}$ TMD interface of desensitised branch2 (asymmetric).

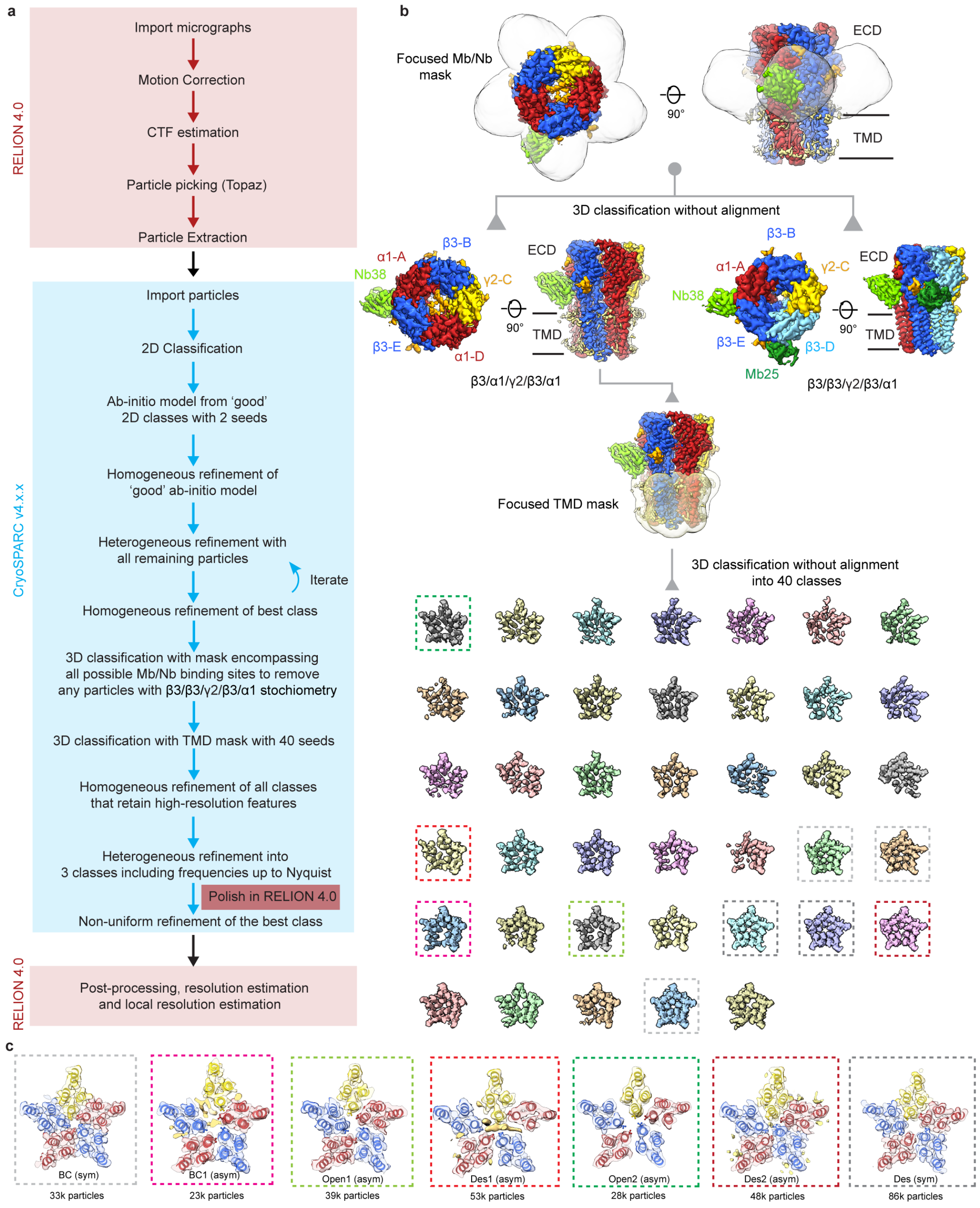


**Extended Data Fig. 10 | α1β3γ2 triheteromer processing pipeline.**

**a,** The detailed processing pipeline used for the α1β3γ2 triheteromer. **b,** A soft Mega/nanobody mask was used to classify the two different receptor assemblies present. The particles were subjected to 3D classification without alignment with 4 seeds. Particles belonging to the β3-E/α1-D/β3-C/β3-B/α1-A with both Mb25 and Nb38 bound were not processed further. Particles belonging to the β3-E/α1-D/γ2-C/β3-B/α1-A class with just Nb38 bound were subjected to a 3D classification with a TMD soft mask. In total 40 seeds were used. Classes with well-defined high-resolution features i.e clear helical pitch, were subjected to further refinement. **c,** Cross-sections at the level of the 9′ activation gate for the final maps including the number of particles in the final reconstruction.

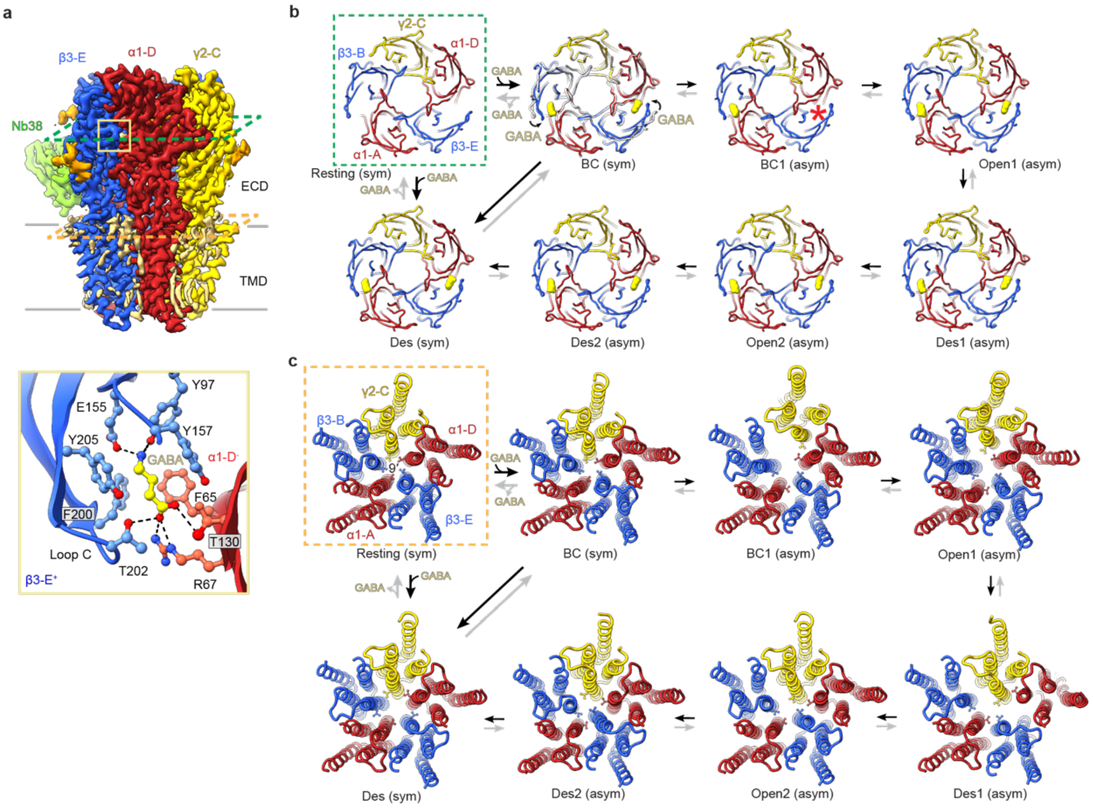


**Extended Data Fig. 11 | The α1β3γ2 triheteromer gating motions.**

**a,** Cryo-EM map of the α1β3γ2 triheteromer viewed parallel to the membrane plane highlighting the plane of the GABA binding sites in green and the plane of the upper part of the TMD at the level of the headgroups of the outer leaflet in orange (top)**.** One GABA-binding pocket viewed from the extracellular space. The amino acid side chains lining the binding site are shown as ball and sticks. Dashed lines indicate hydrogen bonds (bottom). **b,** Views of the ECD cross-sectioned at the level of the GABA binding pocket showing the ligand occupancy of each state. Pockets labelled with a red Asterix indicate the absence of GABA **c,** Cross-section of the TMD depicting the movement of the alpha helices.

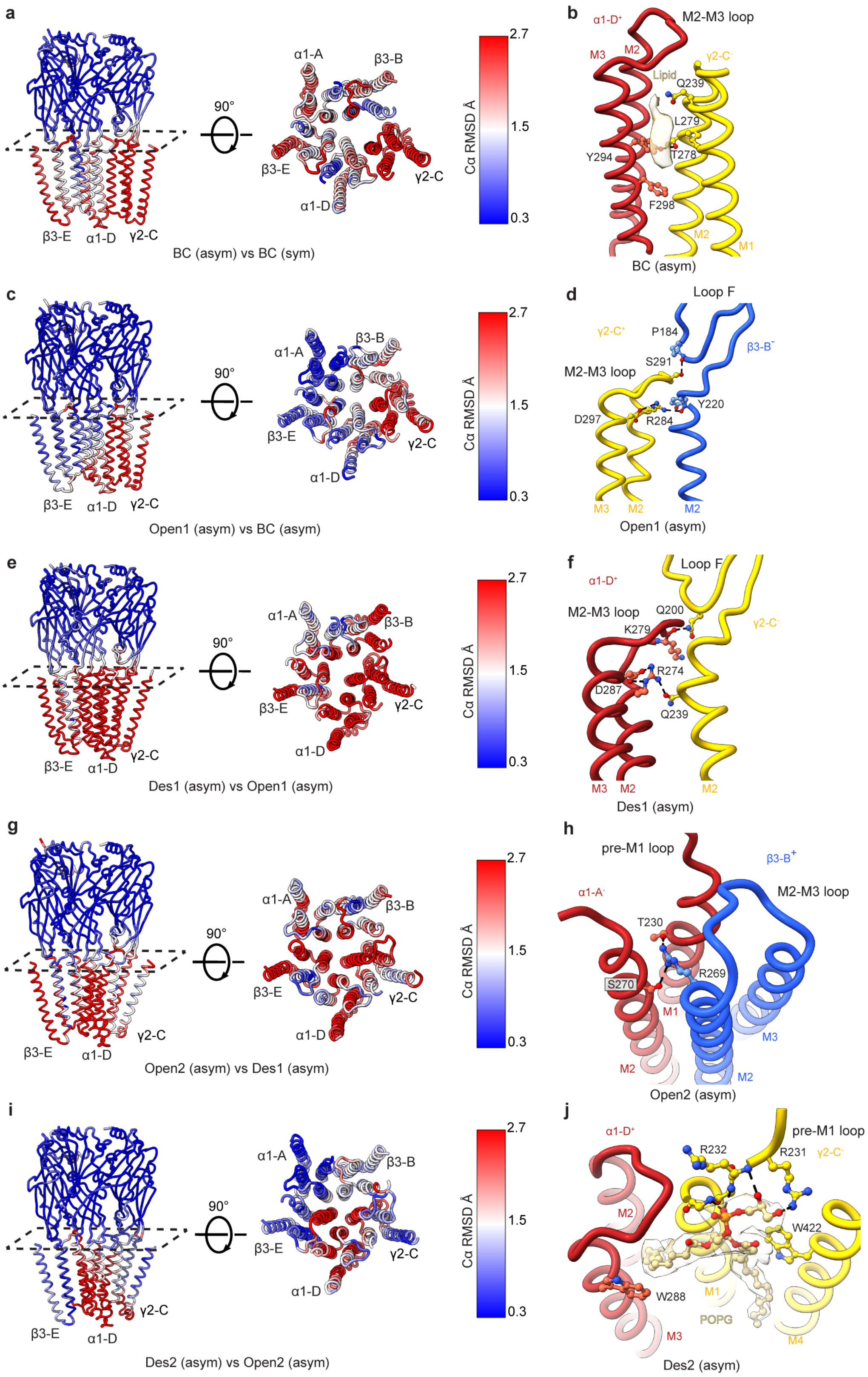


**Extended Data Fig. 12 | The mechanism of α1β3γ2 gating.**

**a,** Plots of the per-residue Cα r.m.s.d. between the bound-closed (asymmetric) and bound-closed (symmetric). **b,** Key residue interactions that stabilise the bound-closed (asymmetric) state. **c,** Plots of the per-residue Cα r.m.s.d. between the open1 (asymmetric) and bound-closed (asymmetric). **d,** Key residue interactions that stabilise the open 1 (asymmetric) state. **e,** Plots of the per-residue Cα r.m.s.d. between the desensitised 1 (asymmetric) and open 1 (asymmetric). **f,** Key residue interactions that stabilise the desensitised 1 (asymmetric) state. **g.** Plots of the per-residue Cα r.m.s.d. between the open 2 (asymmetric) and desensitised 1 (asymmetric). **h,** Key residue interactions that stabilise the open 2 (asymmetric) state. **i,** Plots of the per-residue Cα r.m.s.d. between the desensitised 2 (asymmetric) and open 2 (asymmetric). **j,** Key residue interactions that stabilise the desensitised 2 (asymmetric) state.
