## Supplementary Information for "GABA_A_ receptor gating imaged on the millisecond timescale"

### Supplementary Information Content

### 1. Supplementary discussion

#### 1.1 Time-resolved cryo-EM on the millisecond timescale - experimental setup and considerations

The experimental setup was similar to that described previously[1], with the grid placed 35 mm above the liquid ethane level and an atomizer spray adjusted to 5 mm above the surface of the ethane (Supplementary Fig. 1a-b). The atomizer sprayer is controlled by an optical switch at the top of the plunger and generates a jet of small droplets containing the ligand.

Using Newton's law of gravity one can calculate how long it takes for the grid to plunge: $d = \frac{1}{2}gt^2$, where $d$ is the distance the object has to travel, $g$ is the gravitational constant of 9.8 m/s$^2$, and $t$ is the time it takes to travel the distance. Therefore, at free fall from 35 mm, the grid will be plunged in 84 ms. Since the sprayer is located 5 mm above the ethane's surface, the grid will reach the spray droplets in 76 ms. This therefore limits the exposure to the ligand to ~10 ms. The resulting droplets on the grid varied in size but, typically, had a radius of around 2-5 µm (Supplementary Fig. 1c-d). When 2 mg/mL ferritin—chosen for its iron core readily detectable via electron microscopy—was added to the ligand, one could observe it diffusing to the edge of the droplet, where the ice is substantially thick (Supplementary Fig. 1e). This limits the achievable resolution by cryo-EM because more inelastic scattering takes place [2]. To overcome this, data need to be collected just next to the ligand droplet, where no ferritin molecules can be seen (Supplementary Fig. 1f). Nevertheless, since GABA is smaller, it will diffuse further owing to its larger diffusion coefficient, $D_{GABA} = 0.8$ µm$^2$/ms [3,4], whereas $D_{ferritin} = 0.01$ µm$^2$/ms [1,5]. To determine the concentration of the diffusible molecule, one can use the equation for diffusion from a cylindrical source such as the sprayed ligand droplet [6].

$$C = \frac{C_0}{2Dt} e^{\left(-r^2/_{4Dt}\right)} \int_0^a e^{\left(-r'^2/_{4Dt}\right)} I_0\left(\frac{rr'}{2Dt}\right) r'dr'$$

In this equation, $C$ and $C_0$ are the final and initial concentrations of the ligand, respectively, $t$ is the time for diffusion (in this case, 10 ms), $D$ is the diffusion coefficient of the molecule in question, $r$ is the total diffusible distance from the center of the droplet, $a$ is the initial radius of the droplet, and $I_0$ is the Bessel function of the first kind of order zero. This equation was fitted to the available experimental data for ferritin and falls well in line with it[1]. Since the ice is only thin immediately next to the droplet, and droplets differ in size, the concentration of GABA will also differ (Supplementary Fig. 1d). The ice thickness where high-resolution imaging can be performed occurs 1−3 μm away from the edge of the droplet (Supplementary Fig. 1f). Therefore, in the case of a small droplet with a 2-μm radius, the concentration of GABA in this optimal region will range from $0.06\times C_0$ to $0.02\times C_0$. Moreover, the duration of exposure to ligand is a function of the distance from the center of the droplet, thus allowing exposure times as short as 2 ms or even less. As the droplet becomes larger, the concentration of GABA near the droplet also increases, and a droplet with a radius of 5 μm will therefore have $0.3\times C_0$ to $0.1\times C_0$ of GABA immediately next to it (Supplementary Fig. 1g).

The calculations above, however, assume diffusion in 3D space, which is not the case for a grid. On a grid, the ligand must also diffuse over the carbon support, which is covered by a thin layer of liquid. This also assumes the source to be cylindrical, which is not the case, either. Lastly, because the droplet is flying at high speed, the liquid can first expand and then retract upon impacting on the grid, which may further limit the diffusion of agonist[7]. Therefore, to attain a maximal open probability regardless of the size of the droplet, the ligand concentration in sprayed solutions was 100−300 mM.

#### 1.2 Buffer application on the millisecond timescale with no agonist has no structural impact

To assess the physiological relevance of the asymmetric short-lived intermediate states reported in this study, we subjected the receptors to buffer alone during spraying. This negative control was designed to elucidate the structural impact of rapid buffer diffusion on the receptor and determine whether the observed states are solely induced by the presence of agonist. All ligands were diluted in PBS pH=8.0 pr pH=7.4, thus we utilized the same buffer for spraying the negative control grids.

As anticipated, no states other than the resting state were present for the truncated α1β3 $GABA_A$ receptor sprayed with only PBS pH=7.4 (Supplementary Fig. 13). The receptor exhibited both activation and desensitisation gates closed, indicative of its resting state, as corroborated by the pore profile and absence of ligand density in the orthosteric $\beta3^{+}/\alpha1^{-}$ ligand binding pockets in the ECD. The Cα RMSDs in the TMD region between the bound-closed symmetric state and the resting state of α1β3 were consistently below 0.5 Å, suggesting that although agonist binding to the ECD is primed in the BC (sym), the associated structural changes have not yet propagated to the TMD. Importantly, this is true for all receptors reported in this study including the β3 homomer (Fig. 1, Extended Data Fig. 3) and the α1β3γ2 triheteromer (Fig. 4, Extended Data Fig. 10) where HSM for the β3 homomer or GABA for the α1β3γ2 triheteromer prime the receptor in the symmetric bound-closed state before it either enters the intermediate asymmetric states or immediately reaches the symmetric desensitised state.

We further extended these control conditions to the β3 homomer as well and sprayed with PBS pH=8. Compared to α1β3γ2 and α1β3 receptors, β3 homomers exhibit a higher spontaneous activity[8]. This means that the latter exist in equilibrium between the resting and desensitised states, with a small fraction of the receptors open at any time. The mechanism of this spontaneous activity involves proton gating via the His267 17′ gate. When His267 is protonated, it interacts with Glu270 through hydrogen bonds. This interaction activates the receptor, eventually bringing it to the desensitised state[9]. As was the case for the α1β3 diheteromer, no short-lived asymmetric state was observed upon spraying the homomer with

PBS pH=8 (Supplementary Fig. 13). Furthermore, 40% of the particles were in the long-lived symmetric resting state, and 60% were in the long-lived symmetric desensitised state as anticipated due to the spontaneous activity of this receptor type. Therefore, we conclude that merely spraying the grids has no structural impact on the receptor, and that the molecular rearrangements we observed were indeed triggered by the application of the agonist on the millisecond timescale.

### 1.3 The structural impact of the nanodisc scaffold on the reconstituted $GABA_AR$

Choosing a suitable nanodisc scaffold is a crucial consideration to ensure that reconstituted receptors can illustrate functionally meaningful states. All the structures we have presented in this study were solved in nanodiscs formed by the MSP2N2 scaffolding protein. MSP2N2 produces nanodiscs with a uniform size[10] and can theoretically accommodate more than one full-length $GABA_A$ receptors per disc. However, before nanodisc reconstitution, the receptors were solubilised from cell membranes with detergents. It is important to note that the right choice of detergent is critical when working with $GABA_A$Rs. Previously, it has been shown that solubilization of heteromeric receptors with DDM leads to the collapse of the γ2 subunit in particular, and to the uncoupling of the ECD and TMD regions[11,12]. This limitation was overcome by using LMNG followed by MSP2N2 nanodisc reconstitution, which leads to receptors with pseudo-fivefold symmetry around the pore and with intact contacts between the ECD and TMD [13,14]. Nevertheless, it is unclear whether such receptors are indeed functionally active. To address this question, we undertook a systematic characterization of the full-length α1β3γ2L in various scaffolding environments by limiting the ligand application to 10 seconds.

##### 1.3.1 The impact of the nanodisc sizes

The reconstitution of $GABA_A$ receptors into MSP nanodiscs, which harbour a native-like lipid bilayer, preserves allosteric interactions[13,14]. However, concerns remain regarding the potential limitations posed by their small size on the motions of the TMD, and the possibility that extensive washing during on-bead nanodisc assembly could remove essential lipids[15]. To address these issues, we solved the structures of α1β3γ2 $GABA_A$Rs in MSP2N2 nanodiscs with 130 Å diameter by forming the discs in solution, in the presence of an excess of lipids (Supplementary Fig. 14a-c; Supplementary methods). 1mM GABA was then added to the sample to fully desensitize the receptor. Comparison of the atomic models of the GABA-bound full-length α1β3γ2 receptor's long-lived desensitised state (Extended Data Fig. 6) obtained from nanodiscs formed on-beads *versus* in solution failed to reveal significant differences. Certainly, the respective HOLE pore profiles were nearly indistinguishable, and the Cα-RMSD was < 0.5 Å. The differences between the pore profiles of these structures and the α1β2γ2 $GABA_A$ receptor reconstituted in saposin A discs in complex with GABA (PDB: 6X3Z)[15] might therefore arise from inherent subunit differences (β3 versus β2) as well as sidechain-rotamer choices during modelling (Supplementary Fig. 14b). Furthermore, lipid–protein interactions are clearly preserved (Supplementary Fig. 13c-f). For example, prominent lipid densities are observed at the $\beta3^{+}/\alpha1^{-}$ interface in the long-lived desensitised state. Moreover, PIP2 fully occupies both binding sites in all nanodisc-reconstituted structures of full-length α1-containing receptors (Supplementary Fig. 14f).

##### 1.3.2 The impact of the nanodisc scaffold

Ligands were mixed with α1β2γ2L receptors directly on EM grids, followed by immediate blotting and plunge-freezing, all accomplished within approximately 10 s. We applied the same technique to receptors reconstituted in saposin A or SMALP nanodiscs to determine the potential structural impact of different nanodisc scaffolds.

Employing the 10 s protocol, receptors within saposin A nanodiscs were exposed to 1 mM GABA plus 10 μM etomidate, yielding the structure of the GABA-plus-etomidate-bound long-lived desensitised state. We also solved the resting state of the receptor reconstituted in saposin A nanodiscs. The resting and long-lived desensitised states solved under these conditions exhibited no significant differences when compared to those determined in the MSP nanodiscs. However, in the 10-s application dataset with 1 mM GABA plus 10 μM etomidate, approximately 20% of the particles obtained in saposin A discs contained a collapsed γ2 subunit (Supplementary Fig. 15a-c), an occurrence not observed in MSP nanodiscs. This phenomenon is likely attributable to the saposin A-reconstitution process, as in previous studies receptors assembled in saposin A discs necessitated the exclusion of particles with damaged γ2 subunits[15]. Interestingly, the damage occurs during ligand application. Taken together, saposin A nanodiscs therefore may not be representative of the actual membrane environment and could result in the emergence of non-physiological states.

During nanodisc reconstitution, the receptor must be first solubilised from the membrane by detergents. However, the latter may lead to potential irreversible damage to the protein or to the loss of essential lipids that cannot be recovered upon reconstitution in a lipid nanodisc. To avoid detergent solubilization altogether, we used styrene-maleic acid (SMA) copolymers, which disrupt membranes and extract receptors surrounded by a layer of native cellular lipids[16].

SMA copolymers allow the direct isolation of membrane-embedded receptors in complex with endogenous lipids, in disc-like particles called SMALPs. These lipid discs might therefore be more representative of a physiologically relevant environment in terms of chemical composition[17]. However, it is unknown whether receptors isolated in this way can interconvert among states in a meaningful timescale and whether such conformational changes are consistent with functional data. Recently, the structure of the glycine receptor (GlyR) was solved in SMALPs[18]. The authors of this study reported several distinct states of the receptor bound to the full agonist glycine as well as the partial agonists taurine and GABA. Along with the canonical resting and desensitised states, the authors also presented a putative open and 'expanded-open' state. The authors use the term 'expanded-open' to describe a state whose pore's narrowest constriction has a diameter of ~7 Å, much larger than expected from electrophysiological pore-sizing experiments[19,20]. Interestingly, the open state reported in this study is symmetrical along the pore and the Cα-RMSD between the open and desensitised state in the TMD region is only 0.6 Å. This also contrasts to studies where ligand application on the millisecond timescale on a heteromeric nAChR embedded in its native lipid environment report an asymmetric gating cycle for an otherwise pseudo-symmetric receptor in the resting state[21].

It is important to note that the experiments reported by Yu et al. were performed under time constraints, namely the sample was frozen within 10 s. The authors therefore propose that by using their 'time-resolved' approach one could capture the open channel structure. The states observed indeed have a pore radius large enough to allow a fully hydrated chloride ion to go through. However, it is crucial to discuss if such states are physiologically meaningful. Yu et al. report that in the presence of 10 mM glycine, the GlyR adopts an open state in 55% of particles, a desensitized state in 15% of particles and an expanded-open state in 35% of particles. The fraction in these three states, however, is not consistent with the state occupancies expected from electrophysiological observations, which report that 84.1 ± 11.8% of the receptors would reach the desensitised state when glycine is applied for 9 s[22]. This, together with the fact that the authors report an 'expanded-open' state, raises the question as to whether any of these structures resemble the actual open states of the GlyRs in native membranes. Moreover, the authors of the study also report that on the same timescale, all receptors are desensitised when reconstituted into MSP2N2 nanodiscs, which is what is expected on the basis of electrophysiology data[23].

To ascertain whether SMALP nanodiscs allow a structural analysis of $GABA_A$ receptor gating rearrangements, we solubilized the full-length α1β3γ2L assembly using the protocol used by Yu and coworkers[18] for the α1-GlyR. The $GABA_A$ receptor was supplemented with 1 mM GABA and 10 μM etomidate for 10 s. As expected, most of the particles had densities for GABA bound below loop C at the $\beta3^+/\alpha1^-$ interfaces. However, the TMD region was dramatically distorted. Unlike the uncoupled state in Saposin A, in SMALPs, the density for the γ2 subunit's TMD is entirely fragmented and no longer makes contacts with the TMDs of the neighbouring subunits. Moreover, the interface between the α1-A and β3-B subunits is also broken and no density for etomidate can be observed (Supplementary Fig. 15d-g).

The cause for this dramatic collapse remains unclear. One possible explanation is that the SMA co-polymers cannot provide enough lateral support to the disc leading to a decreased rigidity relative to native membranes and MSP2N2 nanodiscs. It is, therefore, evident that the receptor cannot undergo physiological gating motions in SMALPs because it cannot reach the 'canonical' long-lived desensitised state. Therefore, pLGICs conformations in SMALPs should be considered with caution.

On the other hand, the resting state in SMALPs is intact, with coupled ECD and TMD in all subunits, suggesting that the collapse may only occur in the presence of agonist. The structure of the resting state in SMALPs sheds light on the native lipid environment of the receptor. Compared to that in the resting state of MSP2N2-disc-reconstituted $GABA_A$Rs, no significant differences regarding lipid densities can be observed beyond $PIP_2$ (Fig.2; Supplementary Fig. 15c-d). Since MSP-reconstituted receptors show a similar lipid profile (Supplementary Fig. 14c-f), it appears that essential lipids remain associated with the receptors during detergent solubilization or are re-supplemented by the addition of BBE during nanodisc formation. We further show that receptors reconstituted in SMALPs have variable $PIP_2$ occupancy, which seems to be critical for the gating of the receptor as $PIP_2$-bound receptors failed to undergo opening (Fig. 2, Fig. 3). Such differences in $PIP_2$ occupancy could not be observed for the MSP2N2-reconstituted receptor (Supplementary Fig. 14). This likely stems from the addition of bovine brain extract (BBE) during nanodisc formation, which can provide additional $PIP_2$ molecules and lead to the full occupancy of both binding sites. The addition of the BBE is however crucial as it supplements essential lipids, such as cholesterol and POPG, that are required for channel gating (Fig. 3, Fig. 4).

### 1.4 The cholesterol content of the nanodisc

Synaptic $GABA_A$Rs that contain gamma2 subunits associate with protein partners, such as GARLH4 and Neuroligin-2[24-26], that "capture" receptors in close proximity to neurotransmitter release sites[27,28]. Many other mechanisms modulate the location, trafficking and signalling of $GABA_A$Rs in a wider cellular context[29]. It has been established that synaptic membranes are particularly enriched in cholesterol and sphingomyelin[30], which form microdomains with high rigidity, akin to lipid rafts. These are likely to further impact the distribution of receptors at the cell surface[31]. Interestingly, it has been shown that depleting cholesterol from cell membranes can negatively affect the modulation of $GABA_A$ receptors by diazepam[32]. Furthermore, cholesterol has been shown to play a key role in signalling and localization of the *Torpedo* nAChR[33]. We therefore tested the effect of altering the cholesterol level on the $GABA_A$ receptor by artificially adding extra cholesterol to the disc. This was done by incubating the nanodiscs with soluble MβCD-cholesterol complexes, a strategy that has been extensively used to enrich the level of cholesterol in cellular membranes[34].

To calculate the amount of cholesterol in a nanodisc:

1) The molecular weight of cholesterol is $M.W_{(cholesterol)} = 386.7$g/mol
2) The molecular weight of the $GABA_A$ receptor-nanodisc complex is $M.W_{(receptor)} = 500\,000$ g/mol.
3) In Supplementary Fig. 5 the amount of $GABA_A$ receptor-nanodisc complex used for measurement was $A_{280nm} = 0.8$ mg/mL. 20µL samples was used meaning a total of 16µg receptor was present.
4) We can thus calculate that $n_{(receptor)} = mass_{(receptor)}/M.W_{(receptor)} = 3.2\times10^{-11}$. moles. In the case of cholesterol, if 1µg cholesterol is detected in the sample then:

   $$n_{(cholesterol)} = mass_{(cholesterol)}/\ M.W_{(cholesterol)} = 260\times10^{-11} \text{ moles.}$$
5) This means that upon detection of 1µg of cholesterol, we would have $n_{(cholesterol)}/n_{(receptor)}$ ~ 80 cholesterol molecules per receptor-nanodisc complex.

As shown in Supplementary Fig. 16, 'standard' MSP discs contain only small amounts of cholesterol (~0.1 µg), which translates to ~8 molecules of cholesterol per disc. The amount of cholesterol in the standard preparation, however, does not mimic the synaptic cholesterol concentration and is therefore, likely representative of the extrasynaptic environment[35]. To raise the cholesterol level of our nanodiscs to values typically found in synaptic membranes, we incubated them with a high concentration of cholesterol in the form of soluble complexes with methyl-β-cyclodextrin (MβCD). We found that the added cholesterol became incorporated into the nanodiscs only when this incubation step took place during the formation of the nanodisc, but not if already formed nanodiscs were incubated with the MβCD−cholesterol complexes. When 6.6 mM MβCD−cholesterol was added, ~1μg of cholesterol was detected in the sample which is a 10-fold increase when compared to the 'standard' procedure and translates to ~80 cholesterol molecules per disc. Importantly, the cholesterol content becomes saturated as incubations with complexes at a concentration of 13 mM failed to further increase the level of cholesterol of the nanodiscs (Supplementary Fig. 16).

It has been estimated that the number of dipalmitoylphosphatidylcholine (DPPC) lipid molecules in an empty MSP2N2 disc is ~300[36]. In the case where a membrane protein is inserted, the amount of lipid will decrease. The volume of the $GABA_A$ receptor's TMD is $4.5\times10^4$ $Å^3$ (calculated with the volume function in ChimeraX[37]). The volume of the nanodisc can be calculated readily since it is an ellipsoid with axes with dimensions roughly A=B=45Å, C=19 Å. The volume of the ellipsoid is thus $4/3\times\pi\times A\times B\times C = 16\times10^4$ $Å^3$. The $GABA_A$ receptor therefore takes up roughly 1/3 of the nanodisc volume, therefore leaving space for ~200 lipids. Since we have 80 molecules of cholesterol in the disc and a total of 200 lipids, the concentration of cholesterol is roughly 40% of the total lipids which is similar to the amount measured in the mammalian synaptic lipidome[30].

### 1.5 The lipid insulation of the open states

During ligand application on the millisecond timescale, $GABA_A$ receptors transition through short-lived asymmetric bound-closed, open, and desensitised states. This is true for all receptor assemblies examined in this study, including β3 homomers (Fig. 1), α1β3 diheteromers (Fig. 3), and α1β3γ2 triheteromers (Fig. 4). We further demonstrate that lipids can enter the pore through the inter-subunit space (ISS), blocking chloride ion entry during the short-lived desensitised states and some bound-closed states. Interestingly, in the open states, we observe significant asymmetry without lipid obstruction. This is likely due to differences in the ISS between the short-lived open and desensitised states.

For lipids to enter the pore, the ISS must be large enough to accommodate an acyl chain. Considering the van der Waals (VDW) radius, the acyl chain would have a semi-minor VDW radius between that of methane (1.88 Å) and ethane (2.95 Å)[38]. Therefore, the gap must be able to fit an ethane molecule.

To experimentally determine the ISS radius that allows lipid entry without completely blocking the pore, we focused on the symmetric desensitised state of α1β3γ2. In this state, POPG is bound at the $\beta3^{+}/\alpha1^{-}$ interface, with one acyl chain protruding into the ISS but not extending past the M2 helix to block the pore (Supplementary Fig. 17a-b). This state exemplifies the ISS radius needed to permit lipid entry while preventing complete pore blockage. Using HOLE analysis (with start point for the calculation in the middle of the lipid bilayer just next to the outer side of the M1/M3 helices and end point in the middle of the pore), we found that the minimum radius of the cavity at the POPG binding site is 2.6 Å[39]. When the ISS radius decreases to 2 Å or less, lipid acyl chains can no longer protrude, insulating the pore from lipid entry (Supplementary Fig. 17c-e). This analysis was extended to all open states reported in this study to verify pore insulation from lipids (Supplementary Fig. 10). In the EM maps, no lipid density was observed occluding the pore in any open state. As shown in Supplementary Fig. 10, all open states, except for the α1β3 branch 1 open state, have a minimum ISS radius of 2 Å or less, indicating insulation from lipids. The α1β3 open branch1 state maintains an ISS radius of approximately 3 Å, suggesting that lipid tails may enter the ISS and reach the pore. However, the lack of clear lipid density in the pore (Fig. 3b) suggests that any lipid block that may take place in this particular open state is unfavourable.

### 2. Supplementary Materials and Methods

**2.1 Saposin A purification**

The pET45 vector containing the Saposin A gene was purchased from Addgene and was transformed into Rosetta(DE3) *E.coli* cells (Sigma). 3 L of cells were grown in TB media to $OD_{600nm}$=1, and the expression of Saposin A gene was induced with 1mM IPTG for 4 h at 28°C. Cells were collected by centrifugation and the pellets were resuspended in 20mM HEPES, 150mM NaCl pH=7.6 supplemented with cOmplete protease inhibitor tablet. Next, the cells were homogenized with a cell disruptor at a pressure of 38,000 psi, and the lysate was clarified at 30,000×g for 30 min at 4°C. The supernatant was collected and heated to 85°C for 10mins and centrifuged again at 30,000×g for 30 min at 4°C. The supernatant was applied on a HisTrap™ High Performance (Cytiva) 5mL column, washed and eluted with the following buffers:

1) Wash 1: 15CVs of 20mM imidazole, 20mM HEPES, 150mM NaCl pH=7.6
2) Wash 2: 15CVs of 40mM imidazole, 20mM HEPES, 150mM NaCl pH=7.6
3) Elution: 5CVs of 500mM imidazole, 20mM HEPES, 150mM NaCl pH=7.6

The eluate was dialyzed overnight in SnakeSkin 3.5kDa MW cutoff dialysis tubing (Thermo Fisher Scientific) against buffer containing 20mM HEPES, 150mM NaCl pH=7.6. 3C protease was added to Saposin A in a 1:100 molar ratio to cleave the $His_6$-tag during the dialysis step. The sample was then purified with reverse nickel chromatography. The flowthrough was collected, concentrated to 9mg/mL, snap frozen in $LN_2$ and stored at -80°C until further use.

**2.2 $GABA_A$R purification and reconstitution into Saposin A nanodiscs**

The procedure for nanodisc reconstitution of the full-length α1β3γ2L was the same as previously described[13,14] with the difference that the MSP2N2 scaffolding protein was replaced with Saposin A.

**2.4 Cholesterol enrichment of MSP2N2 nanodiscs**

Cell pellets of the full-length α1β3γ2L were resuspended in PBS pH=7.4 supplemented with cOmplete protease inhibitor tablet (Roche). For nanodisc reconstitution, a 10% (w/v) solution of lauryl maltose neopentyl glycol (LMNG, Anatrace) was prepared in water and added at 1% (w/v) to the cell suspension. Cells were solubilized for 1h at 4°C, and cell debris and other insoluble material were removed by centrifugation (15,000×g, 15 min). The supernatant from

1L of cells was incubated with 300μL of 1D4 affinity resin and gently mixed at 4°C for approximately 2 h.

The resin was recovered by centrifugation at 300×g, 4°C and washed 3 times with 50mL PBS pH=7.4 containing 0.1% (v/v) LMNG. The beads were then collected by centrifugation at 300×g, 4°C and incubated for 30min with 450μg of phosphatidylcholine (POPC, Avanti) and bovine brain lipid (BBE) extract (type I, Folch fraction I, Sigma-Aldrich) mixture (POPC:BBE = 70:30 w/w). For the cholesterol enrichment experiments performed during disc formation, the desired concentration (1.6mM to 13mM) of soluble cholesterol in the form of MβCD-cholesterol (Sigma) was added together with the lipids. 100μL MSP2N2 (5mg/mL) was then added to the beads and further incubated for 30min. Next, Bio-beads SM-2 (Bio-Rad) equilibrated in PBS were incubated with the resin for 90 minutes. The beads were then washed 6 times with 1mL of PBS pH=7.4 and eluted overnight with 5mM 1D4 peptide, PBS pH=7.4. When cholesterol enrichment was performed after disc formation, 4mM MβCD-cholesterol was added after the washes and further incubated for 2h followed by six more washes and overnight elution.

**2.5 Cholesterol content measurement**

To quantify the cholesterol content of the nanodiscs used for $GABA_AR$ reconstitution, 20μL samples at $A_{280nm}$=0.8mg/mL were mixed with 200μl ice-cold MeOH, vortexed, and sonicated in an ice bath for 15 min. Then 400μL chloroform were added, and the sample was again vortexed and sonicated for 15 min. Next, 150μL ice-cold water was added, followed by vortexing and sonication for 15 min. The sample was then spun at 16,000×g for 15 min and the organic layer (lower layer containing the lipids) was carefully removed. Another 400μL of chloroform was added to re-extract the lipids, followed by vortexing, sonication and a second 16,000×g, 15 min centrifugation. The 800μL sample was evaporated in a vacuum desiccator at 45°C. The cholesterol concentration was measured fluorometrically using the Sigma cholesterol assay kit (CS0005) following the manufacturer protocol. Samples were measured in triplicate.

**2.3 Assembly of large MSP2N2 nanodiscs**

To reconstitute the full-length α1β3γ2L $GABA_AR$ into larger nanodiscs, cell pellets were resuspended in PBS pH=7.4 supplemented with cOmplete protease inhibitor tablet (Roche).

For nanodisc reconstitution, a 10% (w/v) solution of lauryl maltose neopentyl glycol (LMNG, Anatrace) was prepared in water and added at 1% (w/v) to the cell suspension. Cells were solubilized for 1h at 4°C, and cell debris and other insoluble material were removed by centrifugation (15,000×g, 15 min). The supernatant from 1L of cells was incubated with 300µL FLAG M2 monoclonal antibody affinity gel (Sigma), incubated for 2h at 4°C with constant rotation. The resin was recovered by centrifugation at 300×g, 4°C and washed three times with 50mL PBS pH=7.4 containing 0.1% (v/v) LMNG. The protein was eluted with 100µM FLAG peptide, PBS pH=7.4. The eluted protein was mixed with MSP2N2 (5 mg/ml), and POPC:BBL (70:30 w/w) stock in molar ratio of 1:1:100, respectively and incubated for 30 min. Next, Bio-beads SM-2 (Bio-Rad) equilibrated in PBS were incubated with the resin for 90 min. 300µL 1D4 beads were then added for a second purification step aiming to remove the empty nanodiscs. After 2h incubation at 4°C with constant rotation, the beads were washed six times with 1mL of PBS pH=7.4 and eluted overnight at 4°C with 5mM 1D4 peptide, PBS pH=7.4.

##### 2.4 Cryo-EM grid preparation

Prior to vitrification on EM grids, purified α1β3γ2L receptor samples were pre-incubated with 2µM Mb38. For the resting state in Saposin A, no Mb38 was added. 3.5 µL of the sample were applied to a freshly glow-discharged (PELCO easiGlow, 30 mA for 120s) gold R1.2/1.3 or R0.6/1 300 mesh UltraAuFoil grid (Quantifoil) and blotted for 3.5s before plunge-freezing in liquid ethane using a Leica EM GP2 plunger (Leica Microsystems; 95% humidity, 14 °C). The following ligands were used for each dataset: for the Saposin A dataset, 1 µM GABA plus 10 µM etomidate were used, and for the SMALP dataset, 1mM GABA plus 10 µM etomidate were used. For the 10s ligand-application approach, the ligands were mixed directly with the protein on the grid.

For the time-resolved truncated-α1β3 dataset, 3µM Mb25 was added to the sample. 3.5 µL of sample was applied to glow-discharged Quantifoil 1.2/1.3 (PELCO easiGlow, 30 mA for 60 s) and manually plunged at ~8 °C and ~95% humidity. Instead of ligand, PBS pH=7.4 was sprayed as a negative control. In the case of the β3 homomer, no fiducial nano/megabody was added and the buffer sprayed was PBS pH=8. The total PBS exposure time was limited to ~10ms by adjusting the spraying nozzle to 5mmm above the surface of the liquid ethane. The grid was released by free fall 35mm above the ethane level and spraying was initiated by an optical trigger.

#### 2.5 Calculation of pore radii and conductance in open states

To accurately quantify the radii of the asymmetric open states, we used PoreAnalyzer[40], a tool designed to compute pore dimensions by fitting ellipsoids to asymmetric channels. Since the pores are not symmetric, a simple circular cross-section cannot accurately capture their shape. Instead, the more flexible geometry of an ellipsoid, defined by two radii—the semi-minor and semi-major axes—is used to represent the cross-sectional asymmetry. PoreAnalyzer reports both of these radii, along with an estimated conductance for each state. The conductance is calculated by modelling the pore as a series of ellipsoidal segments. For each segment, the local resistance is derived from its geometry, and these are summed along the pore axis to yield a total resistance. Conductance is then obtained using Ohm's law. To improve accuracy, the underlying conductivity function has been benchmarked against molecular dynamics simulations of ion permeation through pores of varying shape and asymmetry. This calibration allows the model to better capture the effects of geometric irregularities on ion flow.

Unlike HOLE as implemented in Coot, where users can manually define the start and end points for pore calculation, PoreAnalyzer automatically selects the permeation pathways. However, in our structures, multiple tunnels with distinct geometries were present, which complicated automated selection. To ensure that the correct tunnel, i.e. the one with the largest diameter, was chosen, we introduced computational mutations to occlude the narrower, off-path tunnels. Specifically, we mutated pore-lining residues forming the smaller tunnels to bulkier residues in Coot, effectively blocking their selection during analysis. The following *in silico* mutations were applied: for α1β3 branch1 open, β3-C A252F; for α1β3 branch2 open, β3-B I255R; for α1β3γ2 open1, α1-A P253W; for α1β3γ2 open 2, α1-A I271F.

### 3. Supplementary Figures and Tables

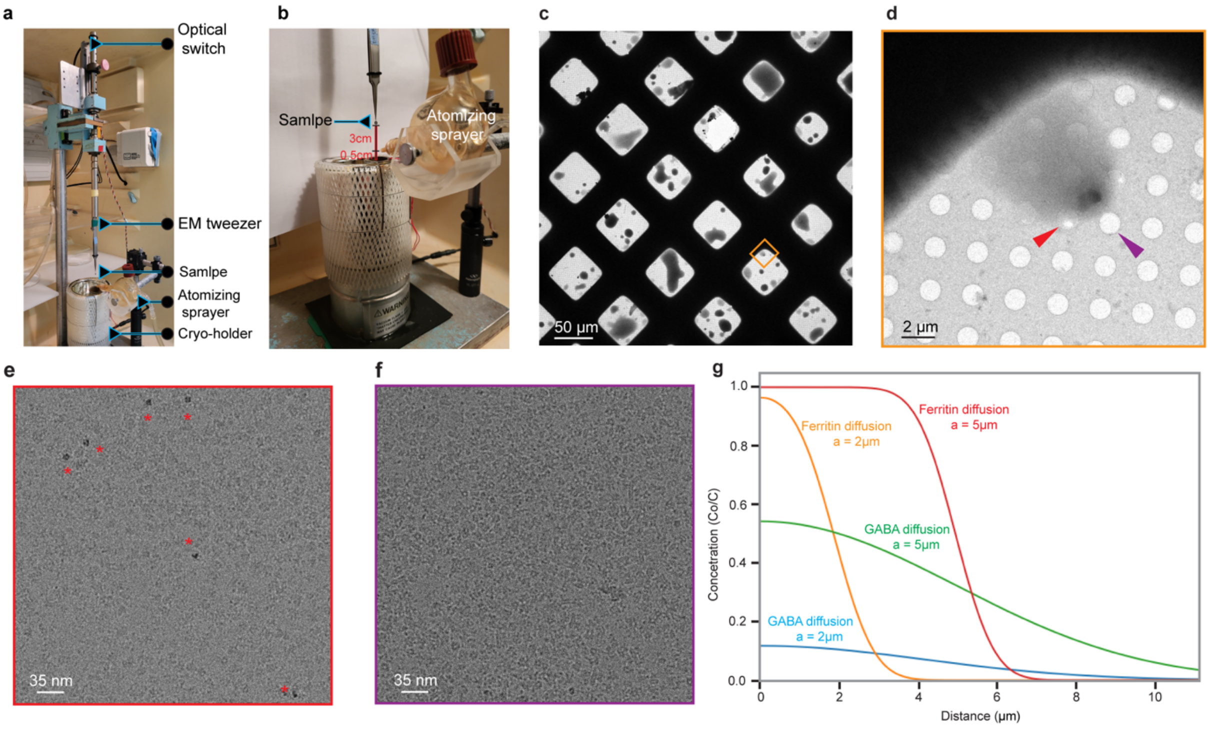

**Supplementary Figure 1 | Time-resolved cryo-EM setup and sprayed ligand diffusion on the millisecond timescale.**

**a,** A tweezer holding the grid is positioned 35 mm above the surface of a liquid ethane cup. **b,** An atomising sprayer, filled with 1mL ligand solutions, is positioned 5 mm above the ethane surface. This limits the ligand exposure 10ms maximum, prior to vitrification. **c**, Representative image of a grid after ligand spraying, shown at atlas magnification (81×). The ligand droplets are seen as dark spots. **d**, Higher magnification image of the cropped grid square region marked with an orange outline in **(c)**. **e**, Image of the grid hole indicated by a red arrowhead in **(d)**, shown at the magnification used for data acquisition (96,000×) and a nominal defocus of -2.8 µm. The electron-dense objects marked with red asterisks are ferritin molecules, used as spraying fiducials, which have a substantially higher contrast than the surrounding $GABA_A$ receptor particles. **f**, Image of the grid hole indicated by a purple arrowhead in (d), shown at the magnification used for data acquisition (96,000×) and a nominal defocus of -2.8 µm. This image has a better contrast than **(e)** as the ice is thinner. **g**, Diffusion curves for ferritin and GABA, calculated for droplet radii (a) of 2 µm and 5 µm, respectively, using the equation described in the Supplementary discussion section 1.1.

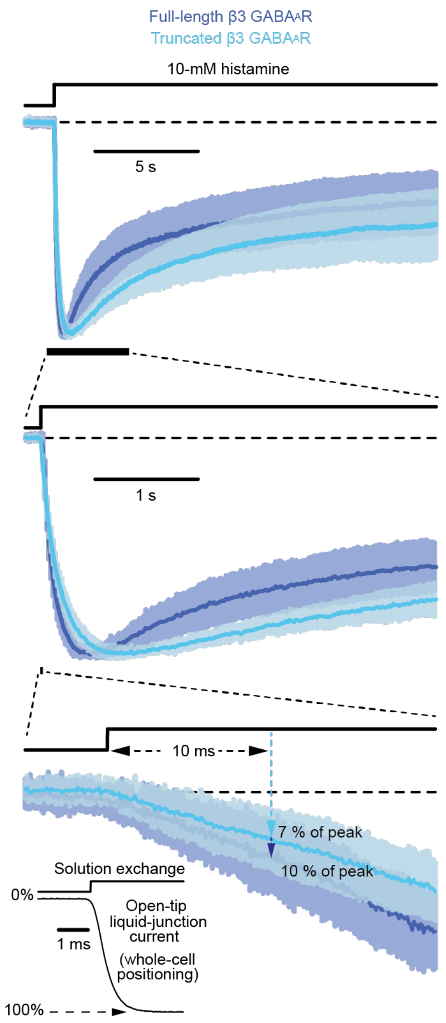

**Supplementary Figure 2 | Electrophysiological responses of full-length and truncated β3 $GABA_A$Rs to the fast application of histamine.**

Inward currents recorded in the whole-cell configuration are displayed at different time scales. Responses are shown as the mean ± one standard deviation (SD) of normalized currents recorded from different cells. Averaged data from the wild-type, full-length β3 GABAAR homomer correspond to a total of 11 responses recorded from 11 different cells, and those from the truncated β3 construct to a total of 12 responses from 12 different cells. The applied potential was ~ –60 mV. Black dashed lines denote the zero-current baseline. The inset of the bottom panel illustrates the speed of solution exchange in these whole-cell experiments (t10–90% = 0.9 ms) as estimated upon switching the concentration of external KCl at the tip of an open patch-clamp pipette.

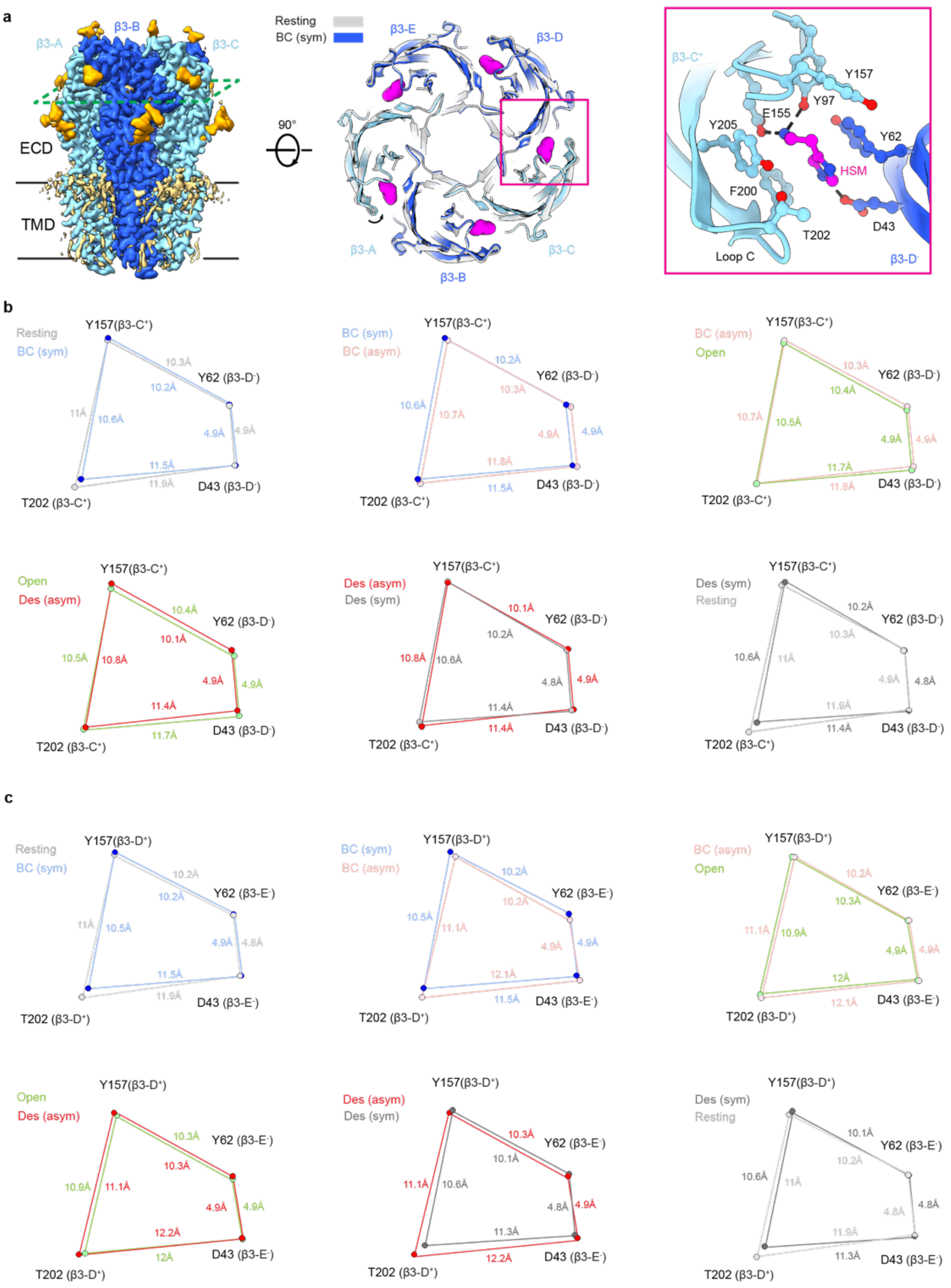

**Supplementary Figure 3 | Conformational changes in the neurotransmitter binding site during activation of the β3 homomeric GABA$_A$ receptor.**

**a,** Cryo-EM map of the β3 homomer viewed parallel to the membrane highlighting the plane of the HSM binding sites (left). View of the ECD cross-sectioned at the level of the HSM binding pocket showing the resting state in grey and the bound-closed symmetric state coloured (middle). One HSM-binding pocket viewed from the extracellular space. The amino acid side chains lining the binding site are shown as balls and sticks and HSM is coloured in magenta. Dashed lines indicate hydrogen bonds (right). **b,** Schematic diagram illustrating the changes in distances (Å) of key residues of the β3-C$^+$/β3-D$^-$ neurotransmitter binding site in the resting, bound-closed (symmetric), bound-closed (asymmetric), open, desensitised (asymmetric) and long-lived desensitised states. **c,** Schematic diagram illustrating the changes in distances (Å) of key residues of the β3-D$^+$/β3-E$^-$ neurotransmitter binding site in the resting, bound-closed (symmetric), bound-closed (asymmetric), open, desensitised (asymmetric) and long-lived desensitised states.

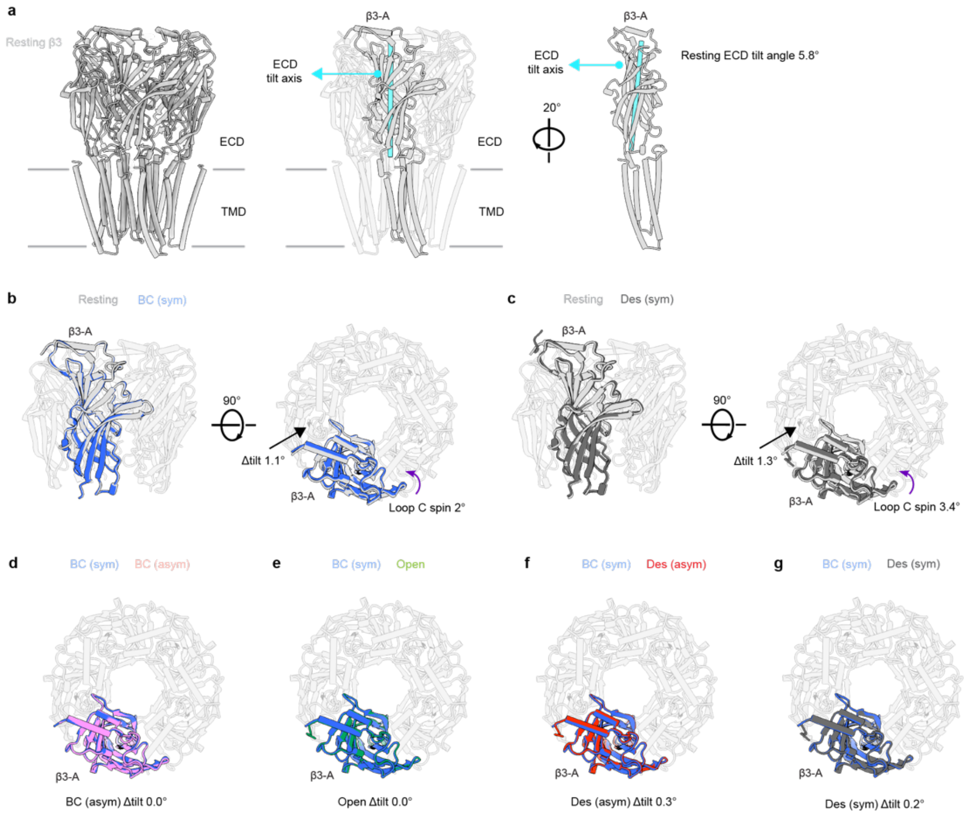

**Supplementary Figure 4 | Tertiary structural rearrangements in the extracellular domain during activation of the β3 homomeric $GABA_A$ receptor.**

**a,** Side view of the β3 homomer resting state model viewed parallel to the membrane plane (left). Subunit β3-A highlighted in grey with the rest of the subunits transparent. The β3-A tilt axis is shown in cyan. The tilt axis was defined by two points: the centre of mass (COM) of residues 91, 93 and 125 in the upper part of the ECD and the COM of residues 46, 59, 151 and 211 in the lower part of the ECD (middle). The β3-A subunit is highlighted and rotated by 20° (right). The tilt of the ECD was calculated as the dihedral angle between the COM of residues 91, 93 and 125 of β3-A, the COM of the ECDs of all five subunits (residues 1 to 218), the COM of the TMDs of all five subunits (residues 218 to 448) and the COM of residues 46, 59, 151 and 211 of β3-A. **b,** Side view of the ECD portion of the β3 homomer comparing the resting and bound-closed symmetric states. The tilt of the bound-closed symmetric state was calculated with respect to the resting state's whole ECD and whole TMD COMs while the difference in tilt (Δ tilt) was calculate as the difference in dihedral angles of the bound-closed symmetric state and the resting state. Loop C spin was calculated as the angle between residue β3-A T202 Cα (tip of loop C) in the resting state, the COM of the resting β3-A ECD and the β3-A T202

Cα of the bound-closed state. **c,** Comparison in loop C rotation and tilt between the long-lived desensitised and resting states. Comparison of Δ tilt between the bound-closed symmetric state

Supplementary Figure 13 continued

and **d,** bound-closed asymmetric state; **e,** open state; **f,** desensitised asymmetric state and **g,** long-lived desensitised state.

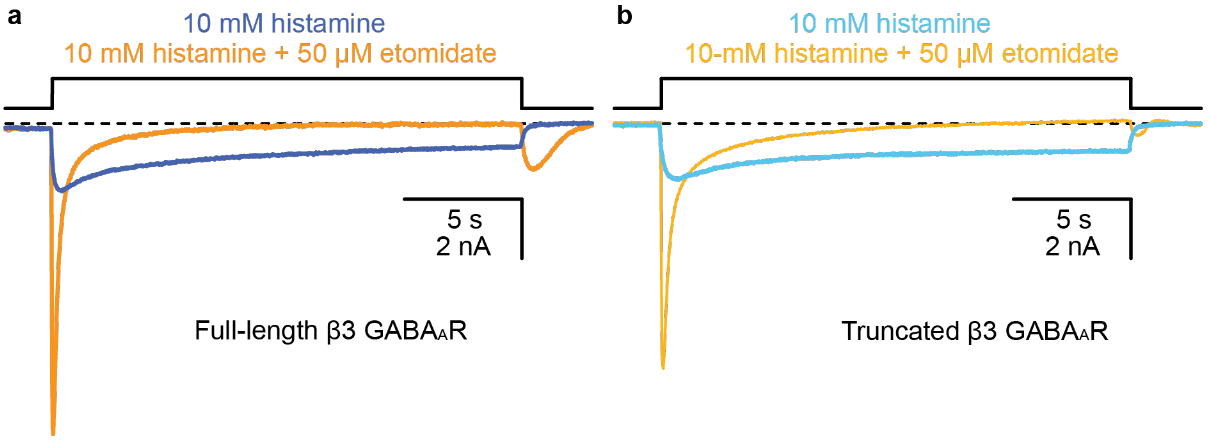

**Supplementary Figure 5 | Comparison of histamine- and histamine + etomidate-gated currents on the full-length and truncated β3 GABA$_A$Rs.**

Whole-cell inward currents recorded from the full-length and truncated β3 GABA$_A$R homomers. The responses shown are representative examples of currents recorded from different cells. **a,** The average peak currents recorded from the full-length channel were 2.7 ± 2.4 nA (*n* = 12; 10 mM histamine) and 11 ± 3.6 nA (*n* = 9; 10 mM histamine + 50 µM etomidate). **b,** The average peak currents recorded from the truncated β3 construct were 2.5 ± 2.1 nA (*n* = 14; 10 mM histamine) and 9.9 ± 4.6 nA (*n* = 8; 10 mM histamine + 50 µM etomidate). Upon washout of the mixture of histamine and etomidate, a “rebound” current[8] was observed for both constructs. The applied potential was ~ –60 mV. Black dashed lines denote the zero-current baseline.

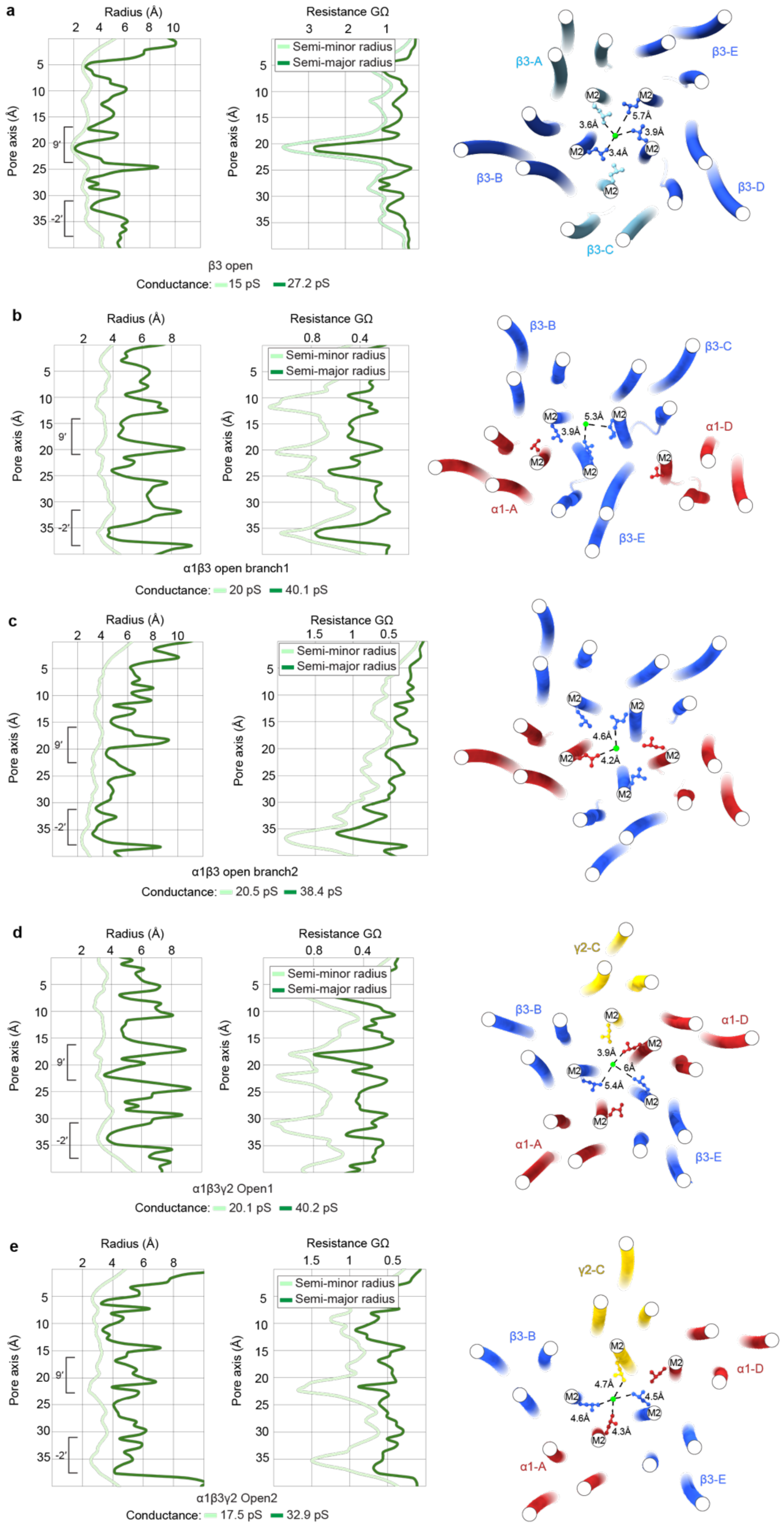

**Supplementary Figure 6 | Pore radii and estimated conductance of GABA$_A$ receptor open states.**

**Supplementary Figure 6 continued**

The semi-minor radius, semi-major radius and the conductance of the open states was analysed with the PoreAnalyzer package[40]. The plots of radii as determined by HOLE[39] (semi-minor radius) and PoreAnalyzer (semi-major radius) are shown on the left panel. The resistance estimation of the state is shown in the middle panel alongside the calculated conductance value underneath in pS for both the semi-minor radius (determined by HOLE) and the semi-major radius (determined by PoreAnalyzer). The experimentally determined conductance of the β3 homomer homologue, the β1 homomer, is 18 pS[41]. The experimentally measured main conductance state of α1β3 is 19 pS and for α1β3γ2 the main conductance state is 28 pS[42]. The right panel represents a cross-section of the TMD as seen from the extracellular space above the level of the 9′ activation gate. A placeholder $Cl^-$ pseudo atom (shown in lime) was placed in Coot to serve the role of a marker to measure the distances from the 9′ Leu activation gate for the **a,** β3 homomeric open state; **b,** α1β3 branch1 open state; **c,** α1β3 branch2 open state; **d,** α1β3γ2 open1 and **e,** α1β3γ2 open2.

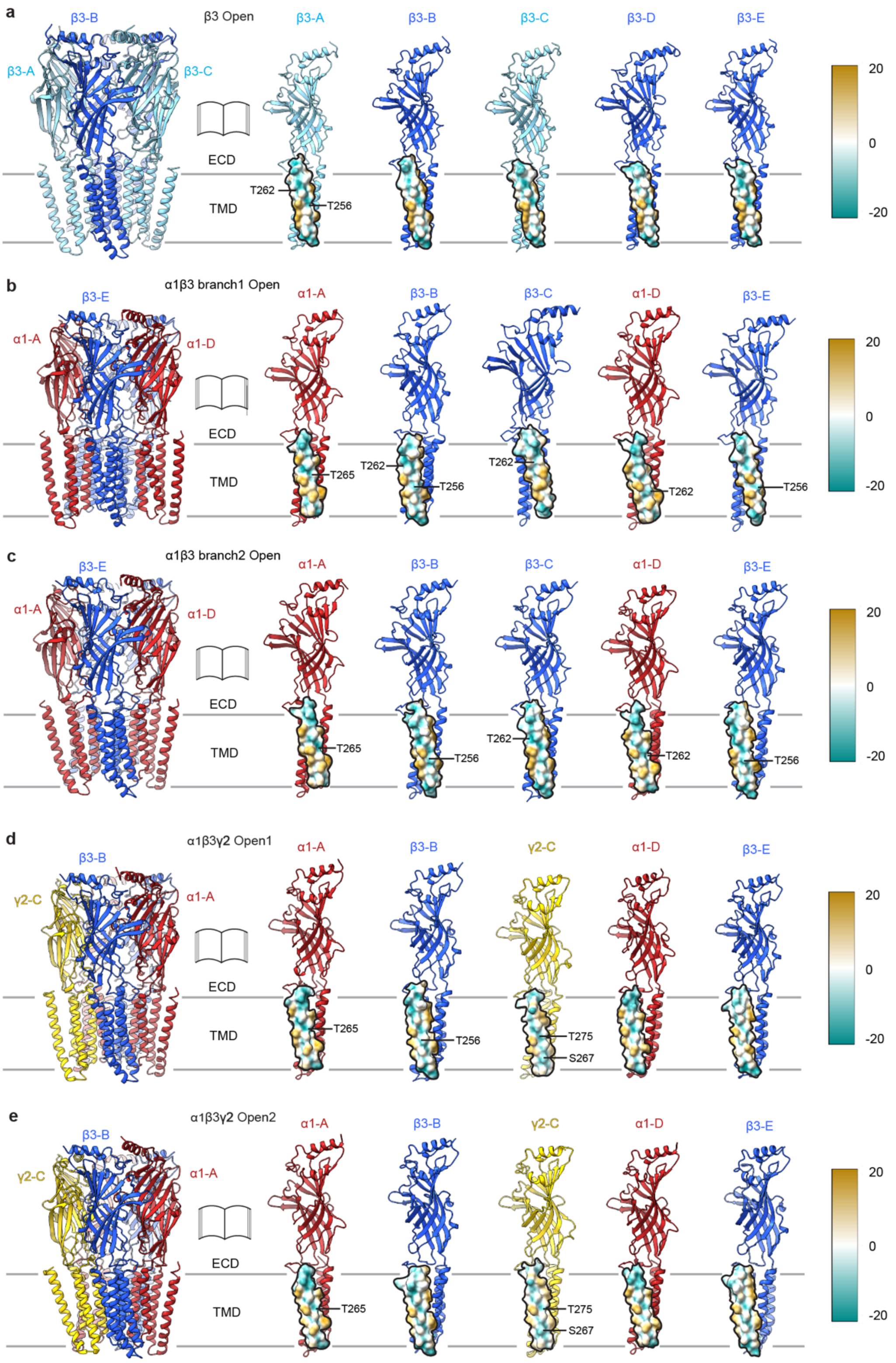

**Supplementary Figure 7 | Hydrophilicity of the pore-lining M2 Helices in $GABA_A$ Receptor Open States.**

Side views of the open states are shown with each subunit shown individually for the **a,** β3 homomeric open state; **b,** α1β3 branch1 open state; **c,** α1β3 branch2 open state; **d,** α1β3γ2 open1 and **e,** α1β3γ2 open2. The M2 helices were coloured by hydrophobicity in ChimeraX – cyan for most hydrophilic and wheat for most hydrophobic. For each subunit, the side of the M2 helix shown is facing the pore of the receptor. The pore lining residues that could participate in the dehydration of $Cl^-$ ions are highlighted. Mutations in the following residues have been shown to cause epilepsy indicating impaired GABAergic signalling: α1Thr262Ala[43], α1Thr265Ile[44], β3Thr256Ala[45], β3Thr262Ile[46] and γ2Ser267Phe[47].

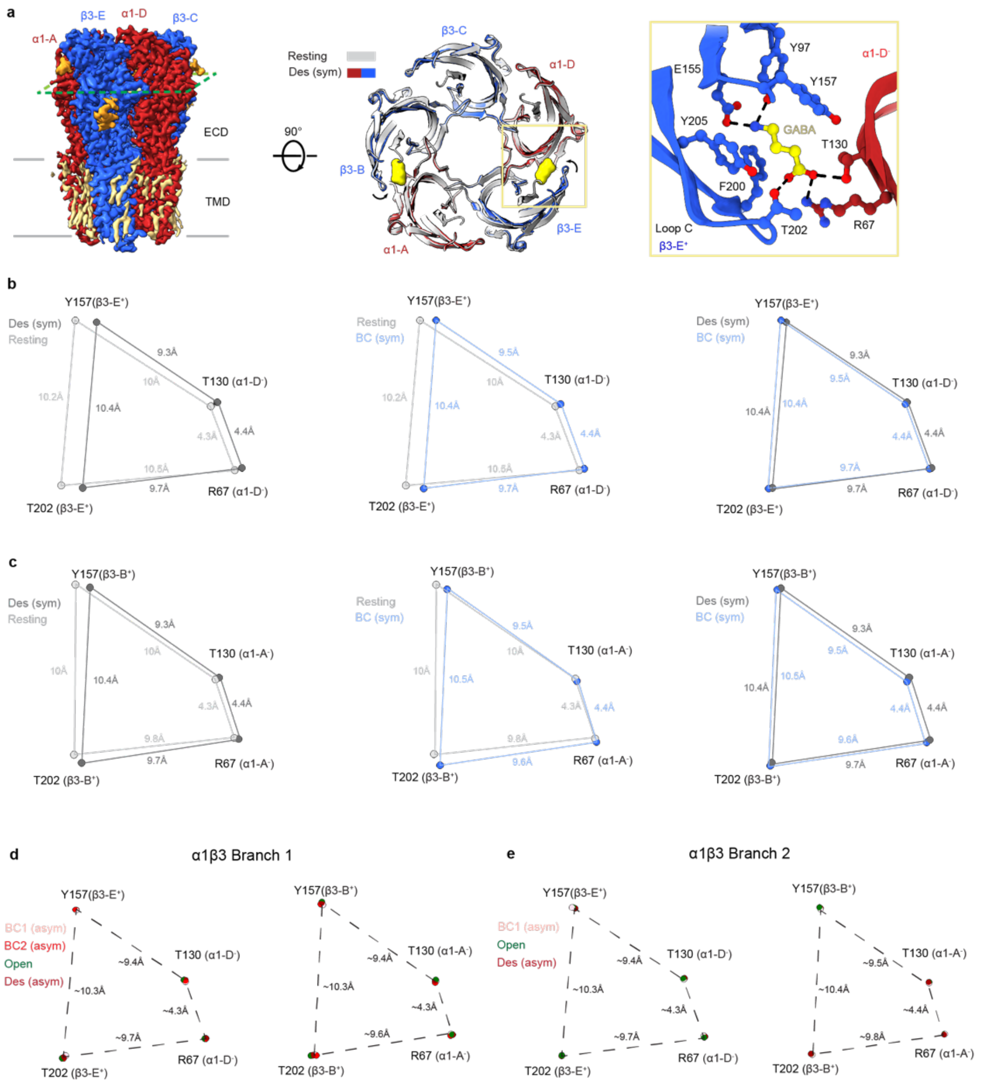

**Supplementary Figure 8 | Conformational changes in the neurotransmitter binding site during activation of the α1β3 diheteromeric GABA$_A$ receptor.**

**a,** Cryo-EM map of the α1β3 diheteromer viewed parallel to the membrane highlighting the plane of the GABA binding sites (left). View of the ECD cross-sectioned at the level of the GABA binding pocket showing the resting state in grey and the bound-closed symmetric state coloured (middle). One GABA-binding pocket viewed from the extracellular space. The amino acid side chains lining the binding site are shown as balls and sticks and GABA is coloured in yellow. Dashed lines indicate hydrogen bonds (right). **b,** Schematic diagram illustrating the changes in distances (Å) of key residues of the β3-E$^+$/α1-D$^-$ neurotransmitter binding site in the resting, bound-closed (symmetric) and long-lived desensitised states. **c,** Schematic diagram illustrating the changes in distances (Å) of key residues of the β3-B$^+$/α1-A$^-$ neurotransmitter binding site in the resting, bound-closed (symmetric) and long-lived desensitised states. **d,**

Schematic diagram illustrating the changes in distances (Å) of key residues of the β3-E$^{+}$/α1-D$^{-}$ (left) and β3-B$^{+}$/α1-A$^{-}$ (right) neurotransmitter binding site in the bound-closed 1 (asymmetric), bound-closed 2 (asymmetric), open and desensitised (asymmetric) states of branch 1. The distances remain largely the same. **e,** Schematic diagram illustrating the changes in distances (Å) of key residues of the β3-E$^{+}$/α1-D$^{-}$ (left) and β3-B$^{+}$/α1-A$^{-}$ (right) neurotransmitter binding site in the bound-closed 1 (asymmetric), open and desensitised (asymmetric) states of branch 2. The distances remain largely the same.

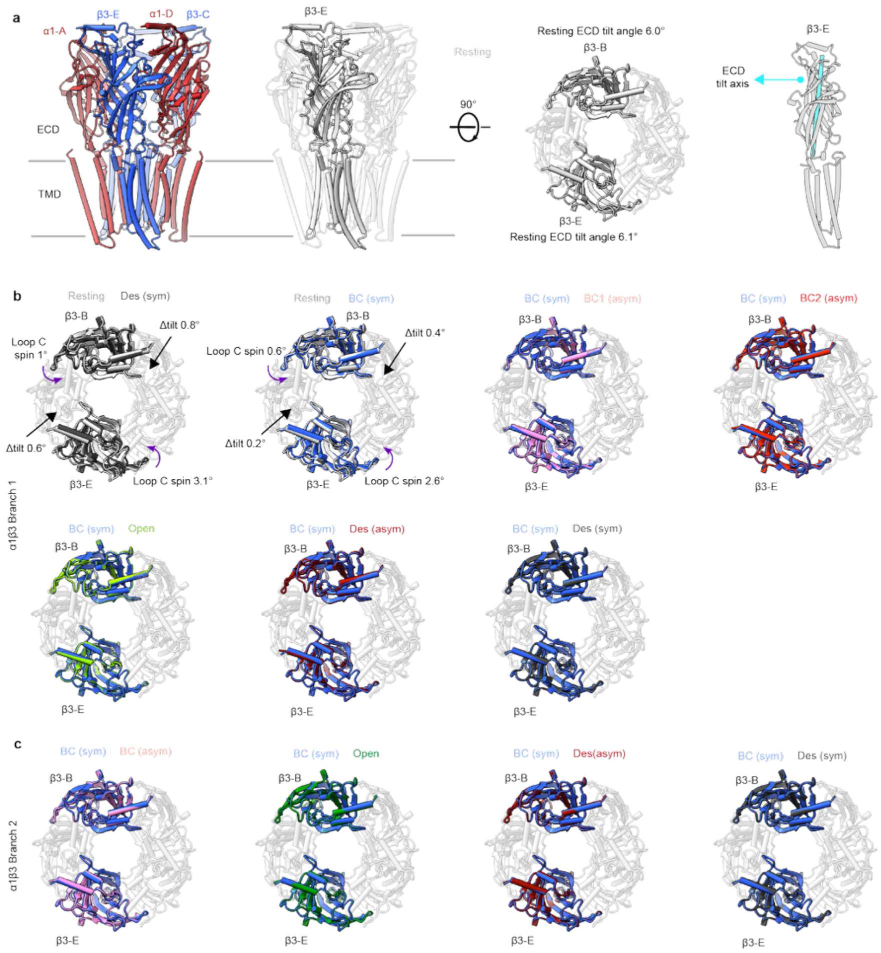

**Supplementary Figure 9 | Tertiary structural rearrangements in the extracellular domain during activation of the α1β3 diheteromeric $GABA_A$ receptor.**

**a,** Side view of the α1β3 diheteromer resting state model viewed parallel to the membrane plane (left). Subunits β3-B and β3-E highlighted in grey with the rest of the subunits transparent (middle). The β3-E is highlighted, and the tilt axis is shown in cyan (right). The tilt angle was calculated as in supplementary figure 13. **b-c,** The Δ tilt and loop C rotation calculations were performed as described in supplementary figure 13. The comparison is between the right and left state labels on top of each ECD as viewed from the extracellular space. No significant differences can be observed in the ECD tilt or loop C rotation for the agonist-bound states.

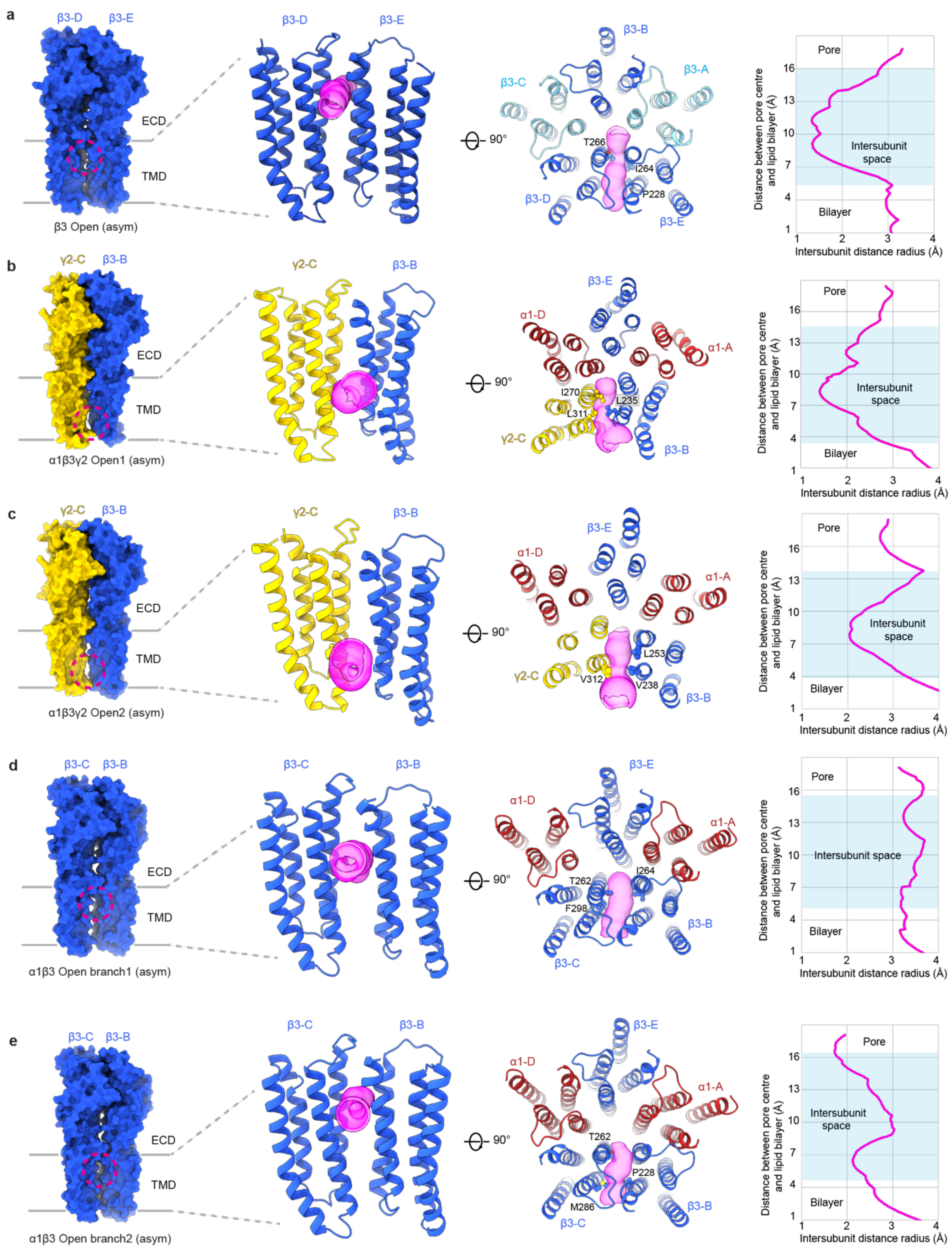

**Supplementary Figure 10 | The inter-subunit space of the GABA$_A$ receptor open states.** The panels show the surface representation of the interfaces with widest inter-subunit spaces for each open state of the β3 homomer, α1β3 diheteromer and α1β3γ2 triheteromer (left). A zoom in of the TMD region viewed parallel to the membrane showing the representation of the inter-subunit openings in magenta (middle-left). The TMD as viewed from the extracellular space showing the inter-subunit opening in magenta (middle-right). HOLE plot of the inter-subunit space dimensions (right). **a,** β3 homomer open state. **b,** α1β3γ2 open 1 state. **c,** α1β3γ2 open 2 state. **d,** α1β3 open state branch 1. **e,** α1β3 open state branch 2.

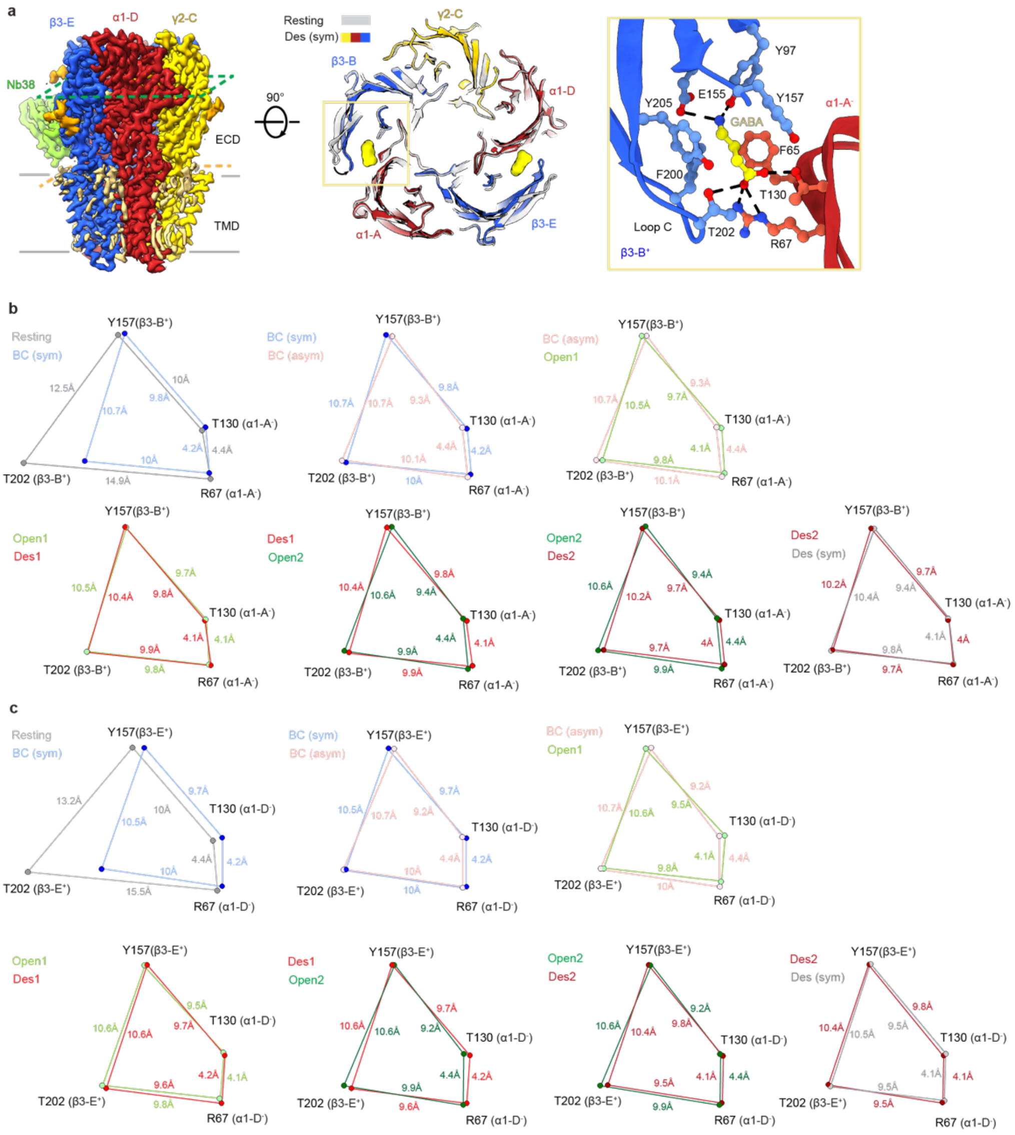

**Supplementary Figure 11 | Conformational changes in the neurotransmitter binding site during activation of the α1β3γ2 triheteromeric $GABA_A$ receptor.**

**a,** Cryo-EM map of the α1β3γ2 triheteromer viewed parallel to the membrane highlighting the plane of the GABA binding sites (left). View of the ECD cross-sectioned at the level of the GABA binding pocket showing the resting state in grey and the bound-closed symmetric state coloured (middle). One GABA-binding pocket viewed from the extracellular space. The amino acid side chains lining the binding site are shown as balls and sticks and GABA is coloured in yellow. Dashed lines indicate hydrogen bonds (right). **b,** Schematic diagram illustrating the changes in distances (Å) of key residues of the $β3\text{-}B^{+}/α1\text{-}A^{-}$ neurotransmitter binding site in the resting, bound-closed (symmetric), bound-closed (asymmetric), open 1, desensitised 1 (asymmetric), open 2, desensitised 2 (asymmetric) and long-lived desensitised states. **c,**

Schematic diagram illustrating the changes in distances (Å) of key residues of the β3-$E^+$/α1-$D^-$ neurotransmitter binding site in the resting, bound-closed (symmetric), bound-closed (asymmetric), open 1, desensitised 1 (asymmetric), open 2, desensitised 2 (asymmetric) and long-lived desensitised states.

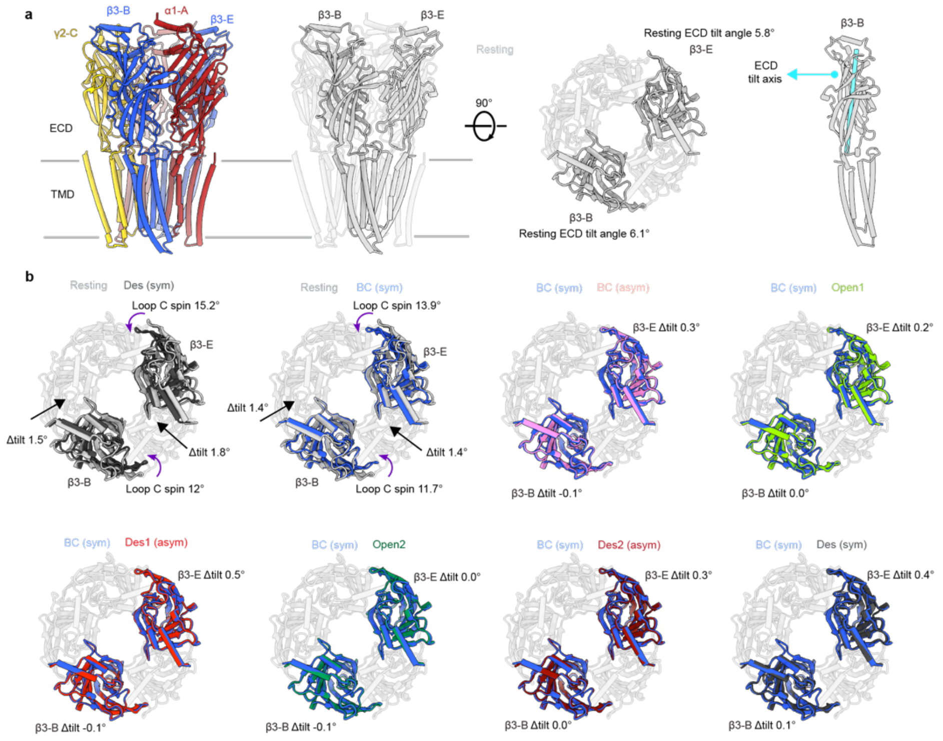

**Supplementary Figure 12 | Tertiary structural rearrangements in the extracellular domain during activation of the α1β3γ2 triheteromeric $GABA_A$ receptor.**

**a,** Side view of the α1β3γ2 triheteromer resting state model viewed parallel to the membrane plane (left). Subunits β3-B and β3-E highlighted in grey with the rest of the subunits transparent (middle). The β3-E is highlighted, and the tilt axis is shown in cyan (right). The tilt angle was calculated as in supplementary figure 13. **b,** The Δ tilt and loop C rotation calculations were performed as described in supplementary figure 13. The comparison is between the left and right state labels on top of each ECD as viewed from the extracellular space. No significant differences can be observed in the ECD tilt or loop C rotation for the agonist-bound states.

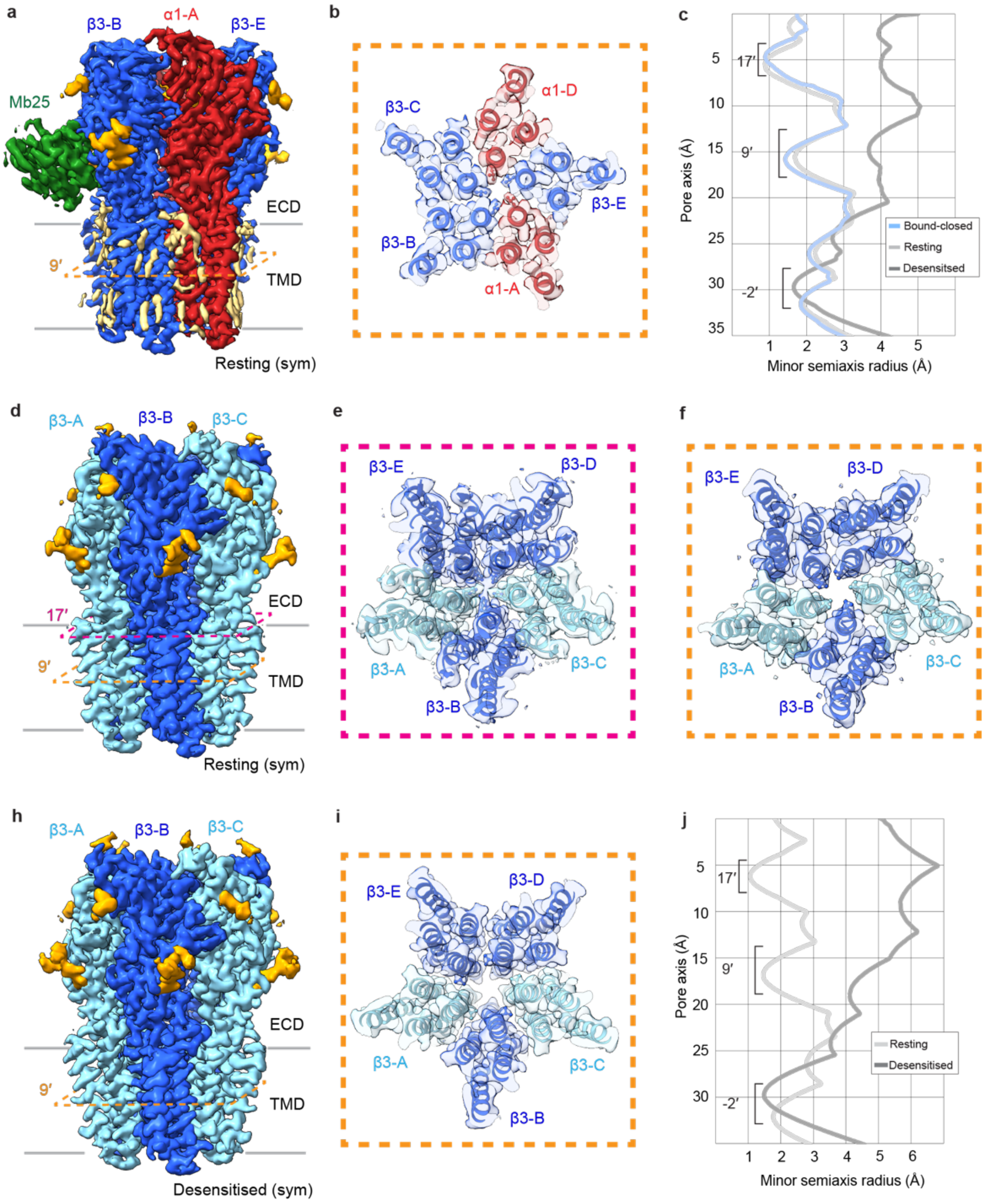

**Supplementary Figure 13 | Buffer spraying has no structural impact on the GABA$_A$ receptor.**

**a,** Side view of the cryo-EM map of the truncated α1β3 sprayed with PBS. The orange dotted line is positioned at the level of the 9′ activation gate. **b**, Cross-section of the 9′ activation gate as viewed from the extracellular space showing the closure of the 9′ Leu residues indicating the receptor is in the resting state. **c**, Pore profiles of α1β3 in the long-lived resting (sym), short-lived bound-closed (sym) and long-lived desensitised (sym) states. **d,** Side view of the cryo-EM map of the resting truncated β3 sprayed with PBS. The orange dotted line is positioned at the level of the 9′ activation gate and the magenta dotted line at the level of the 17′ gate. **e**,

Cross-section of the 17′ gate as viewed from the ECD showing the His267 17′ residues pointing towards the pore indicating the receptor is in the resting state. **f,** Cross-section of the 9′ activation gate as viewed from the extracellular space showing the closure of the 9′ Leu residues indicating the receptor is in the resting state. **h,** Side view of the cryo-EM map of the truncated long-lived desensitised β3 homomer sprayed with PBS. The orange dotted line is positioned at the level of the 9′ activation gate. **i,** Cross-section of the 9′ activation gate as viewed from the extracellular space showing the opening of the 9′ activation gate Leu residues and closure of the -2′ desensitisation gate Ala residues indicating the receptor is in the long-lived desensitised state. **j**, Pore profiles of β3 in the long-lived resting (sym) and long-lived desensitised (sym) states.

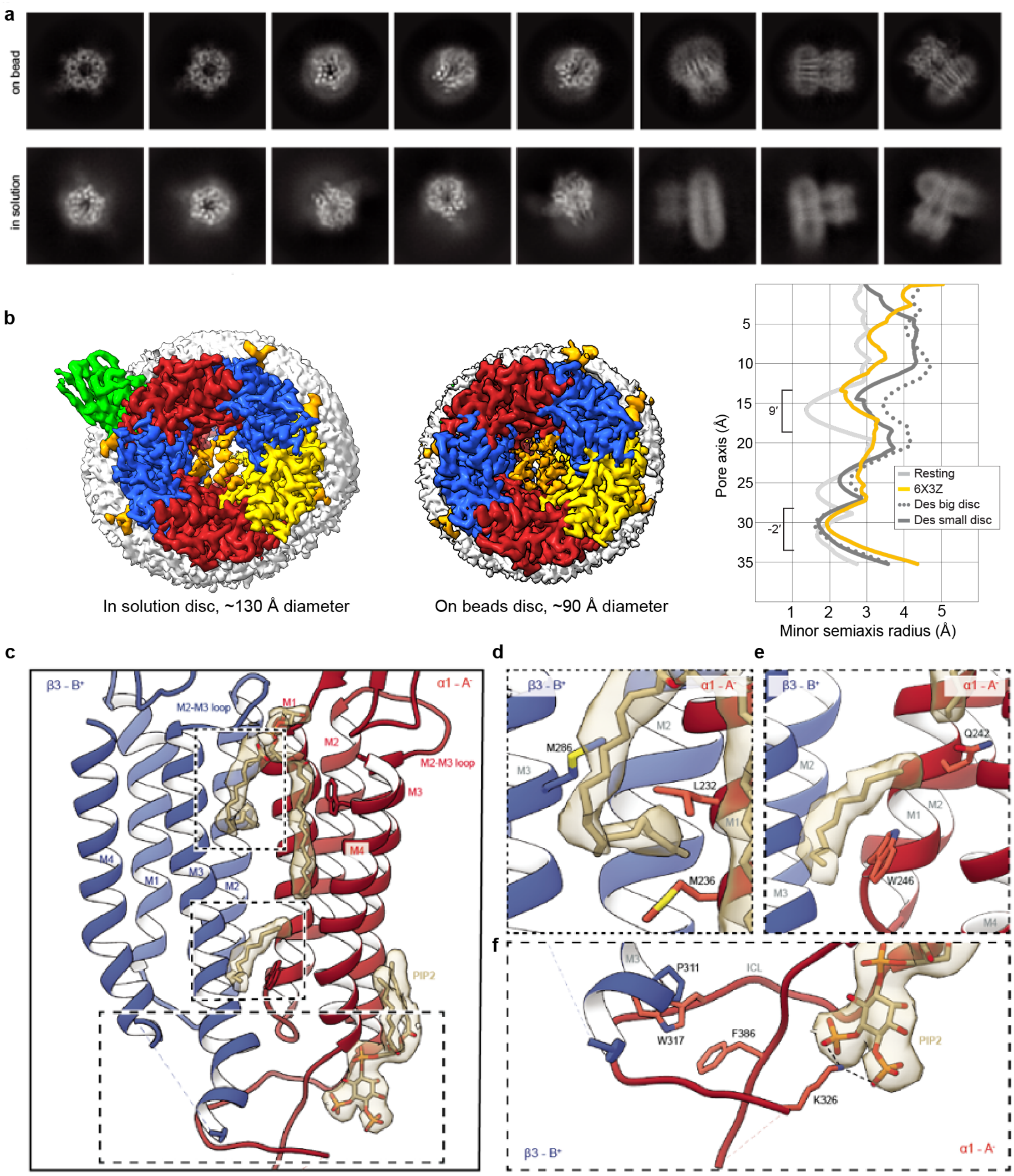

**Supplementary Figure 14 | The impact of nanodisc size on the GABA$_A$ receptor structure.** **a**, 2D classes of receptors reconstituted in MSP nanodiscs while still bound to the affinity resin, termed on beads (top) and receptors reconstituted in MSP nanodiscs in solution, after elution from the affinity resin (bottom). **b**, Top views of receptors including their nanodisc densities (white) reconstituted in solution (left) and on beads (middle). Pore profiles of various α1β3γ2L receptors compared to the GABA-bound α1β2γ2 receptor in Saposin A nanodisc (PDB: 6X3Z)[13] (right). **c**, Selected lipid interactions with the receptor at the $\beta3^+/\alpha1^-$ interface. Acyl chains are occupying the general anaesthetic binding site **(d)**, and the neurosteroids binding site **(e)**. **f**, The extended α1 intracellular loop interacts with $PIP_2$.

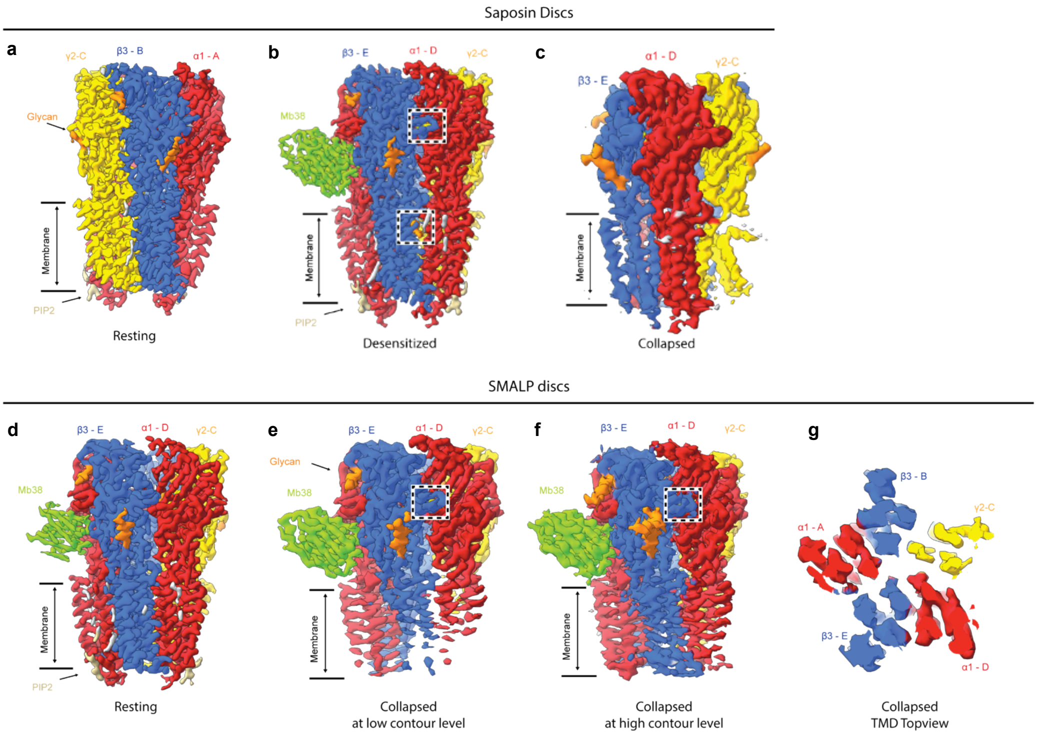

**Supplementary Figure 15 | The structural impact of nanodisc reconstitution on the full-length α1β3γ2L GABA$_A$ receptor.**

**a-c**, Cryo-EM maps of the saposin A disc reconstituted full-length α1β3γ2L GABA$_A$ receptor in the resting, desensitised (GABA+ETM) and collapsed state (+ GABA). **d-g**, Cryo-EM maps of the SMALP disc reconstituted full-length α1β3γ2L GABA$_A$ receptor in the resting and collapsed state. The collapsed state is represented at various volume levels (**e-f**). **g**, A slice through the TMD as seen from the extracellular space of the collapsed receptor in SMALPs.

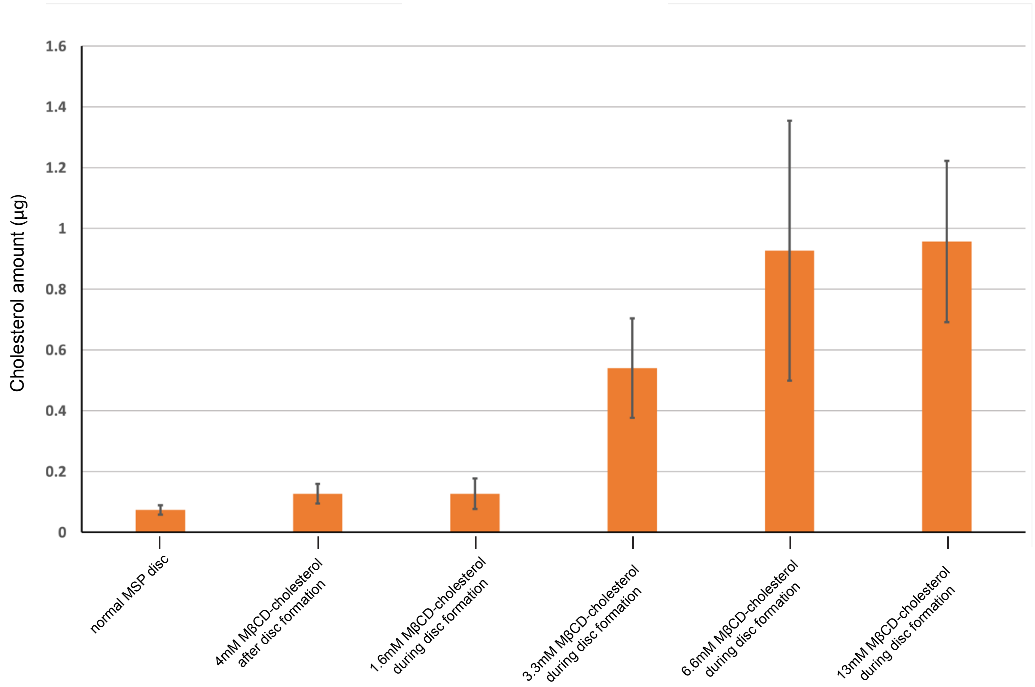

**Supplementary Figure 16 | Cholesterol content of the MSP2N2 disc upon MβCD-cholesterol loading.**

A plot showing the cholesterol content of MSP2N2 discs under different conditions. Cholesterol cannot be added effectively once the discs have already formed. However, if cholesterol enrichment is performed during disc assembly, the cholesterol content increases linearly with the concentration of MβCD–cholesterol used. Saturation is achieved roughly at 7mM MβCD-cholesterol. The measurements were performed in triplicates and represented as mean ± standard deviation.

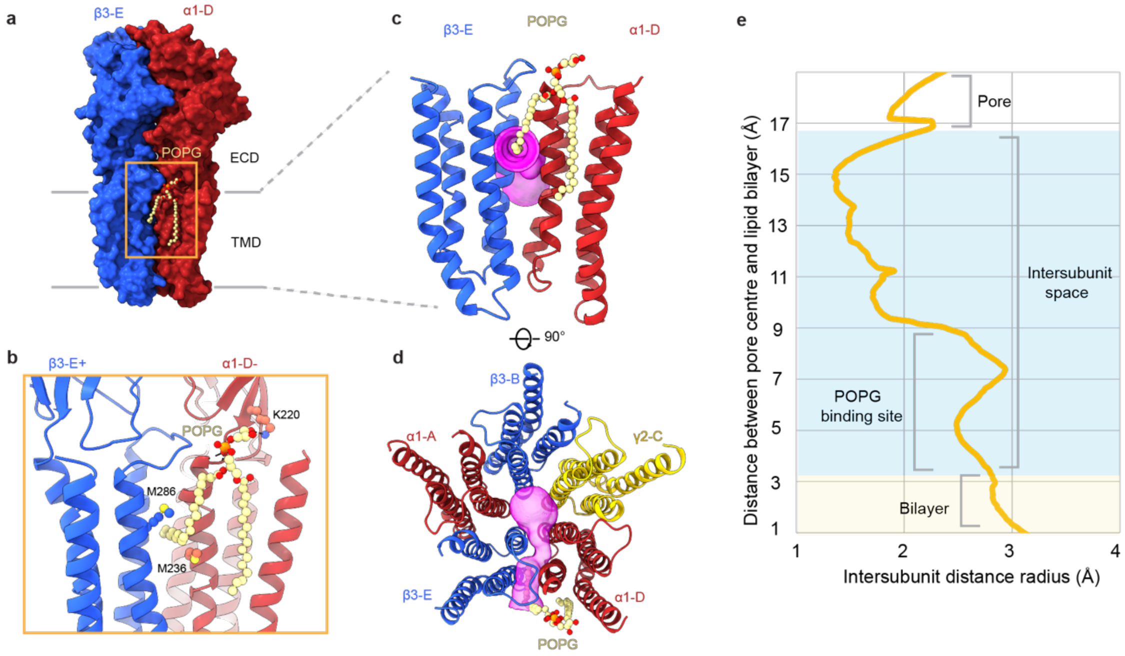

**Supplementary Figure 17 | The inter-subunit space of the α1β3γ2 symmetric long-lived desensitised state.**

**a,** Surface representation of the β3-E$^{+}$/ α1-D$^{-}$ interface in the long-lived desensitised state highlighting the binding site of POPG. **b,** Close up of the POPG (shown as balls and sticks; carbons coloured in khaki, oxygens in red, phosphates in orange) binding site. **c,** Close up of the transmembrane region showing the representation of the inter-subunit gap in magenta. **d,** The TMD as viewed from the extracellular space showing the inter-subunit gap in magenta. **e,** HOLE plot of the inter-subunit gap dimensions.

**Supplementary table 1 | Interface analysis and coupling**

The solvation energy and interface area were calculated for all time-resolved state of β3, α1β3 and α1β3γ2 by the PDBe PISA server (https://www.ebi.ac.uk/pdbe/pisa/)[48]. The models of the structures were split into ECDs and TMDs. The following residues forming the ECD and TMD region were included during the calculation – for α1 residues 1 to 222 – ECD, residues 223-418 – TMD; for β3 residues 1 to 218 – ECD, residues 219 to 447 – TMD; and γ2 residues 1 to 232 – ECD and residues 233 to 429 - TMD. The table reports interactions between neighbouring ECDs, neighbouring TMDs and intra-subunit ECD-TMD interactions.

**Supplementary Video 1 | Gating cycle of the human β3 $GABA_A$ receptor upon HSM application (<10ms)**

The video first shows the β3 $GABA_A$ receptor viewed parallel to the membrane. The receptor then rotates by 90° highlighting the ligand binding pockets of the resting state. The video then transitions between the resting state and the HSM-bound state of the ECD. Then, we focus on the transmembrane domain and show the transitions between the bound-closed (symmetric), bound-closed (asymmetric), open (asymmetric), desensitised (asymmetric) and desensitised (symmetric) states. The receptor is then rotated by 90° to become parallel to the membrane and the transitions are shown again.

**Supplementary Video 2 | Gating cycle of the human α1β3 $GABA_A$ receptor upon GABA application (<10ms, branch 1)**

The video first shows the α1β3 $GABA_A$ receptor viewed parallel to the membrane. The receptor then rotates by 90° highlighting the ligand binding pockets of the resting state. The video then transitions between the resting state and the GABA-bound state of the ECD. Then, we focus on the transmembrane domain and show the transitions between the resting (symmetric), bound-closed1 branch1 (asymmetric), bound-closed2 branch1 (asymmetric), open branch1 (asymmetric), desensitised branch1 (asymmetric) and desensitised (symmetric) states. The receptor is then rotated by 90° to become parallel to the membrane and the transitions are shown again.

**Supplementary Video 3 | Gating cycle of the human α1β3 $GABA_A$ receptor upon GABA application (<10ms, branch 2)**

The video first shows the α1β3 $GABA_A$ receptor viewed parallel to the membrane. The receptor then rotates by 90° highlighting the ligand binding pockets of the resting state. The video then transitions between the resting state and the GABA-bound state of the ECD. Then, we focus on the transmembrane domain and show the transitions between the resting (symmetric), bound-closed branch2 (asymmetric), open branch2 (asymmetric), desensitised branch2 (asymmetric) and desensitised (symmetric) states. The receptor is then rotated by 90° to become parallel to the membrane and the transitions are shown again.

**Supplementary Video 4 | Gating cycle of the human α1β3γ2 $GABA_A$ receptor upon GABA application (<10ms)**

The video first shows the α1β3γ2 $GABA_A$ receptor viewed parallel to the membrane. The receptor then rotates by 90° highlighting the ligand binding pockets of the resting state. The video then transitions between the resting state and the GABA-bound state of the ECD. Then, we focus on the transmembrane domain and show the transitions between the resting (symmetric), bound-closed1 (asymmetric), open1 (asymmetric), desensitised1 (asymmetric), open2 (asymmetric), desensitised2 (asymmetric) and desensitised (symmetric) states. The receptor is then rotated by 90° to become parallel to the membrane and the transitions are shown again.
